# Fruit flies of the subgenera *Drosophila*, *Siphlodora* and *Sophophora* use different mechanisms to adapt their daily activity to long photoperiods

**DOI:** 10.64898/2026.09.29.754759

**Authors:** Peter Deppisch, Francesca Sara Colizzi, Charlotte Helfrich-Förster

## Abstract

Over the course of evolution, many species have adapted to the environmental conditions found at higher latitudes. This is likely also true of the circadian system, which regulates daily behavior and helps measure day length to facilitate adaptation to the seasons, though how the circadian clock adapts to latitude remains largely unknown. Here, we investigated daily activity patterns under long photoperiods and in constant darkness, as well as the underlying circadian clock network, in 52 *Drosophila* species from around the world. We found a strong, significant positive correlation between the clock’s plasticity, measured as the phase difference between morning and evening activity, and the average geographical latitude from which the species originate. The mechanisms underpinning this increase in plasticity differed between species: in the subgenus *Siphlodora*, the clock weakened with increasing latitude, accompanied by a reduction of the neuropeptide Pigment-Dispersing Factor in the small ventrolateral clock neurons (s-LN_v_), whereas in the subgenus *Drosophila* the opposite was true. The subgenus *Sophophora*, by contrast, showed a negative correlation between circadian plasticity and PDF levels in the s-LN_v_ axons. These results suggest that *Drosophila* species have evolved diverse, independent strategies to adapt their circadian clock to higher latitudes.

## Introduction

Life on Earth has evolved under the influence of prominent daily and annual rhythmic changes in environmental light and temperature conditions. The extent of the annual rhythmic changes depends on the latitude. Near the equator, the length of the day (photoperiod) remains at around 12 hours throughout the year, whilst in regions closer to the poles there are drastic differences: very long days in summer, which can last for as long as continuous daylight, and very short days or complete darkness in winter.

To anticipate these daily and annual oscillations on Earth, virtually all organisms developed circadian and circannual clocks, which are thought to improve their fitness (Jabbur et al., 2024). Many organisms at higher latitudes measure the regular changes in photoperiod to adapt their physiology in advance for the coming winter and summer. Such photoperiodic responses include physiological, morphological, developmental, and reproductive adaptations (Goto, 2022). How organisms measure day length remains debated, but growing evidence points to the circadian clock as the underlying time reference, a hypothesis first proposed by Erwin Bünning (1936) and later refined and extended by Colin Pittendrigh (1972).

Many animals, from insects to vertebrates, exhibit bimodal activity rhythms with two active phases: one in the morning (M) and one in the evening (E) (Aschoff and Wever, 1965; Aschoff, 1966; Pittendrigh, 1972; Daan et al., 2001). Pittendrigh and Daan (1976) proposed that these activity phases are controlled by two separate circadian oscillators tracking dawn and dusk, respectively. This dual-oscillator system helps animals adapting to seasonal changes: on hot and long summer days, diurnal animals shift M activity into the late night by advancing its phase and E activity into the early night by delaying its phase, whereas in winter the two activity phases converge and occur during the warmer daytime. This flexible phase relationship by the oscillators not only helps animals coordinate activity with the best time of day but can also serve to measure day length and trigger photoperiodic responses (Pittendrigh (1972), reviewed by (Saunders, 2024), and Helfrich-Förster and Rieger (2024)).

Nevertheless, M and E oscillators are not completely independent of each other but are mutually coupled meaning that there is a limitation for expanding their phase relationship (ΔΨ_M-E_) under extremely long days. In fruit flies, this limitation appears to depend on species and latitude of origin (Rieger et al., 2012; Menegazzi et al., 2017; Beauchamp et al., 2018; Bertolini et al., 2019; Deppisch et al., 2022): with a few exceptions, high-latitude flies could extend ΔΨ_M-E_ further than low-latitude flies, suggesting they are capable of measuring the length of very long days.

Fruit flies have become key models for studying circadian clock adaptation to environmental cycles, since the molecular and neuronal mechanisms of the circadian clock are well-understood in *D. melanogaster* (reviewed in King and Sehgal, 2020) and are increasingly being uncovered in other Drosophilidae species as well (reviewed in Helfrich-Förster et al., 2020). Potential adaptive strategies for coping with long summer days include polymorphisms in circadian clock genes, differential splicing, and changes in the neuronal clock network in the brain (reviewed in Helfrich-Förster et al., 2020).

High-latitude species of the *Drosophila virilis* group show distinct differences in circadian clock anatomy compared to low-latitude species. Notably, they lack the blue-light photopigment Cryptochrome (CRY) in the large ventrolateral neurons (l-LN_v_)—a subset of their clock neurons—and show markedly reduced levels of neuropeptide Pigment-Dispersing Factor (PDF) in the small ventrolateral neurons (s-LN_v_), which serve as key pacemaker cells (Kauranen et al., 2012; Hermann et al., 2013; Menegazzi et al., 2017; Manoli et al., 2025) (Figure 1). Furthermore, these species show more PDF-positive fibers in the central, lateral and dorsal protocerebrum.

**Figure 1.**
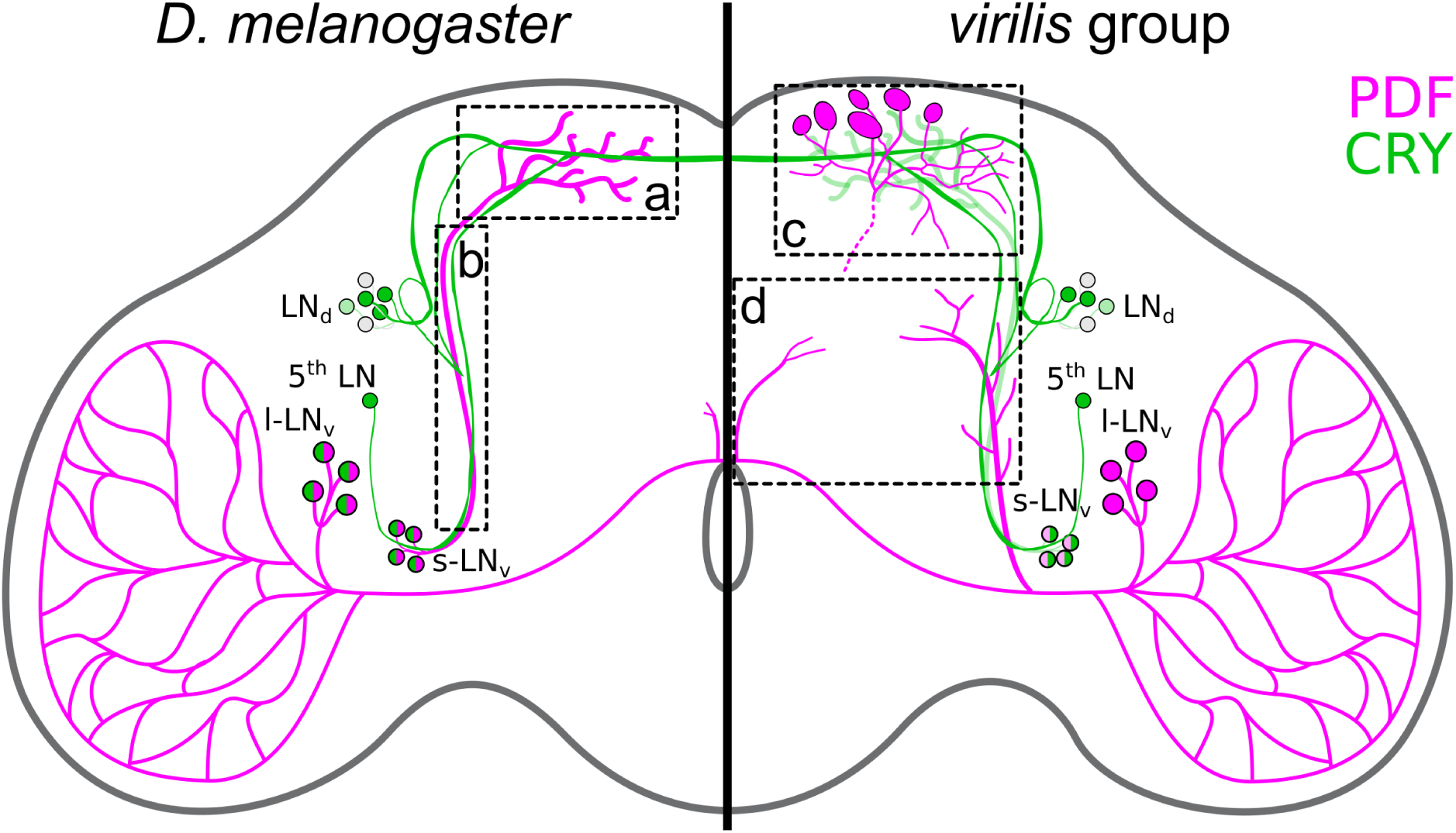
Simplified clock network in the brain of *Drosophila melanogaster* (left) and *Drosophila* species from the *virilis* group (right). Only a subset of clock neurons is shown: neurons associated with morning activity (small ventrolateral neurons, s-LN_v_) and evening activity (the fifth lateral neuron, 5^th^ LN, and dorsolateral neurons, LN_d_). PDF and CRY expression are shown in magenta and green, respectively. The *virilis* group species display markedly weaker PDF expression in the s-LN_v_, contain extra PDF-positive cells in the dorsolateral protocerebrum, extra PDF fibers in the central brain, and lack CRY expression in the large ventrolateral neurons (l-LN_v_). Stippled rectangles indicate the brain areas used to assess PDF staining across the different *Drosophila* species in this study: a, PDF staining intensity in the s-LN_v_ terminals; b, presence of PDF-positive varicosities along the s-LN_v_ axons; c, PDF-fiber density of non-clock neurons in the vicinity of the s-LN_v_ terminals; d, PDF-fiber density in the centrolateral brain.

Functionally, CRY and PDF play opposing roles in the timing of evening (E) activity. Light-activated CRY interacts with the clock protein Timeless (TIM) and promotes its degradation, which advances E activity (Kistenpfennig et al., 2018; Deppisch et al., 2022). PDF, in contrast, delays E activity by acting directly on the E neurons (Liang et al., 2016, 2017; Menegazzi et al., 2017; Schlichting et al., 2019; Vaze and Helfrich-Förster, 2021).

Consequently, flies with reduced or less effective CRY and enhanced PDF signaling to the E neurons can delay E activity until dusk even under very long photoperiods, such as 20 h light and 4 h darkness (LD 20:4). Reducing PDF levels in the s-LN_v_, the major pacemaker neurons, also weakens the circadian clock (Bahn et al., 2009; Hermann et al., 2013; Menegazzi et al., 2017). Weak clocks tend to be more plastic than strong clocks, adapting more easily to extreme photoperiods (Abraham et al., 2010; see also discussion in Bertolini et al., 2019; and Vaze et al., 2024). Recent findings further suggest that the *Pdf* gene may be a hotspot locus for the evolution of circadian plasticity, facilitating the spread of certain *Drosophila* species into new environments while limiting high-latitude colonization in others (Shahandeh et al., 2024).

We hypothesized that *D. melanogaster* possesses an ancestral circadian clock network that has been preserved in fly species that did not colonize high-latitude environments (Helfrich-Förster et al., 2020). This hypothesis is supported by observations in several phylogenetically distant species living in tropical areas, including *D. hydei*, *D. mercatorum*, *Zaprionus* sp., and *Bactrocera oleae*, which retain a CRY/PDF distribution pattern in their brains nearly identical to that of *D. melanogaster* and likewise exhibit robust circadian rhythms with low plasticity (Beauchamp et al., 2018; Bertolini et al., 2019). In contrast, species that successfully invaded far northern habitats, such as flies of the *virilis* group, evolved distinct modifications including severely reduced expression of PDF in the s-LN_v_, the emergence of additional PDF-positive fibers from the l-LN_v_ projecting into the protocerebrum, and the loss of CRY in the l-LN_v_ (Hermann et al., 2013; Menegazzi et al., 2017; Beauchamp et al., 2018). These modifications were accompanied by a weakening of their circadian clocks and at the same time with increased plasticity, necessary to adapt to long summer photoperiods.

However, there seem to exist multiple ways the circadian clock can adapt to high latitude habitats. For instance, *Chymomyza costata*, like the *Scaptodrosophila*, is positioned outside the main *Drosophila* radiation near the base of the Drosophilinae lineage (Finet et al., 2021; Tanaka et al., 2022), yet carries a clock network closely resembling that of *D. melanogaster* (Bertolini et al., 2019). Despite this, it has successfully colonized extreme northern regions such as Eastern Siberia, Northern Lapland, Iceland, and Northern Japan (Hackman et al., 1970). This suggests that circadian adaptation to high-latitude environments may be based on several distinct evolutionary solutions. Notably, most of the species studied to date belong to only a few groups, mainly the *virilis* and *repleta* groups within the subgenus *Siphlodora,* whereas *D. melanogaster* belongs to the subgenus *Sophophora* (Figure 2). Given that the family Drosophilidae comprises approximately 4,400 species (Finet et al., 2021), our current understanding is based on a very limited sample.

**Figure 2.**
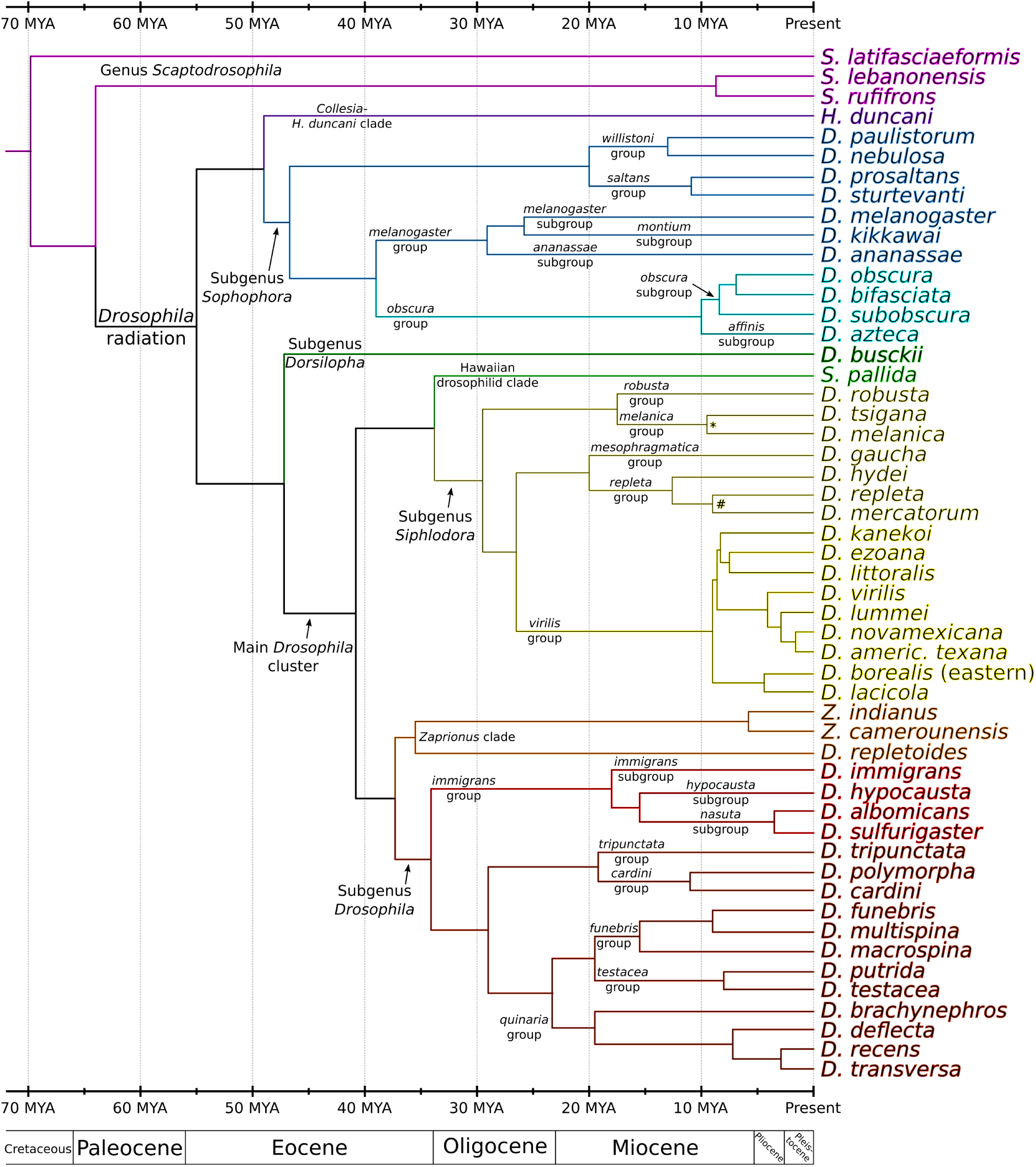
Phylogenetic relationship of the 52 fruit fly species used in this study. The divergence of the species is displayed as a chronogram with a timescale in million years ago (MYA). Relationships and divergence events were inferred from the studies of Morales-Hojas et al. (2011), Oliveira et al. (2012), Russo et al. (2013), Izumitani et al. (2016), Finet et al. (2021), Suvorov et al. (2022), and Tanaka et al. (2022). Note that the nodes are estimates and become less certain the further back in time one goes. The symbols * and # indicate divergence times that are controversial between different studies.

To address this gap, we investigated the circadian clock network and daily activity patterns under long photoperiods in 52 *Drosophila* species from around the world, covering large parts of the phylogeny. We found a strong and significant correlation between the phase difference in morning and evening activity ΔΨ_M-E_ and the latitude of the species’ place of origin. However, the PDF and CRY expression pattern in the clock network did not correlate with latitude across all *Drosophila* species, and not all high-latitude species had weak circadian clocks. These results suggest that *Drosophila* species have evolved multiple independent strategies to adapt their circadian clock to higher latitudes.

## Results and Discussion

### Phylogeny and origin of the selected *Drosophila* species

The Drosophilidae can be subdivided into the subfamilies Steganinae and Drosophilinae, the latter of which includes the genus *Drosophila* (Yassin, 2013). The radiation within the Drosophilinae, which began with the mass extinction at the Cretaceous–Paleogene, has produced over 4,000 currently described species (O’Grady and DeSalle, 2018; Finet et al., 2021), grouped into different subgenera and clades (Figure 2). The best-studied and most species-rich subgenera are *Sophophora*, *Siphlodora*, and *Drosophila*. Outside these groups is the genus *Scaptodrosophila*, which split from the genus *Drosophila* about 60 to 70 million years ago (Russo et al., 2013). Drosophilids are found on every continent except Antarctica and inhabit almost all climate zones, with most species being restricted to specific regions, whilst others are found worldwide.

To capture both the broad phylogenetic diversity within Drosophilinae and its varied geographic distribution, ranging from cosmopolitan species to those restricted to specific latitudes and climate zones, we selected 52 species from different subgenera, species groups, and continents for this study. The selected strains are easy to breed under standard *Drosophila* laboratory conditions and adequately represent the known phylogeny of fruit flies (Figure 2). Collection sites for the Drosophilidae species were obtained from the Taxodros website (https://www.taxodros.uzh.ch/), which compiles site reports from the relevant literature. Figure 3 shows the latitudinal distribution of recorded collection sites for each of the 52 species, grouped by phylogenetic relatedness. Figure 3—figure supplements 1 and Supplementary file 1 show the geographical distribution of each species and Figure 3—figure supplement 2 shows the variations in photoperiod as a function of latitude and season.

**Figure 3.**
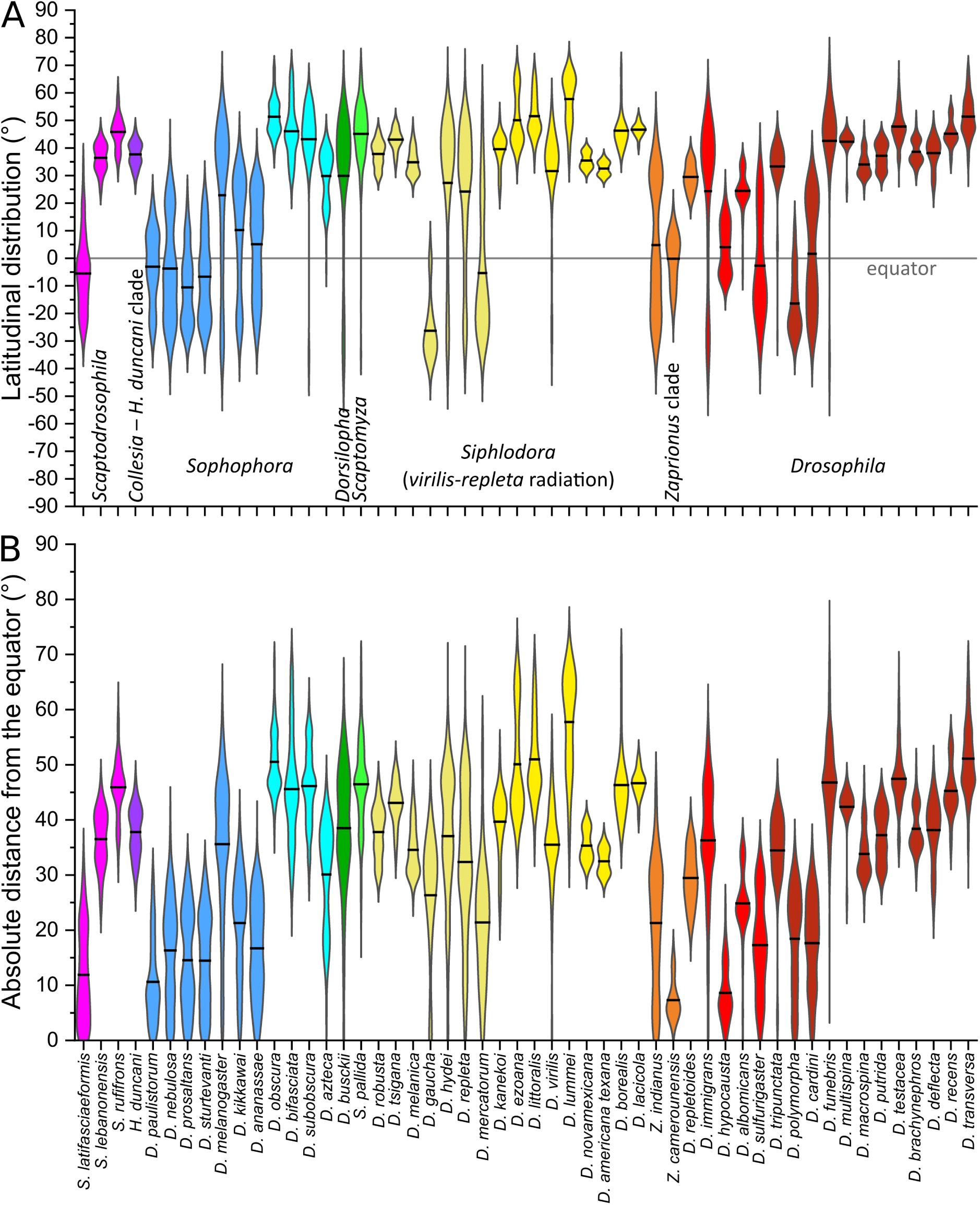
Latitudinal distribution pattern of the 52 *Drosophila* species investigated in this study. Reported collection sites were extracted from Gerhard Bächli’s Taxodros website. Species are arranged by phylogenetically related group, using the same color code as in Figure 2). **(A)** Violin plots of the global latitudinal distribution of all species with arithmetic means (thick lines). **(B)** Violin plots of the absolute distance of each species’ latitudinal coordinates from the equator. The online version of this article includes the following figure supplements for figure 3: **Figure 3—figure supplement 1.** Geographical distribution of the species (from TaxoDros). **Figure 3—figure supplement 2.** Variation of photoperiod in dependence of latitude and season. **Supplementary file 1**. Interactive maps of single Drosophila species’ distribution. **Supplementary file 2.** Descriptive statistics of geographical distribution records.

From the genus *Scaptodrosophila*, we studied three species: *S. latifasciaeformis*, a true equatorial species; *S. lebanonensis*, distributed mainly in arid, temperate regions of the Mediterranean and the Middle East; and *S. rufifrons*, restricted to the temperate Palearctic, with its northernmost records from southern Sweden and Siberia (Figure 3—figure supplements 1 and Supplementary file 1).

The remaining 49 species investigated belong to the *Drosophila* radiation, within which the subgenus *Sophophora*, together with the *Collesia* clade, separated about 50 million years ago from the subgenus *Dorsilopha* and the main *Drosophila* cluster. The latter comprises the subgenera *Siphlodora* and *Drosophila* as well as the two outgroup clades of Hawaiian *Drosophila* and *Zaprionus* species (Figure 2). Out of the *Collesia* clade, we studied only one species, *Hirtodrosophila duncani*, while we investigated 11 species of the *Sophophora* subgenus, including the model species, *D. melanogaster*. Of the subgenus *Dorsilopha*, we again studied only one species, *D. busckii*. We further studied one Hawaiian species, *Scaptomyza pallida*, and 3 species of the *Zaprionus* clade. This leaves 16 species each from the subgenera *Siphlodora* and *Drosophila* (Figure 2).

All major fly groups include species living close to the equator as well as others originating from high latitudes, making it possible to correlate behavioral and neurobiological traits with latitude of origin. For more detailed information on the species’ global distribution, see Figure 3—figure supplements 1 and Supplementary files 1 and 2.

### Daily activity pattern under 12h:12h light-dark conditions (LD12:12)

Under LD 12:12 with 1.5 hours of simulated dawn and dusk, most species showed bimodal activity patterns, with bursts of activity in the morning and evening, a siesta in between and very little activity at night, as is typical for *D. melanogaster* (e.g., Helfrich-Förster, 2000) (Figure 4A). In some cases, morning activity was rather low, as previously reported for species of the *virilis* group (Bahn et al., 2009; Beauchamp et al., 2018). A few species showed high nocturnal activity, a third small activity bout during midday, or completely lacked the siesta (Figure 4A).

**Figure 4.**
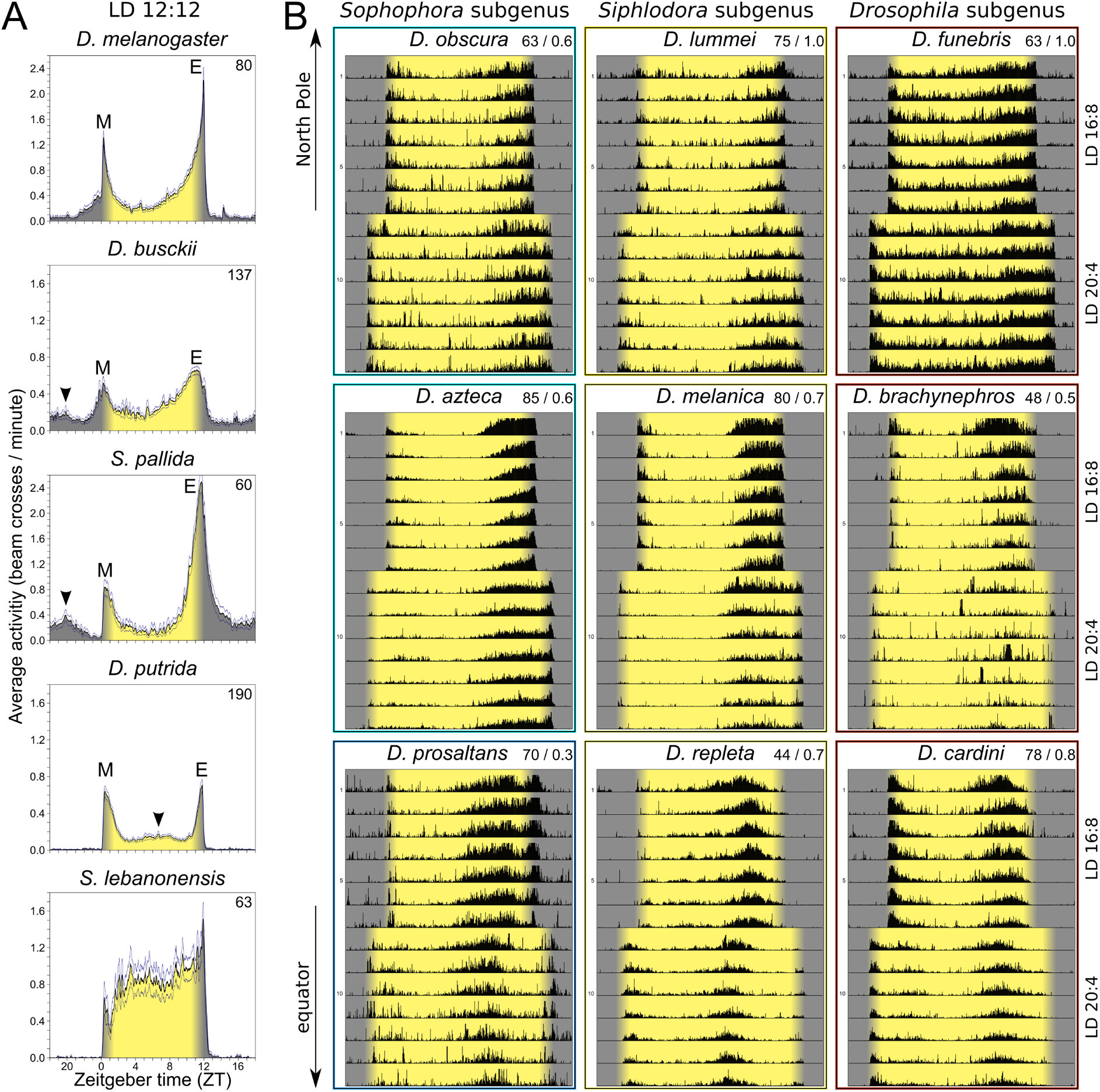
Average activity profiles under LD 12:12, and average actograms under LD 16:8 and 20:4. **(A)** Average activity profile of *D. melanogaster* and four species with slightly different activity patterns. Besides the typical morning (M) and evening (E) activity bouts, *D. busckii* and *S. pallida* showed nocturnal activity bouts, and *D. putrida* displayed midday activity (arrowheads). *S. lebanonensis* exhibited high activity throughout the day, lacking a siesta. The number of flies averaged for each profile is indicated in the top right corner. Thin lines above and below the mean indicate the standard error of the mean. **(B)** Average actograms of representative species from the three main *Drosophila* subgenera. The first row shows high latitude species, the second row species from temperate climate zones, and the bottom row species from low latitudes; only *D. repleta* is cosmopolitan and also occuring in temperate regions. The first seven days show recordings under LD 16:8, and the last seven days recordings under LD 20:4. The light phase is highlighted with a yellow background, the light-ramping (dawn/dusk simulation) is highlighted with a beige background, and the dark phase with a grey background. The frame colors refer to the phylogenetic color-coding from Figure 2. The number of flies and the upper activity threshold (lower for less active species and higher for highly active species) are given in the top right corner of each actogram. Each black line in the actograms represents an averaged 3-minute bin.

### Daily activity pattern under long photoperiods (LD16:8 and LD 20:4)

To test whether species can increase ΔΨ_M-E_ under long photoperiods, we recorded the flies for seven days in LD 16:8, followed by another seven days in LD 20:4. Figure 4B shows average actograms of nine representatives spanning different latitudes and the three main subgenera (*Sophophora*, *Siphlodora*, and *Drosophila*). These actograms demonstrate that species from high latitudes can shift their evening activity later into the day under long photoperiods than equatorial species can.

This pattern holds across all species and strains examined (Figure 5 under LD20:4, Figure 5—figure supplement 1 under LD16:8). Under LD 20:4, the M and E peaks of mostly low-latitude species failed to track dawn and dusk (some individuals of *D. kikkawai*, *D. kanekoi*, *D. sulfurigaster bilimbata*, and *D. deflecta* could not even synchronize to the cycle and instead free-ran at their endogenous period). By contrast, the M and E peaks of most high-latitude and some cosmopolitan species tracked dawn and dusk closely (e.g., *D. ezoana* and *D. funebris*). Among six *D. funebris* strains sampled from five different latitudes, the high-latitude strains from Alaska, Germany, and Hokkaido tracked dusk more closely than strains from lower latitudes (Figure 5, last row).

**Figure 5.**
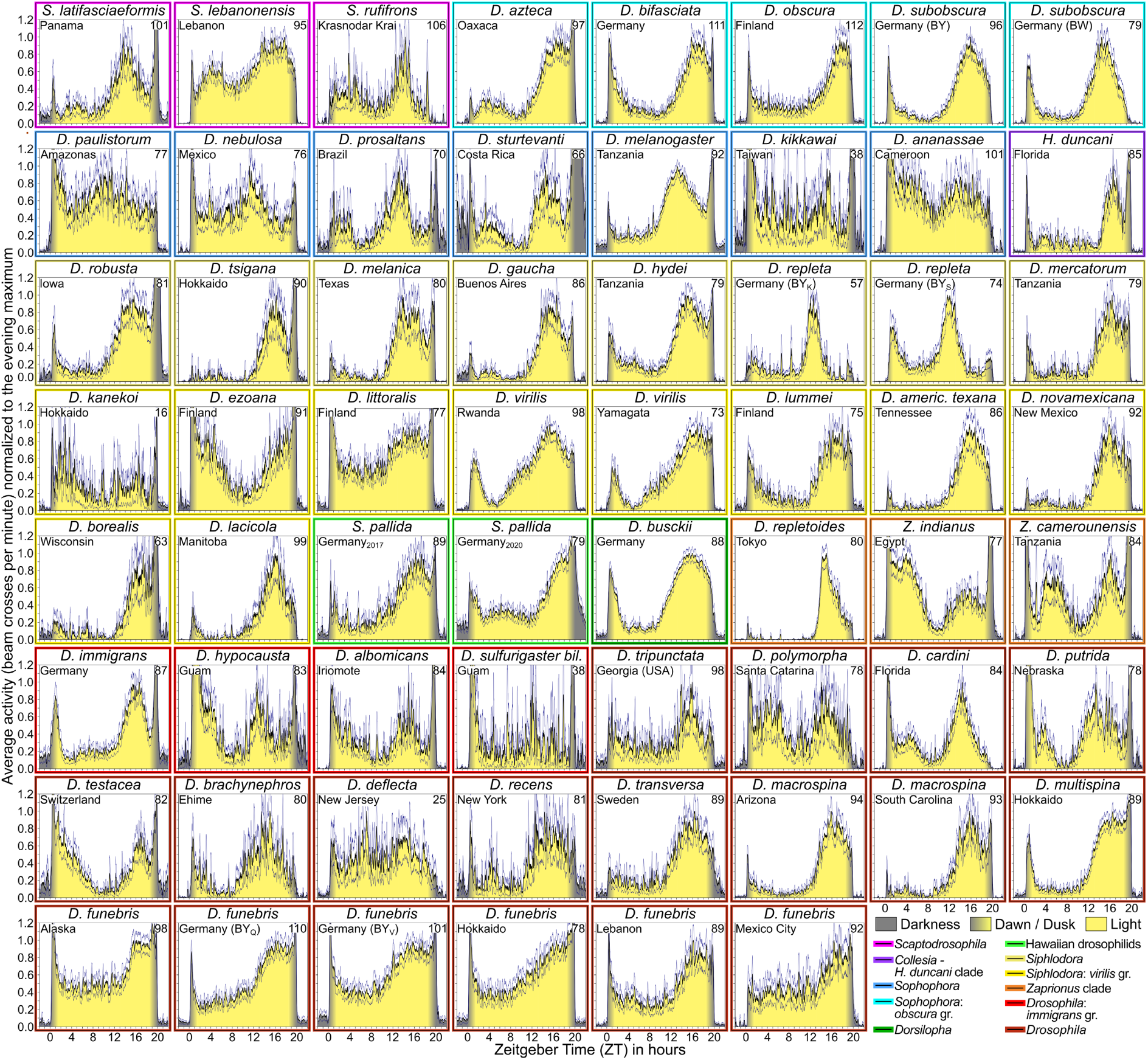
Average daily activity profiles of all tested species and strains in LD 20:4. Curves are normalized to the maximum amplitude of the circadian-driven morning or evening peaks. The origin of the species (or strain) is indicated in the upper left corner of each diagram, and the number of flies included is indicated in the upper right corner. Individuals that free-ran under LD 20:4 were omitted from the calculation of the daily activity profile (concerning the species *D. kikkawai*, *D. kanekoi*, *D. sulfurigaster bilimbata*, and *D. deflecta*). Frame colors refer to the phylogenic color-coding from Figure 2 (also shown in the lower right corner). During normalization, maximal activity during the evening peak was set to 1. This was done to improve visualization, since some species were very inactive. The magnitude of startle responses to dawn and dusk events was disregarded (e.g., the lights-off startle response was unusually high and long-lasting in *D. sturtevanti*, second row). The online version of this article includes the following figure supplements for figure 5: **Figure 5—figure supplement 1.** Average daily activity profiles of all tested species and strains in LD 16:8. **Figure 5—figure supplement 2.** Average activity profiles showing absolute activity values. **Supplementary file 3.** Determined phases of M and E peaks and calculated ΔΨ_M-E_ values for all species and all tested photoperiods.

Nevertheless, there are exceptions to this general pattern. For example, *S. rufifrons* from the temperate climate zone could not track dusk, showing an activity profile similar to its tropical relative *S. latifasciaeformis* (Figure 5, first row). Interestingly, *S. lebanonensis*, which was unimodal under LD12:12, became bimodal under LD 20:4, indicating that its activity is controlled by M and E oscillators that overlapped in LD 12:12 rather than being absent (Figure 4A average day at the bottom and Figure 5, first row). The remaining species, whether from outgroups or from groups represented by only a single species in our sample, fit well into the general pattern of M and E peak timing.

In addition to M and E activity, several species displayed sudden increases in their activity during simulated twilight, resulting in sharp peaks at both dawn and dusk (e.g., *D. sturtevanti*, Figure 5, row 2; *Z. indianus* and *Z. camerounensis*, Figure 5, row 5). Other species showed this only at dusk (e.g., *D. robusta* and *D. tsigana*, Figure 5, row 3), or only at dawn (e.g., *D. cardini*, Figure 5, row 6). These dawn and dusk peaks closely resemble the lights-on and lights-off peaks observed under abrupt (rectangular) light-dark cycles, which are known startle responses to the sudden changes in lighting (Rieger et al., 2012; Schlichting et al., 2019). We assume the peaks observed here reflect the same type of startle response, since our fixed 1.5-hour twilight transition, chosen as a compromise across the full latitudinal range of species tested, is unnaturally short compared to the natural twilight duration experienced by high-latitude species in summer. We therefore excluded these dawn and dusk peaks from further analysis and instead focused on the underlying circadian-driven M and E peaks.

### ΔΨ_M-E_ correlates significantly with latitude

As done previously (Beauchamp et al., 2018; Deppisch et al., 2022), we determined the peak phases of morning and evening activity and calculated the phase difference ΔΨ_M-E_ between them under LD 20:4, to assess the ability of each species to adapt to long photoperiods. The determined phases of M and E peaks as well as the calculated ΔΨ_M-E_ values for all species except *D. deflecta* are listed in Supplementary file 3. For *D. deflecta*, we were unable to calculate peak phases, since its activity pattern was too irregular for clear M and E peaks to be discernible (Figure 5, row 7). We therefore used n = 51 for all analyses involving ΔΨ_M-E_. Spearman correlation analysis revealed a highly significant correlation between ΔΨ_M-E_ and the mean absolute distance of each species’ distribution from the equator (Figure 6A; Spearman’s rank correlation coefficient ρ_(51)_ = 0.770, p < 0.001). A similar correlation emerged when ΔΨ_M-E_ was plotted against the polarmost (most poleward) latitudinal collection site of each species (Figure 6—figure supplements 1 and 2) as well as against the original latitude of the tested strains (Figure 6—figure supplement 3). For the latter analysis we treated each strain of a given species separately (e.g., four strains of *D. funebris*, and two strains of *S. pallida*, *D. virilis*, *D. repleta* and *D. macrospina*). Among the four *D. funebris* strains, those from higher latitudes showed a larger ΔΨ_M-E_ than those from lower latitudes, suggesting that ΔΨ_M-E_ increases with latitude even within a single species. This is consistent with previous observations in *D. littoralis* and *D. montana* (Manoli et al., 2025) and observations in *D. melanogaster* (Deppisch et al., 2022).

**Figure 6.**
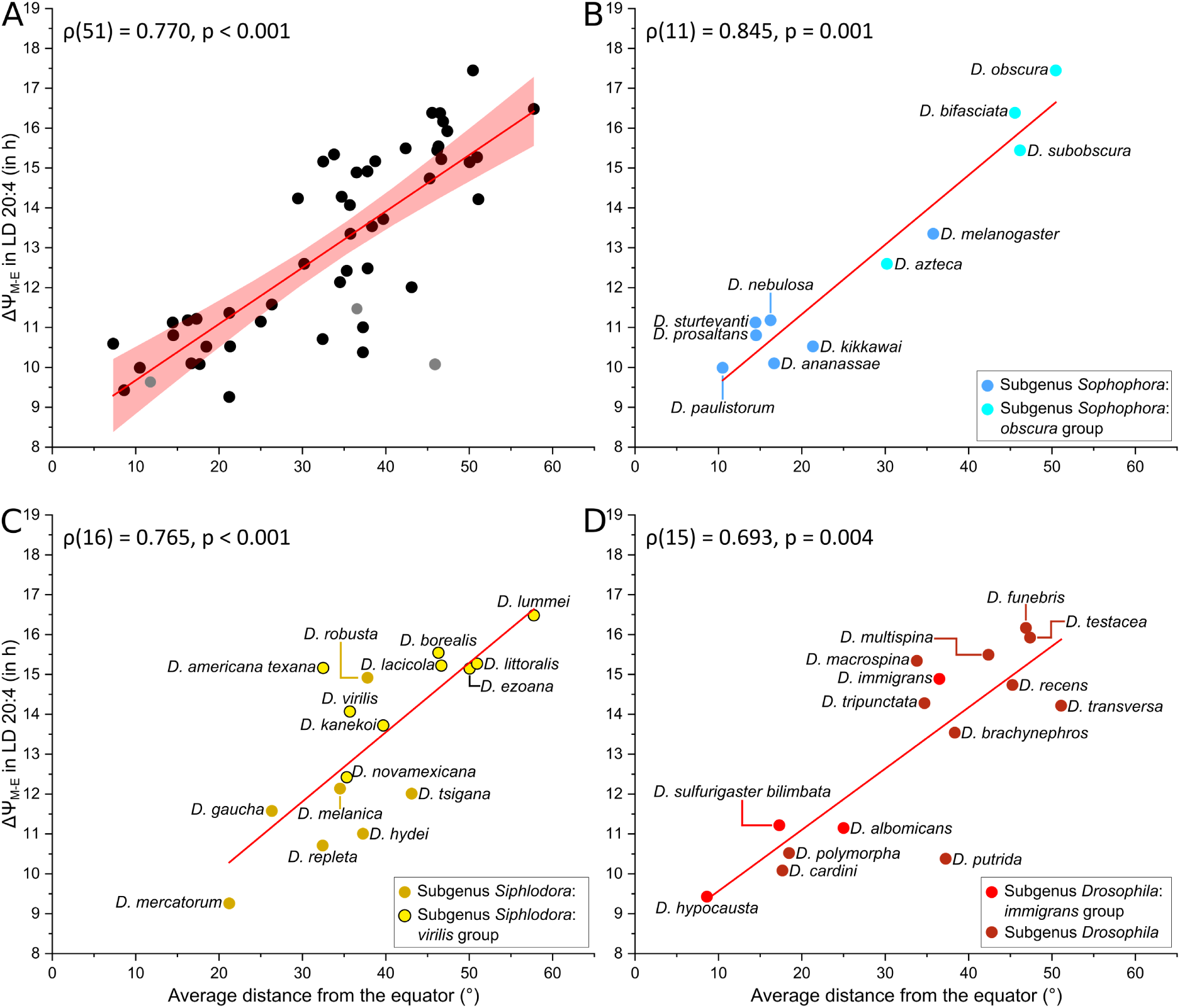
Correlation between ΔΨ_M-E_ in LD 20:4 and average latitudinal distance from the equator in all *Drosophila* species **(A)** and the three different *Drosophila* subgenera **(B-D)** (trimmed by 3 % to exclude potential outlier reports, i.e., false extreme latitudinal values). To avoid distorting the correlation coefficient with duplicate values for species with multiple tested strains, only one strain per species is included (51 species total): *D. subobscura* from Bavaria, *S. pallida* collected in 2020, *D. repleta* from Bavaria (strain S), *D. virilis* from Rwanda, *D. funebris* from Alaska, and *D. macrospina* from Arizona. The three *Scaptodrosophila* species, which fall outside the main *Drosophila* radiation (see Figure 2), are shown in grey. **(B)** *Sophophora* subgenus, **(C)** *Drosophila* subgenus, **(D)** *Siphlodora* subgenus; Regression lines are indicated in red. The 95 % confidence band is illustrated in **(A)**. Spearman’s rank correlation coefficients are indicated in the upper left corner of each panel. The online version of this article includes the following figure supplements for figure 6: **Figure 6—figure supplement 1.** Correlation between ΔΨ_M-E_ and most poleward report of all *Drosophila* species. **Figure 6—figure supplement 2.** Correlation between ΔΨ_M-E_ and latitudinal distribution of all *Drosophila* species. **Figure 6—figure supplement 3.** Correlation between ΔΨ_M-E_ in and the original latitude of all tested strains.

A separate evaluation of the three main subgenera of the *Drosophila* radiation (*Sophophora*, *Siphlodora*, and *Drosophila*), revealed similarly strong, or even stronger correlations, than for the species as a whole population (Figure 6B-D; Figure 6—figure supplements 1 and 2). The relationship was robust even within the *virilis* group of the *Siphlodora*, whose members are predominantly distributed in temperate and boreal regions. Together, these results indicate that a species’ ability to extend the morning-evening phase relationship increases with its average distributional distance from the equator, consistent with our earlier observations in selected species of the *Siphlodora* and *Sophophora* subgenera (Rieger et al., 2012; Menegazzi et al., 2017; Beauchamp et al., 2018; Bertolini et al., 2019; Deppisch et al., 2022).

### CRY distribution in the l-LN_v_ correlates with ΔΨ_M-E_ and latitude in the *Siphlodora* subgenus

To test whether CRY expression in the l-LN_v_ might contribute to the increase in ΔΨ_M-E_ with latitude, we assessed the presence of CRY in the l-LN_v_ across all 52 species and correlated it with both ΔΨ_M-E_ and latitude. CRY expression varied considerably among the different *Drosophila* species. Some species appeared to completely lack CRY in the l-LN_v_, others showed only very weak anti-CRY staining in these neurons, and others showed strong CRY staining, whereas CRY staining in the other clock neurons remained unchanged in all species (Figure 7A shows two examples, Figure 7—figure supplement 1 illustrates additional species). To quantify this variation, we scored each species on a scale from −3 (no staining) to 0 (staining like in *D. melanogaster*), reflecting the degree of CRY loss in the l-LN_v_.

**Figure 7.**
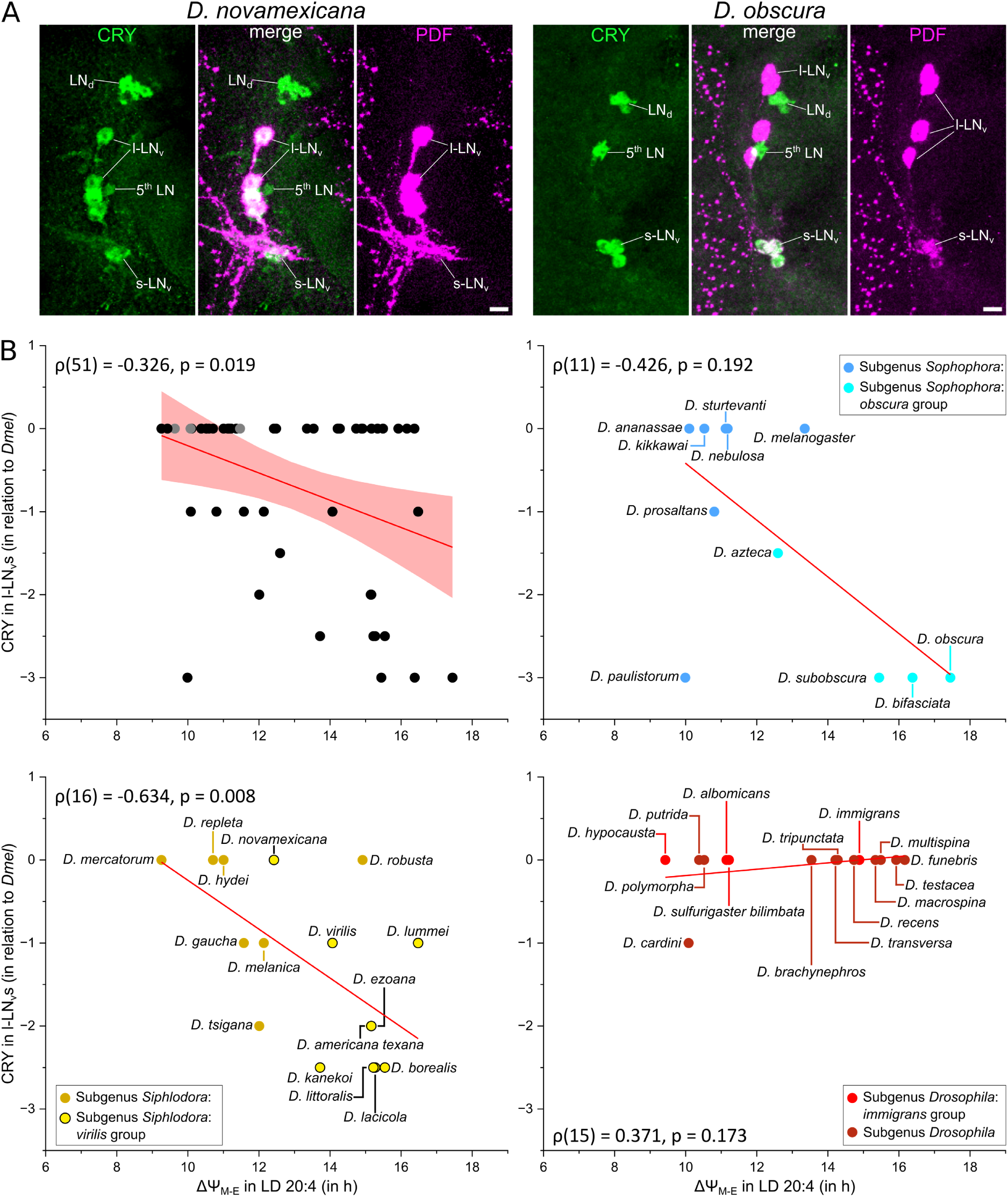
CRY expression in the l-LN_v_ of two *Drosophila* species and correlation of CRY expression with ΔΨ_M-E_ across species. **(A)** Confocal images of the lateral clock neurons of *D. novamexicana* (subgenus *Siphlodora) and D. obscura* (subgenus *Sophophora*), immunostained with anti-CRY (green) and anti-PDF (magenta); middle panels show an overlay of both channels. The dorsolateral neurons (LN_d_) and the fifth lateral neuron (5^th^ LN) belong to the evening neurons, while the small ventrolateral neurons (s-LN_v_) are morning neurons. Note that *D. novamexicana* retains CRY in the four l-LN_v_, while *D. obscura* lacks CRY in these neurons. Scale bar: 10 μm. **(B)** Correlation between CRY presence in the l-LN_v_s and average ΔΨ_M-E_ in all *Drosophila* species and within three main subgenera. The same 51 strains as in Figure 6 are included here. Labeling as in Figure 6. The online version of this article includes the following figure supplements for figure 7: **Figure 7—figure supplement 1.** CRY staining in different species. **Figure 7—figure supplement 2.** Correlation between CRY expression and latitudinal distribution.

CRY staining in the l-LN_v_ showed a weak correlation with both ΔΨ_M-E_ (Figure 7B) and latitude (Figure 7—figure supplement 2) across the full dataset. However, subgenus-level analysis showed that these correlations were significant only within *Siphlodora*, where they were stronger than across the full species population, but not within *Sophophora* or *Drosophila* subgenera (Figure 7B; Figure 7—figure supplement 2). Within *Sophophora*, CRY in the l-LN_v_ tended to decrease with increasing latitude and ΔΨ_M-E_, whereas within *Drosophila* the opposite trend appeared, with CRY tending to increase with both variables. In other words, only in the *Siphlodora* subgenus, CRY in the l-LN_v_ decreased significantly with increasing latitude and this effect was associated with a corresponding increase in ΔΨ_M-E_.

### Density of PDF fibers and PDF staining intensity in different brain areas correlate differently with ΔΨ_M-E_ and latitude

As shown in Figure 1, expression of the neuropeptide PDF varies throughout specific cells and brain areas among previously investigated fly species, and this variability appears to be responsible for differences in activity rhythms in the *virilis* species group (Menegazzi et al., 2017). However, PDF distribution remains largely uncharacterized in most species of the *Sophophora* and *Drosophila* subgenera and their outgroups (Figure 2). Here, we stained the brains of all 52 species with anti-PDF (or anti-PDH) to portray similarities and differences in PDF distribution. We found that the PDF expression pattern depended, to some extent, on the phylogenetic origin of the species.

In *Sophophora*, PDF was found exclusively in the s-LN_v_ and l-LN_v_ (Figure 8, upper row); Some PDF Tri-neurons in the esophageal foramen were occasionally PDF-positive only in young flies as was found previously (Helfrich-Förster, 1997; Nagy, 2017; Selcho et al., 2018). In several species of the subgenus *Siphlodora*, PDF staining in the s-LN_v_ was very low or completely absent, and in most *Siphlodora* species, PDF was additionally detected in non-clock neurons identified as the corazonin-positive neurons of the *pars lateralis* (Figure 8, rows 2 and 3, cyan arrows) (Hamanaka et al., 2007; Manoli et al., 2023). These corazonin-positive neurons branched extensively in the lateral and dorsal protocerebrum (Figure 8, cyan arrowheads), coming into close proximity with fibers from both the evening neurons and the s-LN_v_ terminals. This suggests that PDF from the corazonin neurons could signal to the evening neurons and delay evening activity. In addition, constant PDF release from these non-clock neurons might also disturb the rhythmic signaling of the s-LN_v_, weakening overall clock output.

**Figure 8.**
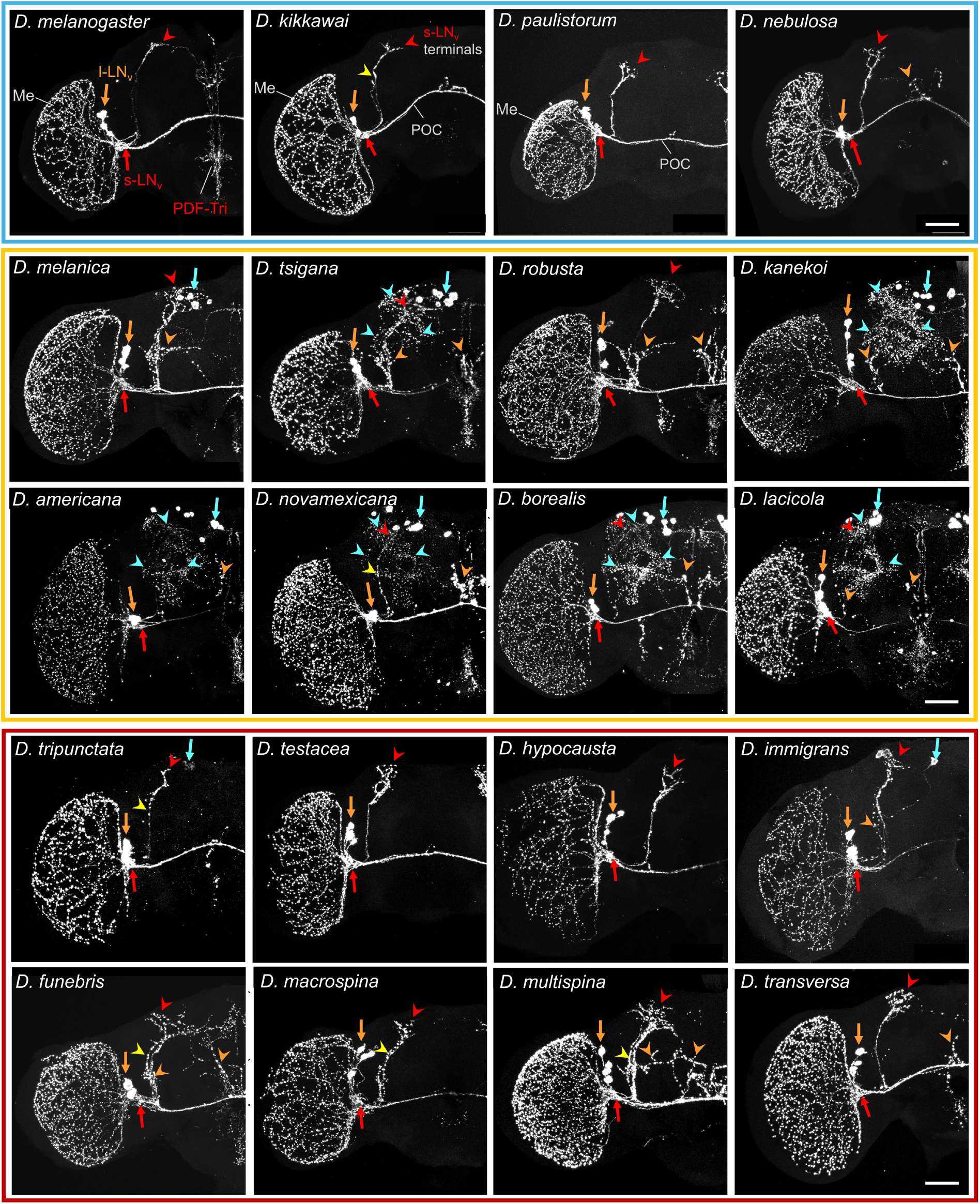
PDF expression pattern in the left brain hemisphere of selected species of the *Sophophora* (blue frame), *Siphlodora* (yellow frame) and *Drosophila* (red frame) subgenera. Orange arrows point to the l-LN_v_; orange arrowheads indicate l-LN_v_ fibers leaving the posterior optic commissure (POC) and projecting into the centrolateral brain. Red arrows mark the s-LN_v_; red arrowheads mark their dorsal terminals. Yellow arrowheads indicate varicose projections of the s-LN_v_ toward the dorsal brain. Cyan arrows mark putative PDF-positive corazonin cell clusters; cyan arrowheads mark their projections in the brain. Scale bar: 50 μm.

In the subgenus *Drosophila*, PDF was occasionally present in *pars lateralis* neurons that may correspond to the corazonin-positive neurons (Figure 8, row 4, cyan arrows), though labeling was weak and fibers arising from these neurons were rarely visible. In this subgenus, dorsally projecting s-LN_v_ axons often contained numerous varicosities (Figure 8, rows 4 and 5, yellow arrows), suggesting PDF release into the lateral protocerebrum and signal transmission to the evening neurons. The s-LN_v_ terminals were also more strongly stained with the PDF-antibody, with more arbors and varicosities than in the other two subgenera (Figure 8, rows 4 and 5, red arrowheads).

In some species across all three subgenera, varicose fibers from the l-LN_v_ left the posterior optic commissure and projected into the centrolateral brain (Figure 8, orange arrowheads). These extra l-LN_v_ projections were most pronounced in the *Siphlodora* subgenus and were sometimes difficult to distinguish from fibers of the corazonin-positive non-clock neurons.

To test whether any of these PDF staining characteristics correlate with ΔΨ_M-E_ and latitude, we quantified PDF density and staining intensity in the four regions indicated in Figure 1: the centrolateral brain (area d), in the dorsal protocerebrum (area c), the varicosities along s-LN_v_ axons running to the dorsolateral brain (area b), and the s-LN_v_ terminals (area a). Quantification was performed relative to PDF-staining in *D. melanogaster*, using a scale from −3 to 3 for each brain region (−3 = much less PDF staining or fewer PDF fibers than in *D. melanogaster*, 0 = no difference from *D. melanogaster*, 3 = much more PDF staining or more PDF fibers than in *D. melanogaster*).

Spearman correlation analysis revealed a moderate but significant correlation between PDF-fibers in the centrolateral brain and both ΔΨ_M-E_ (Figure 9A) and latitude (Figure 9—figure supplement 1A) across the whole species population. Subgenus-level evaluation showed an even stronger correlation within the *Sophophora* subgenus than across the full dataset (Figure 9B; Figure 9—figure supplement 1B). This correlation was also evident within *Siphlodora* (Figure 9C; Figure 9—figure supplement 1C), but not in the *Drosophila* subgenus (Figure 9D; Figure 9—figure supplement 1D). Together, these results suggest that PDF released from the fibers in the centrolateral brain signals to the E neurons and delays evening activity, producing a larger ΔΨ_M-E_ under long photoperiods in species of the *Sophophora* and *Siphlodora* subgenera.

**Figure 9.**
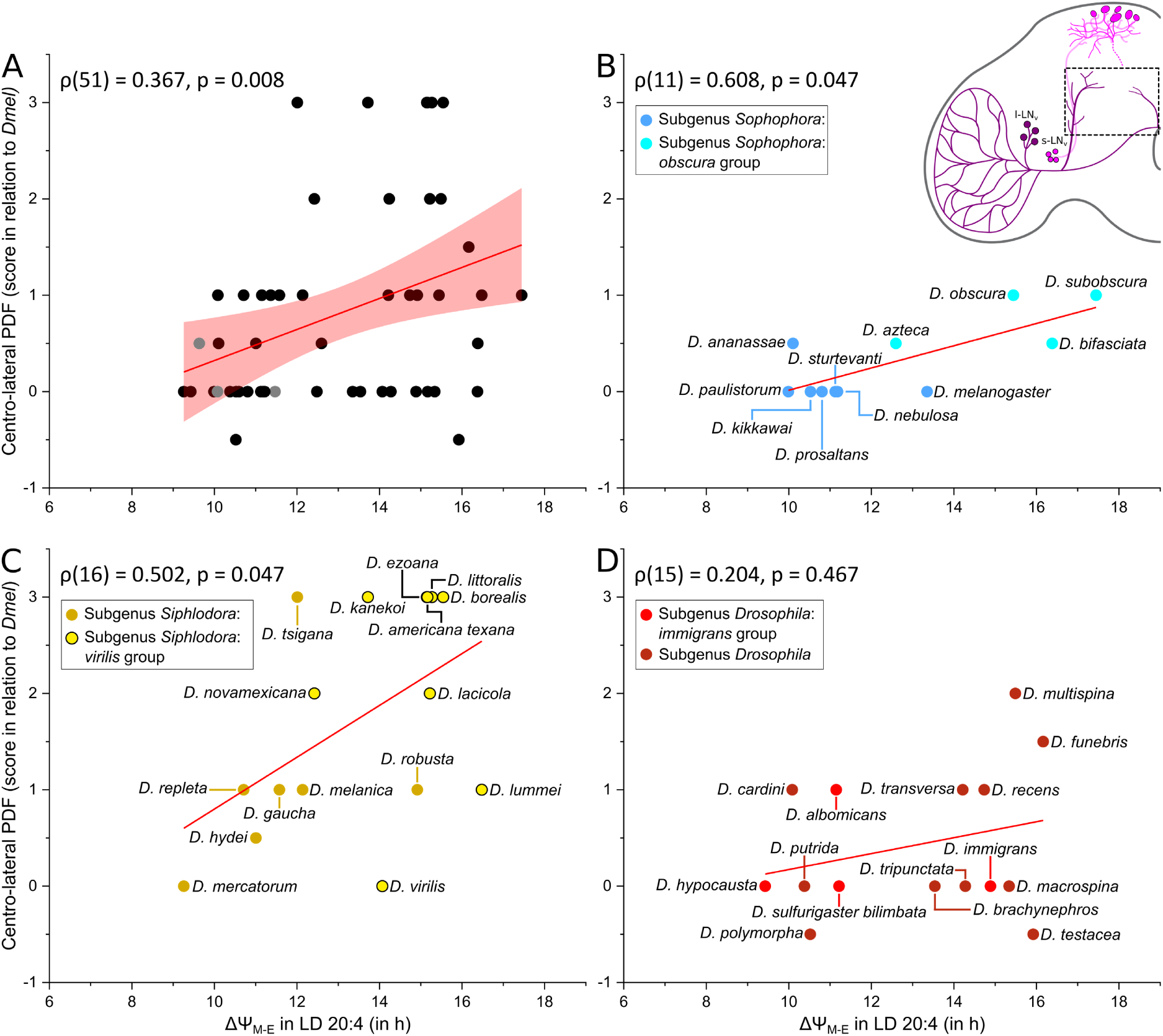
Correlations between the relative abundance of PDF in the centrolateral brain area and ΔΨ_M-E_ in all *Drosophila* species **(A)** and within the three different *Drosophila* subgenera **(B-D**). The inset in the upper right corner indicates the quantified brain region. The same 51 strains as in Figure 6 are included here. **(B)** *Sophophora* subgenus, **(C)** *Drosophila* subgenus, **(D)** *Siphlodora* subgenus. Labeling as in Figures 6 and 7. The online version of this article includes the following figure supplement for figure 9: **Figure 9—figure supplement 1.** Correlations between the relative abundance of PDF in the centrolateral brain area and latitude in all *Drosophila* species.

The presence of PDF in the dorsal brain near the s-LN_v_ terminals and in the ascending s-LN_v_ axons did not correlate with ΔΨ_M-E_ or latitude across the whole species population. Within the *Siphlodora* subgenus, however, PDF presence from the corazonin-positive neurons close to the s-LN_v_ terminals correlated strongly with both measures: greater PDF fiber density in this area was positively correlated with ΔΨ_M-E_ (Figure 10A) and latitude (Figure 10A—figure supplement 1). In addition, within the *Sophophora* subgenus, PDF in the ascending s-LN_v_ axons was negatively correlated with ΔΨ_M-E_ (Figure 10B) and latitude (Figure 10B—figure supplement 1).

**Figure 10.**
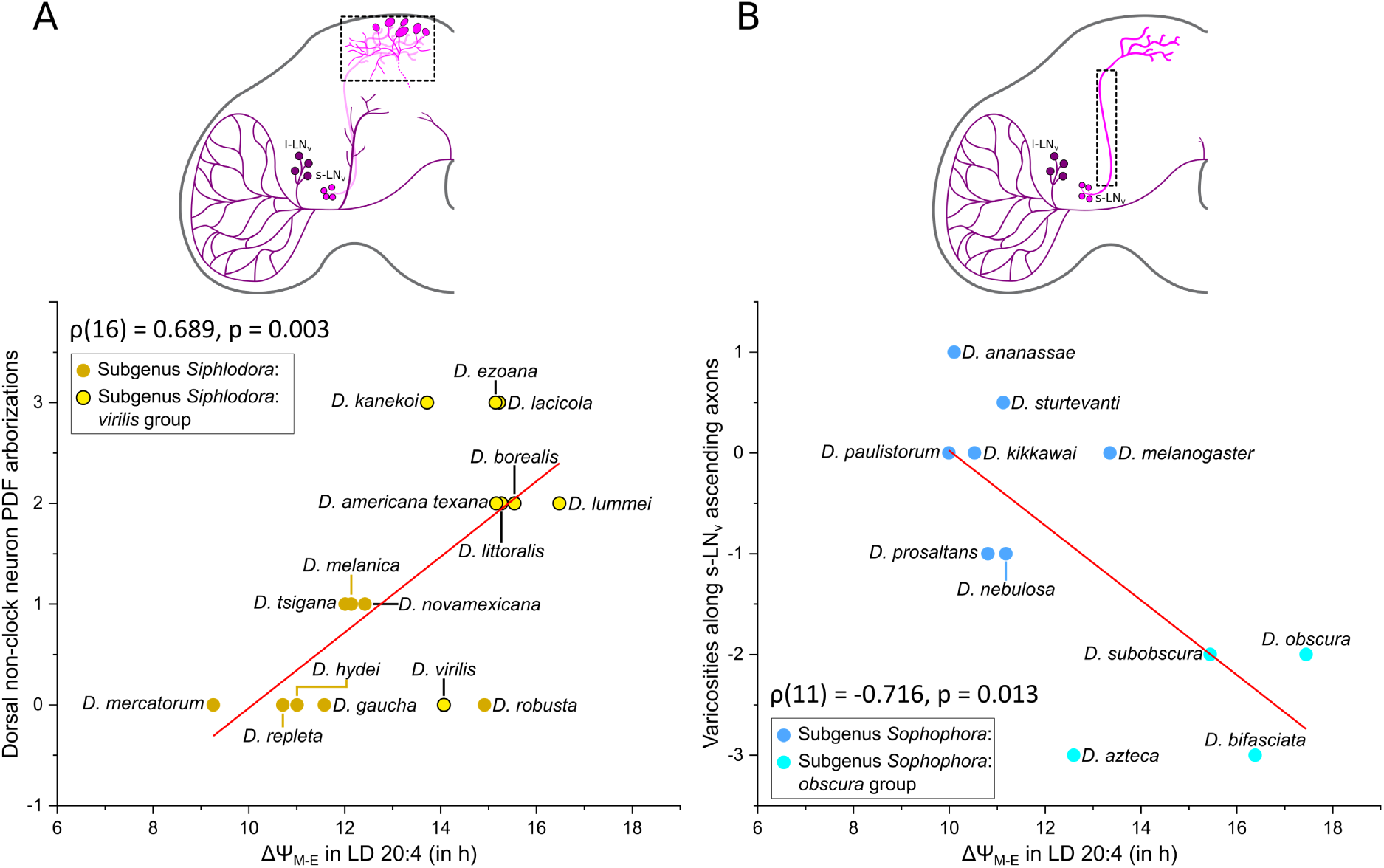
Correlations between the relative abundance of PDF in the dorsal protocerebrum and average ΔΨ_M-E_ within the *Siphlodora* subgenus **(A)**, and between PDF abundance in the ascending s-LN_v_ axons and average ΔΨ_M-E_ within the *Sophophora* subgenus **(B).** The online version of this article includes the following figure supplement for figure 10: **Figure 10—figure supplement 1.** Correlations between the relative abundance of PDF and latitude.

Previous studies have reported that *virilis* group species originating from higher latitudes have less PDF in the s-LN_v_ terminals (Kauranen et al., 2012; Menegazzi et al., 2017; Beauchamp et al., 2018). We found that this negative correlation between PDF presence in the s-LN_v_ terminals and both ΔΨ_M-E_ (Figure 11C) and latitude (Figure 11—figure supplement 1C) held true only within the *Siphlodora* subgenus. In the *Drosophila* subgenus, the trend was instead positive: PDF in the s-LN_v_ terminals increased with latitude (Figure 11—figure supplement 1D) and slightly with ΔΨ_M-E_ (Figure 11D), although without reaching statistical significance. Within the *Sophophora* we did not observe a correlation at all (Figure 11B; Figure 11—figure supplement 1B).

**Figure 11.**
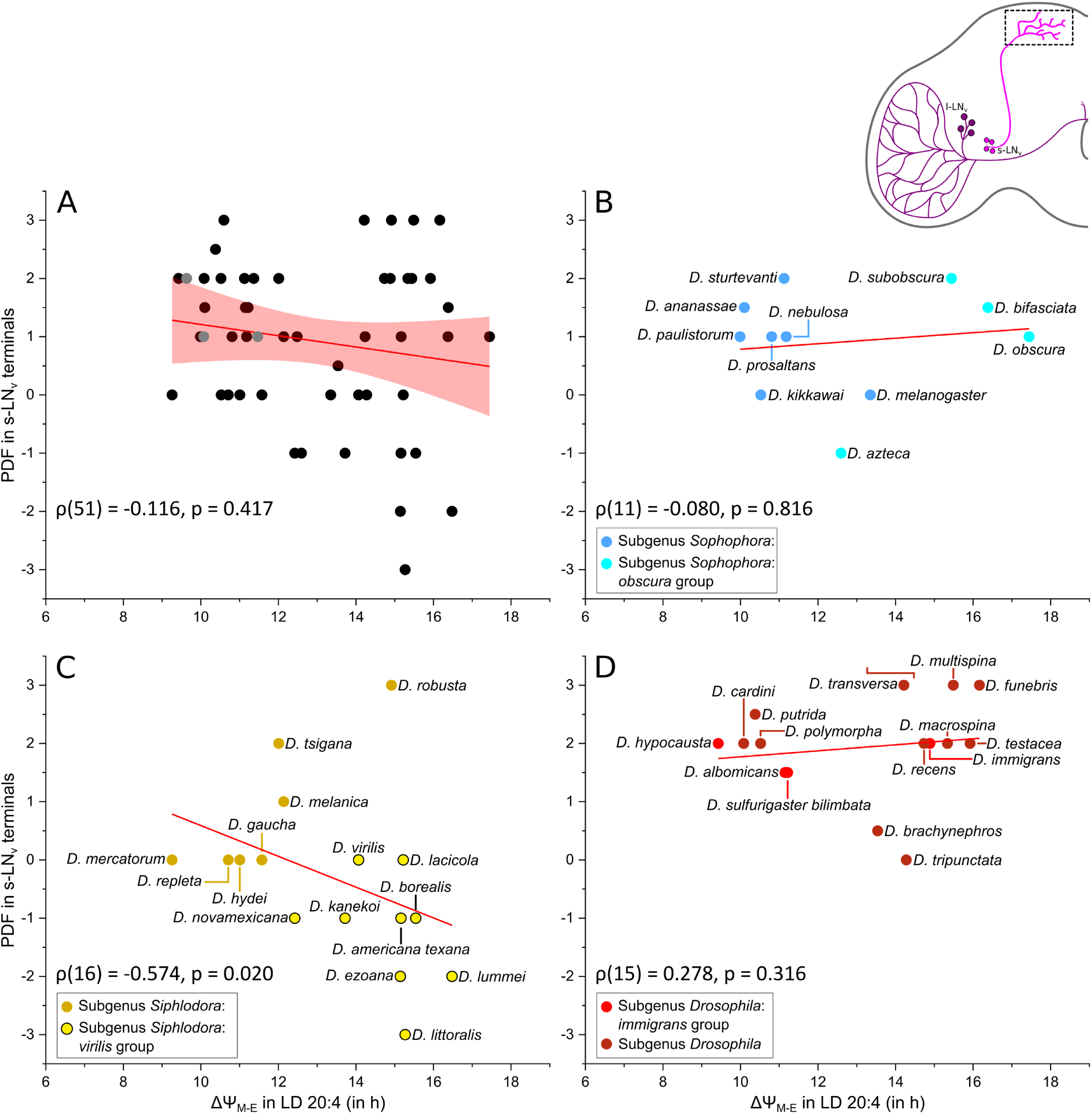
Correlation between PDF in the s-LN_v_ terminals and ΔΨ_M-E_, shown across all *Drosophila* species **(A)** and within the three main subgenera **(B–D)**. PDF staining intensity showed no clear relationship with ΔΨM-E in the *Sophophora* subgenus **(B),** whereas it correlated negatively with ΔΨ_M-E_ in the *Siphlodora* subgenus **(C)** and trended positively in the *Drosophila* subgenus **(D)**. The inset in the upper right corner indicates the quantified brain region. Labeling as in Figure 6. The online version of this article includes the following figure supplement for figure 11: **Figure 11—figure supplement 1.** Correlation between PDF in the s-LN_v_ terminals and latitude.

Again, this result is consistent with earlier observations obtained with selected species of the *virilis-repleta* radiation within the *Siphlodora* subgenus (Beauchamp et al., 2018). Within *D. littoralis* and partly in *D. montana*, both members of the *virilis* group, this same negative correlation was observed across strains originating from different locations: PDF levels in the s-LN_v_ terminals decreased with increasing latitude, while PDF fiber abundance from corazonin-positive neurons increased (Manoli et al., 2025; Manoli et al., 2026).

In summary, we found a strong negative correlation between CRY presence in the l-LN_v_ and both ΔΨ_M-E_ and latitudinal distribution within the species of the *Siphlodora* subgenus, while no such correlation was present in the other subgenera Figure 12). A similar negative correlation existed between PDF staining in the s-LN_v_ terminals and ΔΨ_M-E_ and latitude in *Siphlodora*, whereas the *Drosophila* subgenus demonstrated a positive correlation with latitude only, and the *Sophophora* subgenus showed no correlation at all. *Siphlodora* was also the only subgenus in which PDF fiber density from non-clock (corazonin-positive) neurons close to the s-LN_v_ terminals correlated strongly and positively with both ΔΨ_M-E_ and latitude. In addition, PDF-fiber density in the centrolateral brain correlated positively with both ΔΨ_M-E_ and latitude in *Siphlodora* and *Sophophora*, while PDF in the ascending s-LN_v_ axons correlated strongly and negatively with both measures only in *Sophophora*. (Figure 12). Flies of the *Drosophila* subgenus showed only moderate correIation between PDF in the s-LN_v_ terminals and latitude, which did not extend to ΔΨ_M-E_ (Figure 12).

**Figure 12.**
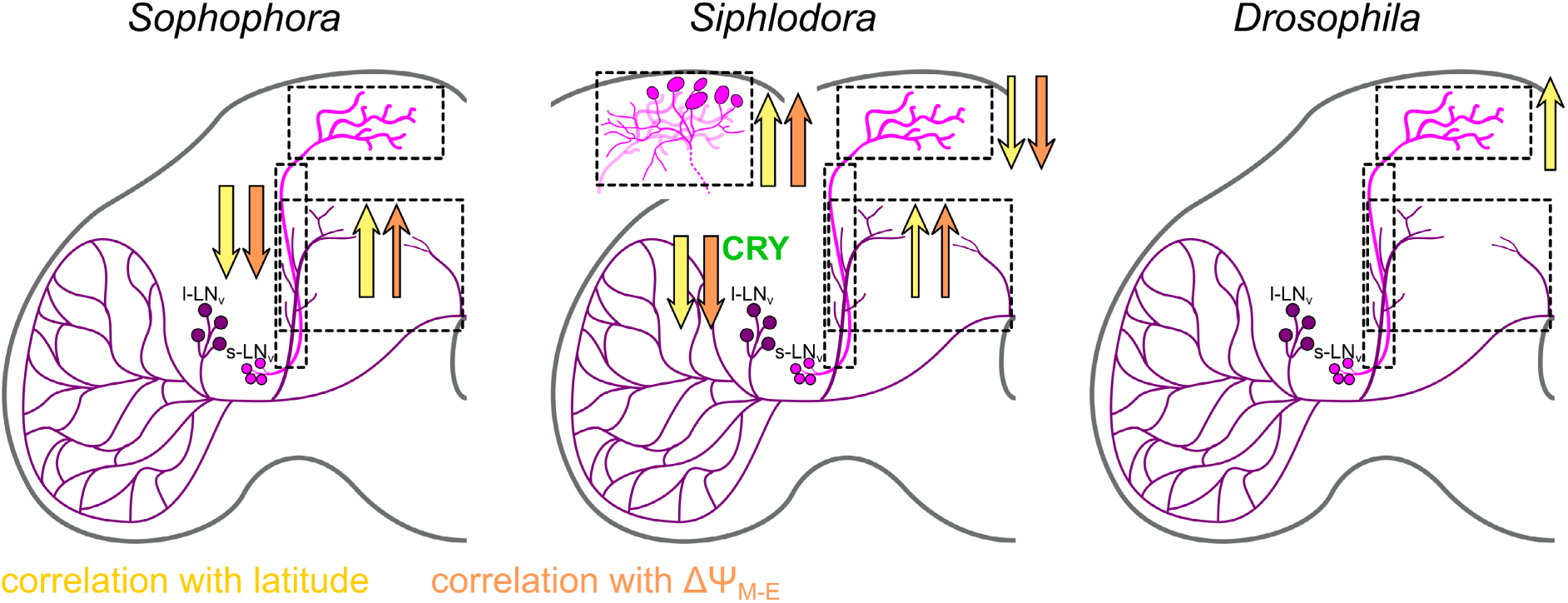
Summary of the correlations between PDF staining (magenta) and latitude/ ΔΨ_M-E_ across the three different *Drosophila* subgenera. Arrow thickness indicates correlation strength; arrow direction indicates the direction of the correlation. Yellow arrows show correlations between PDF and latitude; orange arrows show correlations between PDF and ΔΨ_M-E_. Within the *Siphlodora* subgenus, the expression of CRY in the l-LN_v_ neurons also correlates negatively with ΔΨ_M-E_ and latitude.

Together, these results strongly suggest that the different *Drosophila* subgenera have evolved multiple independent strategies to adapt their circadian clock to high latitudes by improving the ability of the morning and evening oscillators to follow dawn and dusk. Flies of the *Siphlodora* subgenus appear to have achieved this mainly through altered CRY and PDF expression, whereas *Sophophora* species instead show altered PDF arborizations in the brain only. Flies of the *Drosophila* subgenus, in contrast, must have evolved a completely distinct strategy, since none of the CRY or PDF patterns we examined correlated with ΔΨ_M-E_ in this group (Figure 12).

Other possible mechanisms for increasing ΔΨ_M-E_ that we did not investigate here include polymorphisms in clock genes that are not associated with morphological changes. Such polymorphisms have so far mainly been studied in *D. melanogaster* strains originating from different locations and latitudes. For example, two alleles of the *timeless* gene are known (*s-tim* and *ls-tim*), with the frequency of the *ls-tim* allele increasing with latitude (Tauber et al., 2007). The long L-TIM isoform is less light-sensitive than S-TIM and leads to an earlier onset of reproductive dormancy, which is favorable at high latitudes (Sandrelli et al., 2007; Tauber et al., 2007; Abrieux et al., 2020; Lamaze et al., 2022). Flies carrying the *ls-tim* allele also show significantly later evening activity phases, and thus also a larger ΔΨ_M-E_, than low-latitude strains (Deppisch et al., 2022; Lamaze et al., 2022).

Other mechanisms to increase ΔΨ_M-E_ as a response to photoperiod may involve the expression regulation of clock-related genes itself. Shahandeh et al. (2024) found that a mutation in the *cis*-regulatory region upstream of the *Pdf* sequence prevents flies from delaying their evening activity under long photoperiods. This phenotype was observed in *D. sechellia*, a direct sister species of *D. melanogaster* endemic to the equatorial Seychelles islands (≈ 5°S). The authors suggest that this genetic alteration has prevented *D. sechellia* from colonizing higher latitudes.

### Several high-latitude flies have strong circadian clocks

As mentioned in the introduction, one strategy for adapting to very long photoperiods is to increase clock plasticity by weakening its robustness (Abraham et al., 2010). The strength (power) of a circadian clock can be readily measured under constant conditions, such as constant darkness (DD): robust clocks continue to free-run at their own endogenous period, while weak clocks dampen over time until the flies eventually become arrhythmic. An intermediate state between robust rhythmicity and arrhythmicity is complex rhythmicity, characterized by unstable free-running periods that sometimes split into multiple components (see Helfrich-Förster (1998) and Bertolini et al. (2019) for a definition).

Previous studies have shown that high-latitude species of the genus *Chymomyza* and the *virilis* group have weak circadian clocks that dampen quickly and become arrhythmic under DD (Vaze and Helfrich-Förster, 2016; Menegazzi et al., 2017; Beauchamp et al., 2018; Bertolini et al., 2019; Manoli et al., 2025; Manoli et al., 2026). This arrhythmicity is in most cases associated with an extended ΔΨ_M-E_ under long photoperiods. Here, we tested all 52 species for their rhythmic behavior in DD to determine whether this association holds true within the entire Drosophilidae phylogeny.

Representative actograms for all species are shown in Figure 13—figure supplement 1. Rhythmicity ranged from robust, through complex, to weak and arrhythmic, a pattern that held across all three main subgenera and most of the smaller subgroups (Figure 13—figure supplement 1 and 2, Supplementary file 4). The *Siphlodora* subgenus contained the highest number of species with weak rhythms and was the only group in which rhythmicity correlated significantly with latitude: both rhythm power and the percentage of flies with robust rhythms decreased with increasing latitude (Figure 13; Figure 13—figure supplement 3). No such correlation was present when all species were analyzed together (Figure 13; Figure 13—figure supplement 3). These results suggest that the *Siphlodora* subgenus is the only fruit fly clade to have weakened its circadian clock with latitude, likely to allow high plasticity of M and E activity. Indeed, a higher ΔΨ_M-E_ correlated with a lower power of rhythmicity (Figure 13—figure supplement 4).

**Figure 13.**
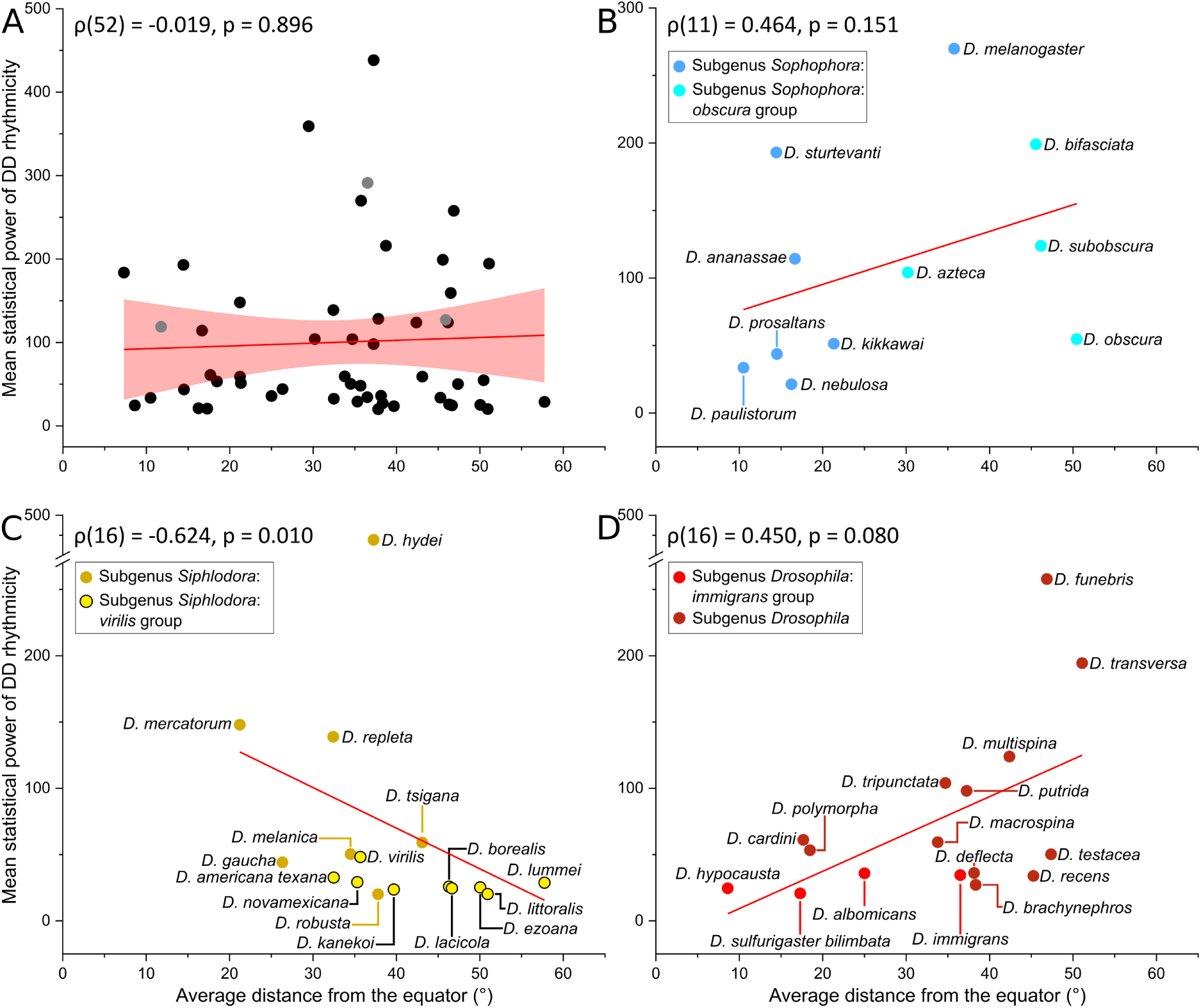
Power of rhythmicity in DD as a function of the mean latitudinal distribution (trimmed by 3 % to exclude potential outlier reports, i.e., false extreme latitudinal values). Spearman’s rank correlation coefficient (ρ) is given in each panel. **(A)** Geographic range does not influence rhythm power across Drosophilidae as a whole. **(B)** and **(D)** No significant correlations between species range and rhythm strength were found in the subgenera *Sophophora* and *Drosophila*. **(C)** Within the *Siphlodora* subgenus, there was a significant negative relationship between latitudinal distribution and rhythm power. The online version of this article includes the following figure supplements for figure 13: **Figure 13—figure supplement 1.** Individual example actograms for all strains. **Figure 13—figure supplement 2.** Percentages of rhythmic flies and rhythm power of all flies. **Figure 13—figure supplement 3.** Correlation between the percentage of rhythmic flies and latitude. **Figure 13—figure supplement 4.** Correlation between ΔΨ_M-E_ and rhythm power. **Supplementary file 4.** Lomb–Scargle analysis results.

Weakened activity rhythms at higher latitudes have also been reported in other insects, including the beetle *Tribolium castaneum* (Abe et al., 2021), the pitcher plant mosquito, *Wyeomyia smithii* (Bradshaw et al., 1998), the parasitoid hymenopteran *Leptopilina heterotoma* (Fleury et al., 1995) and the Antarctic midge, *Belgica antarctica* (Kobelkova et al., 2015). Even some vertebrates, such as Antarctic fish (*Notothenia coriiceps*, *Cottus poecilopus*, and *Myoxocephalus scorpius*) (Wright et al., 2025), the black-legged kittiwake (*Rissa tridactyla*) (Gillies et al., 2026) and reindeer (van Oort et al., 2005), develop arrhythmic activity patterns in long summer days. A recent comparative genomic study of biological clock gene loss and circadian rhythms in fishes found widespread, lineage-specific loss of core clock genes, particularly in convergently evolved polar species (Wright et al., 2025). This trend was significantly correlated with latitude, with higher latitude demonstrating greater circadian gene loss.

Nevertheless, some animals remain robustly rhythmic even at high latitudes. Bumblebees foraging in arctic regions, for instance, show strong activity rhythms, likely because a functional circadian clock is required for sun-compass orientation (Stelzer and Chittka, 2010). Similarly, Arctic ground squirrels (*Urocitellus parryii*) maintain daily rhythms of physiology and behavior throughout the summer, which includes 6 weeks of constant daylight (Williams et al., 2017). These animals retain a very robust circadian clock in the suprachiasmatic nucleus. Their high rhythmicity is likely maintained to reduce thermoregulatory costs, since their thermoneutral zone occurs only during midday. We suspect that the robust rhythmicity we observed in several drosophilids originating from high latitudes (e.g., *D. funebris*, *D. obscura,* and *D. transversa*; Figure 14, Figure 13—figure supplement 2) likewise confers a selective advantage that remains to be clarified. A recent paper shows that in cyanobacteria robust circadian clock at high latitudes is required to align ultraviolet resistance with the appropriate time of day under long photoperiods (Schober et al., 2026), giving robust clocks a fitness advantage over weak clocks at high latitudes in this species (Seki and Ito, 2022; reviewed by Cai and Chiu, 2026; Dani et al., 2026). Future studies will need to determine what fitness advantages the robust clocks of some high-latitude *Drosophila* species provide.

**Figure 14.**
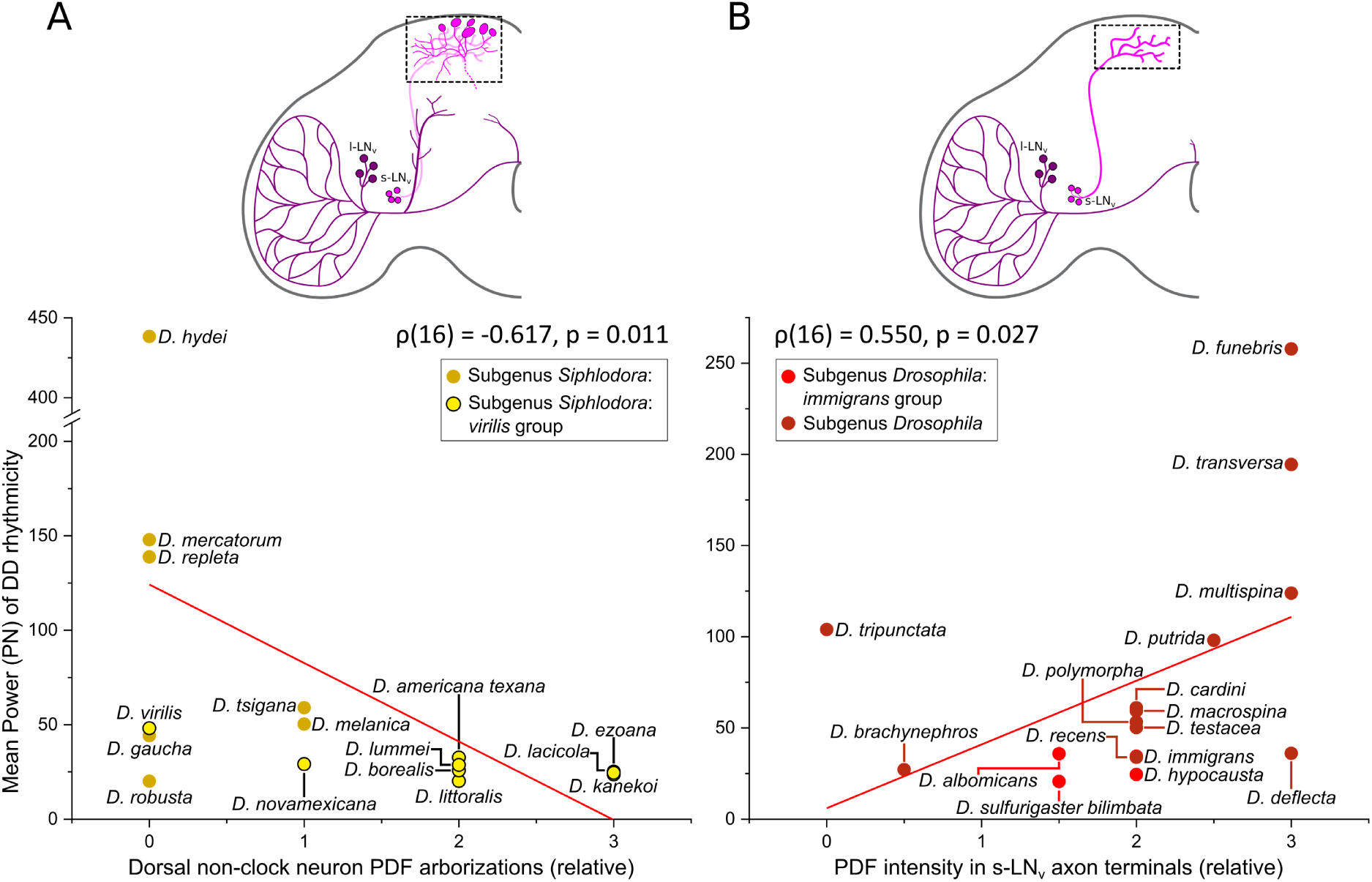
Power of rhythmicity in DD as a function of PDF expression features in the superior protocerebrum of the *Siphlodora* and *Drosophila* subgenera. Spearman’s rank correlation coefficient (ρ) is given in each panel. The insets at the top indicate the quantified brain region. **(A)** A moderate negative correlation between mean power and the PDF intensity in dorsal non-clock CRZ neurons in the *Siphlodora* subgenus. **(B)** A moderate correlation between mean power and PDF intensity in the s-LN_v_ terminals within the *Drosophila* subgenus. The online version of this article includes the following figure supplements for figure 14: **Figure 14—figure supplement 1.** PDF staining in the s-LN_v_ terminals and CRZ neurons in relation to rhythm power. **Supplementary file 5.** Correlation between period and latitude across the whole species set.

### PDF in the superior protocerebrum is important for robust activity rhythms under DD conditions

Previous studies have shown that PDF presence in the s-LN_v_ terminals in the superior protocerebrum is crucial for rhythmicity in DD, whereas PDF released in a non-rhythmic manner from non-clock neurons close to the s-LN_v_ terminals may disrupt rhythmicity (Helfrich-Förster, 1998; Renn et al., 1999; Helfrich-Förster et al., 2000; Bertolini et al., 2019). We therefore tested whether rhythm power correlated negatively with PDF in the nearby corazonin-positive (CRZ) neurons and positively with PDF levels in the s-LN_v_ terminals. This was indeed the case for the *Siphlodora* subgenus: PDF staining intensity from the CRZ neurons near the s-LN_v_ terminals correlated significantly and negatively with rhythm power (Figure 14A) and robust rhythmicity. This negative correlation was particularly pronounced within the *virilis* group (ρ = −0.709, p = 0.032) (Figure 14—figure supplement 1). Furthermore, in the *Siphlodora* subgenus, PDF in the s-LN_v_ terminals correlated significantly with robust rhythmicity in DD (ρ = 0.544, p = 0.030) (Figure 14—figure supplement 1), and showed a weaker, non-significant positive correlation with rhythm power (ρ = 0.405, p = 0.120) (Figure 14—figure supplement 1).

In contrast, in the *Drosophila* subgenus, PDF staining intensity in the s-LN_v_ terminals correlated positively with rhythm power (Figure 14B).

Overall, our findings support a prominent role for PDF in the superior protocerebrum in maintaining robust activity rhythms under DD (Helfrich-Förster, 1998; Renn et al., 1999; Helfrich-Förster et al., 2000). PDF released from different sources in the brain, the l-LN_v_ in *Sophophora* or the CRZ neurons in subgenus *Siphlodora*, appears in both cases to delay evening activity under long photoperiods and thereby increase ΔΨ_M-E_. A decrease in PDF at the s-LN_v_ terminals correlated with weak rhythmicity in *Siphlodora*, whereas high PDF levels in these terminals correlated with robust rhythmicity in the *Drosophila* subgenus. No such correlation was detected in *Sophophora*; however, genetical removal of PDF from the s-LN_v_ terminals in *D. melanogaster* flies consistently lead to arrhythmicity (Helfrich-Förster, 1998; Renn et al., 1999; Grima et al., 2004). This suggests that PDF is constitutively present in *Sophophora* flies regardless of latitude, yet remains essential, as its absence still causes arrhythmicity in this subgenus. Arrhythmicity could also result from a loss of circadian PDF cycling in the s-LN_v_ terminals, as demonstrated in the fly species *Chymomyza costata* (Bertolini et al., 2019), This mechanism would have escaped our attention, as we assessed PDF only in the morning when it was close to its expected maximum level.

Nevertheless, our studies also revealed that not all *Drosophila* species from high latitudes have reduced PDF levels in the s-LN_v_. Species of the *Drosophila* subgenus, in fact, tend to show increasing PDF staining intensity in the s-LN_v_ terminals with increasing latitude, while maintaining robust rhythms in DD.

### The period length of the free-running rhythm in DD does not systematically correlate with latitude

Finally, we tested whether the free-running period length in DD correlates with latitude among rhythmic flies, since such a correlation has been reported for eclosion rhythms in several *Drosophila* species (reviewed in Hut et al., 2013), as well as for activity rhythms of linden bugs (Pivarciova et al., 2016) and jewel wasps (Paolucci et al., 2019). We found no systematic correlation between period and latitude across the whole species set, nor within the *Siphlodora* subgenus alone (Supplementary file 5). Such correlations may nonetheless still exist within strains of the same species sampled across different latitudes, since all the studies cited above examined latitudinal clines within single species rather than across species. Eclosion rhythm periods in *D. subobscura*, *D. littoralis*, and *D. auraria* shorten with increasing latitude (Lankinen, 1986; Pittendrigh and Takamura, 1989; Lankinen, 1993), while adult activity rhythm periods in linden bugs and jewel wasps lengthen with increasing latitude (Pivarciova et al., 2016; Paolucci et al., 2019), showing that latitude can influence period to shift in opposite directions. A study of *Tribolium castaneum* strains spanning a latitudinal gradient across Japan found no systematic change in period with latitude at all (Abe et al., 2021). Altogether these findings show that the effect of latitude on the free-running period varies considerably between species, and no single, generalizable pattern can be expected across different species or populations.

### Limitations of the study

We are aware that the results of this study are purely correlational in nature and do not necessarily indicate a causal relationship. However, a previous study demonstrated that mimicking the PDF and CRY expression pattern typical of high-latitude *virilis*-group species in the brain of low-latitude *D. melanogaster* was sufficient to reproduce the rhythmic behavior typical of high-latitude flies (Menegazzi et al., 2017). This makes it likely that at least some of the correlations observed here reflect genuine causal relationships in nature. It is also probable that PDF and CRY are not the only clock-related factors involved in the adaptation of the fly’s circadian system to high latitudes. The circadian clocks of all animals studied to date contain a variety of neuropeptides with diverse functions (Ono et al., 2021; Tsujino et al., 2022; Reinhard et al., 2024; Neupert et al., 2025). They act as mediators for synchronization among individual clock neurons, in entrainment to light, and as output regulators. Future research must clarify their role in high-latitude adaptation.

Another important limitation of our study is that we restricted our investigation to photoperiod and did not consider ambient temperature. Temperature changes markedly with latitude, and the circadian clock is known to adapt to such changes through thermosensitive splicing of the *period* and *timeless* genes (Majercak et al., 1999; Majercak et al., 2004; Martin Anduaga et al., 2019). Furthermore, we did not account for the altitude from which the different species originated. Like latitude, altitude affects environmental temperature and circadian clock adaptation (Vanlalhriatpuia et al., 2007; Cao and Edery, 2017).

## Conclusions

Our correlational study of circadian clock traits, both clock-related behavior and the underlying clock network in the brain, and geographical origin across 52 *Drosophila* species revealed significant associations between various parameters and latitude. Clock plasticity, measured as the phase difference between morning and evening activity (ΔΨ_M-E_), increased with geographical latitude across the entire Drosophila radiation, including all three main subgenera – *Sophophora*, *Siphlodora,* and *Drosophila*. High clock plasticity can be considered a prerequisite adapting daily behavior to rapidly changing photoperiods at high latitudes, a pattern also observed in mammalian species (Abraham et al., 2010). One route to achieve high clock plasticity is to reduce the robustness of the circadian clock, and indeed, this appears to be the case in several mammals and in *Siphlodora* drosophilids. Fly species of the subgenus *Drosophila*, however, showed the opposite, albeit weak, relationship between clock robustness and latitude: clock strength increased moderately with latitude rather than decreasing, whereas in *Sophophora*, rhythm strength did not change with latitude at all. This is consistent with observations in other species (Abraham et al., 2010; Stelzer and Chittka, 2010; Williams et al., 2017; Cai and Chiu, 2026) and demonstrates that multiple strategies exist for generating clocks suited to control activity rhythms at high latitudes. Given that the three *Drosophila* subgenera diverged at least 40 million years ago, it is reasonable that they evolved different mechanisms for adapting their circadian systems to high latitudes. Despite this diversity of strategies, the neuropeptide PDF, and, in *Sophophora* and *Siphlodora*, the blue-light photopigment CRY, appears to play a crucial role in this adaptation across subgenera. Future research will need to clarify the contribution of other clock-related factors in the adaptation of daily activity to high latitudes.

## Material and Methods

### Fly strains and maintenance

The majority of the 52 tested fly species was obtained from the National *Drosophila* Species Stock Center (NDSSC, Cornell University, NY, USA) and Kyorin-Fly at Kyorin University (Tokyo, Japan). Additional species were provided by individual laboratories. *D. cardini*, *D. recens*, *D. transversa*, and *D. tripunctata* were supplied by Clare Scott Chialvo (Appalachian State University, Boone, NC, USA). *D. testacea* was provided by Steve Perlman (University of Victoria, BC, Canada), and *D. lummei*, *D. ezoana*, and *D. littoralis* by Anneli Hoikkala (University of Jyväskylä, Finland). *D. subobscura* was collected in 2020 in Gerbrunn near Würzburg, Germany. The *D. hydei* and *D. mercatorum* lines originated from Tanzania and were first described in Beauchamp et al. (2018). Furthermore, between 2017 and 2020, several species were collected by the first author near the villages of Oellingen and Sonderhofen (close to Würzburg, Bavaria, Germany). These included *D. bifasciata*, *D. busckii*, *D. immigrans*, *D. repleta*, *Scaptomyza pallida*, and two strains of *D. funebris*.

Species’ identity was confirmed using complementary approaches. Diagnostic morphological traits were used for *D. busckii*, *D. immigrans*, *D. repleta*, and *D. funebris*. Crosses with established wild-type strains were performed for *D. bifasciata* and *D. funebris*. For *D. immigrans*, *D. repleta*, *D. bifasciata*, *D. subobscura*, and *S. pallida*, species identity was further verified by sequencing conserved mitochondrial or nuclear marker genes. Primers were adapted from previously published studies (Simon et al., 1994; Bonacum et al., 2001; Gao et al., 2007; Lapoint et al., 2013; Goldman-Huertas et al., 2015). Detailed information about strain origins is given in Supplementary file 6A.

### Fly husbandry

All species were reared in plastic vials (4.5 cm diameter, 10 cm height) containing approximately 2 cm of food medium. Coffee filter paper or cotton lint was typically placed in the vials to provide surfaces for larval development and pupation and to give adult flies resting sites, reducing the risk of drowning in liquefied food. These materials also helped regulate moisture and humidity within the vials.

Detailed rearing conditions for each species are provided in Supplementary file 6B. Most species were maintained in climate chambers under a 12 h:12 h light–dark cycle (LD 12:12) at either 18 °C or 25 °C with 60 % relative humidity. Some strains, particularly those originating from northern regions, were kept at 21 °C under a long photoperiod (LD 20:4) to prevent reproductive diapause.

Most species were reared on a standard *Drosophila* medium composed of 14.75 % (m/v) cornmeal, 1.00 % soy flour, 1.85 % brewer’s yeast, 0.63 % agar, 4.50 % malt syrup, 4.50 % sugar beet molasses, and 0.25 % methyl paraben. Some tropical species were maintained on a banana medium consisting of the standard cornmeal medium supplemented with approximately two mashed bananas per liter. Mushroom-feeding species were provided with standard medium topped with 4–5 g of squeezed, sliced button mushrooms.

### Locomotor activity recording

Locomotor activity was measured using the *Drosophila* Activity Monitoring (DAM) system (TriKinetics Inc.). Individual flies were anesthetized with CO₂ and placed in glass tubes (65 mm length). One third of each tube was filled with food (2 % agar and 4 % sucrose), and the opposite end was sealed with an air-permeable foam plug. The tubes were inserted into activity monitors (up to 32 tubes per monitor). An infrared beam positioned at the center of each slot recorded every crossing by the fly. Beam breaks were counted in 1-minute intervals and exported as raw text files (.txt) for each monitor.

Male flies were collected 1–7 days after eclosion and transferred to the activity tubes under CO₂ anesthesia. Because the examined species differed in body size, larger species that could not easily turn around in the standard tubes (5 mm external diameter) were recorded in wider tubes (7 mm external diameter). The tube size used for each species is listed in Supplementary file 6C.

Activity was recorded in PVC light boxes equipped with white LEDs. Light intensity was set to approximately 120 lx at the top row of monitors. To mimic natural dawn and dusk conditions, light intensity increased gradually from 0 to 120 lx at the start of each light phase and decreased from 120 to 0 lx at the end, each over a 90-minute interval. Flies were first recorded for two days under LD 12:12 to acclimate to the setup. They were then exposed sequentially to LD 16:8 for seven days and LD 20:4 for seven days. At least two independent experiments were conducted using more than one monitor, either in different light boxes or in separate experimental runs.

For experiments focusing on free-running rhythms, flies were recorded for eight days under LD 12:12 and then transferred to constant darkness for at least eleven days.

### Average day analysis

For each photoperiodic condition, the average daily activity of individual flies was calculated from the last five days of recording. This analysis used a custom macro written in Microsoft Excel by Taishi Yoshii (Okayama University). Activity data from the monitoring system were retained at their original 1-minute resolution. For each fly, this produced a dataset of 1440 values representing the mean activity for every minute of a 24-hour cycle averaged across five days.

Individuals that did not survive until the end of a photoperiodic condition were excluded manually. Mean activity profiles were then calculated for each species and strain by averaging across all remaining flies. Data for LD 12:12 were taken from the entrainment phase of the constant darkness experiments. Average day graphs were plotted using OriginPro (version 2022b).

### Determination of morning and evening activity in different *Drosophila* species

Morning and evening activity peaks were identified following the approach described in Deppisch et al. (2022). Peaks were fitted from the average day profiles using the Quick Fit function in OriginPro. Gaussian curves were fitted to sections of the activity profile surrounding obvious local maxima, and the x_c_ value of the fit was used to determine the timing of each peak.

All visible peaks were initially identified for the three photoperiods (LD 12:12, LD 16:8, and LD 20:4), but only genuine circadian morning and evening peaks were included in further analyses. Maxima caused by masking effects were excluded. The main masking events were the onset and offset of light, which produced startle responses. In some cases, particularly in the morning, circadian peaks coincided with the light onset response.

Fits were generally performed on unsmoothed mean profiles. In rare cases where large activity spikes interfered with peak detection, profiles were smoothed using a moving average of 11 before fitting. In three species (*D. kikkawai*, *D. kanekoi*, and *D. sulfurigaster*), some individuals did not synchronize to LD 20:4; only flies that showed clear entrainment in their actograms were included. Most *D. deflecta* individuals also failed to entrain to LD 20:4. Even after removing unsynchronized flies, the remaining population displayed an atypical, almost unimodal activity pattern. For that reason, this species was excluded from analyses under LD 20:4.

The identified morning and evening peaks were used to calculate the morning–evening phase difference (ΔΨ_M-E_) for each photoperiodic condition.

### Analysis of rhythmicity in constant conditions (DD)

Rhythmicity and free-running periods under constant darkness were evaluated using the Lomb–Scargle periodogram. The analysis was performed with a custom R script. For each fly, the first day in DD following the entrainment phase was excluded, and the subsequent 12 days were analyzed. Only individuals that survived throughout this period were included.

The resulting free-running periods (τ) were verified by visual inspection of individual actograms generated with the ImageJ plug-in ActogramJ (Schmid et al., 2011). This step was necessary because several species displayed lower locomotor activity than the commonly studied *D. melanogaster*, which can reduce the statistical sensitivity of periodogram analyses.

Rhythm quality was also assessed manually based on the actograms. Individuals were classified as clearly circadian or as showing weak, complex, or longer-period rhythms Bertolini et al. (2019). When multiple significant period components were detected in the periodogram and this pattern occurred consistently across most individuals of a strain, the period with the higher average power was considered for further analysis.

For species with very low activity levels or complex rhythmic patterns, the evaluation relied more strongly on actogram inspection, because periodogram methods can become unreliable with sparse data. In such cases, flies that showed obvious rhythmic patterns in the actograms were classified as rhythmic even if the Lomb–Scargle analysis did not yield a statistically significant period. Conversely, periods were accepted when the power value exceeded 10 and the corresponding peaks in the periodogram matched the rhythmic patterns observed in the actograms.

### Immunohistochemistry for PDF and CRY

In each species, immunohistochemical stainings were performed on brains of young male flies that had been entrained for 5–7 days under LD 12:12 at 20 °C. Flies were collected in darkness at ZT 23.5. This sampling time was chosen based on the temporal dynamics of the target proteins CRY and PDF.

In *D. melanogaster*, CRY accumulates during the dark phase and is degraded upon light exposure (Peschel et al., 2009; Ozturk et al., 2013). Sampling near the end of the scotophase therefore maximizes CRY protein levels. In addition, this time point is close to ZT 2, when complex and intensely stained axon terminals of PDF neurons are prominent in *D. melanogaster* (Park et al., 2000; Fernández et al., 2008).

Flies were fixed in darkness in 4 % paraformaldehyde prepared in PBST (phosphate-buffered saline containing 0.5 % Triton X-100) for 2.5–3.5 h. Fixation time varied among species because differences in body size affected the rate at which the fixative penetrated the brain. Exact fixation times for each species are listed in Supplementary file 6C. After fixation, samples were washed three times with PBS for 10 min each. Brains were then removed in cold PBS using forceps. Dissected brains were transferred to a 96-well plate and washed three times in PBS and three times in PBST. Samples were subsequently blocked in 5 % normal goat serum (NGS; in PBST containing 0.02 % NaN₃) overnight at 4 °C, followed by an additional 6 h at room temperature. Brains were then incubated with the primary antibody solution for 1 day at room temperature and for three additional nights at 4 °C. All antibody solutions were prepared in PBST containing 5 % NGS and 0.02 % NaN₃.

The standard primary antibody mixture contained a polyclonal rabbit anti-dCRY (1:1000; Yoshii et al., 2008) and a monoclonal mouse anti-PDF (C7 line; Developmental Studies Hybridoma Bank; 1:2000). In species where this PDF antibody did not produce reliable staining, an alternative combination was used consisting of polyclonal guinea pig anti-CRY (1:200; Benito et al., 2008)) and polyclonal rabbit anti-βPDH (1:400; Dircksen et al., 1987). The βPDH antibody was raised against the pigment-dispersing hormone of the crustacean *Uca pugilator* and has been shown to label PDF-positive neurons effectively in multiple drosophilid species (Helfrich-Förster and Homberg, 1993; Hermann et al., 2013). The specific PDF antibody used for each species is listed in Supplementary file 6C.

After primary antibody incubation, brains were washed six times in PBST. Secondary antibody solutions contained Alexa Fluor^TM^-conjugated goat antibodies (anti-rabbit 488 nm with anti-mouse 635 nm, or anti-guinea pig 488 nm with anti-rabbit 635 nm). Secondary antibodies were applied for 2 h at room temperature and overnight at 4 °C in darkness. Samples were then washed three times in PBST and three times in PBS. Finally, brains were mounted on Superfrost^TM^ microscope slides in VECTASHIELD^®^ antifade mounting medium (VECTOR Laboratories) and sealed with Fixogum industrial glue (Marabu).

Brain samples were imaged using a Leica TCS SPE confocal microscope (Leica Microsystems). For each specimen, at least one complete brain hemisphere was scanned using a 20× objective (ACS APO 20.0×/0.60 IMM) with glycerol-based immersion medium. Image stacks were processed with Fiji, an open-source distribution of ImageJ (Schindelin et al., 2012). Each species was examined for its PDF and CRY expression pattern by the eyes of experienced *Drosophila* chrono- and neurobiologists. Multiple brains per species were examined. For PDF, the presence of the peptide was assessed in the s-LN_v_ and l-LN_v_ clock neuron clusters, as well as in potential additional dorsal non-clock neurons. The extent of axonal varicosities along the dorsal projection of s-LN_v_ neurons was evaluated, together with the density of their axon terminals in the dorsal protocerebrum. In addition, PDF abundance in the centrolateral brain was assessed. The analyzed brain regions are illustrated in Figure 1.

Expression patterns were scored relative to the reference species *D. melanogaster*. The staining pattern in this species was defined as the baseline (score 0), and lower or higher signal levels were scored on a scale from −3 to +3 (see Table S6, Appendix section 6.6).

For CRY, the presence of the protein was evaluated across major clock neuron clusters, with particular attention to the lateral neurons and the DN1 cluster. CRY staining intensity in the l-LN_v_ cluster was scored from −3 (absent or nearly absent) to 0 (comparable to *D. melanogaster*).

### Generation and analysis of global distribution data and extraction of Drosophila distribution data

Geographic distribution data for the analyzed species were obtained from TaxoDros, curated by Gerhard Bächli ((Bächli, 2024), https://www.taxodros.uzh.ch/). This database compiles distribution records for most known members of the family Drosophilidae, including geographic coordinates extracted from published studies and collection reports dating back to the early 20^th^ century.

Coordinates were retrieved from the website version dated 10 January 2024 using the pegi3s/taxodros_gis image from the Bioinformatics Docker Images Project https://pegi3s.github.io/dockerfiles/ (López-Fernández et al., 2021). The extracted dataset contained decimal latitude and longitude coordinates for all available records.

All entries were manually reviewed to correct obvious errors, such as swapped latitude and longitude values or locations falling offshore. The coordinates of the northernmost and southernmost records were also verified and corrected when necessary; for example, reports associated with tropical fruit stores or markets were excluded. If not already present in the database, the collection sites of the strains used in this study were added to the dataset.

Additionally, the North American records of *D. transversa* were deleted as they are considered misinterpretations for other *quinaria* group species (Wheeler, 1960; Ginsberg et al., 2019). Likewise, the Eurasian *D. testacea* was misinterpreted for *D. neotestacea* in North America (Grimaldi et al., 1992; Werner et al., 2018). They were removed accordingly. Scarse records of *D. borealis* and *H. duncani* were complemented with non-redundant data from the Global Biodiversity Information Facility (GBIF, https://www.gbif.org/).

### Data analysis

To avoid complications arising from positive and negative latitude values when calculating species averages across hemispheres, all latitude coordinates were converted to absolute values. For each species, we calculated the arithmetic mean, the median, and the most poleward record (typically the northernmost record, except for *D. gaucha*).

To reduce the influence of potential outliers in the database, we additionally calculated a 3 % trimmed mean of the latitude values. These numbers were later used in the figures containing the average distance from the equator. Likewise, instead of relying solely on a single extreme record, we determined the 97^th^ percentile of the latitude distribution as an additional estimate of the poleward range limit.

### Correlation

Spearman’s rank correlations were calculated for different parameters using OriginPro.

## Supporting information

Supplementary file 1_SpeciesDistributionMaps

Supplementary file 2_GeographicDistributionStatistics

Supplementary file 3_M-,E-peaks

Supplementary file 4_DD_statistics_Lomb-Scargle

Supplementary file 5_Period_AvLatitude_Corr

Supplementary file 6_Strain_Information

Main Figures in High Resolution

## Resource availability

All data supporting this study are available within the main text or the Supplemental Information. Additional details can be obtained from the first author upon request.

## Acknowledgements

We thank Hanna Ullrich for help with locomotor activity recordings, and Michelle Adam for immunohistochemical experiments. We thank Anneli Hoikkala, Pamela Menegazzi, Steve Perlman, and Clare Scott Chialvo for providing fly strains, and Enrico Bertolini for providing an R-script used in rhythmicity analysis. We also acknowledge Giulia Manoli and Taishi Yoshii for critical comments on the manuscript. The work was supported by the German Research Foundation (DFG), FO207/15-2 and the DAAD (BMFTR - P33 - Project-ID: 57834638).

## Author contribution

P.D. designed and performed all experiments and wrote the first version of the manuscript, F.S.C. was involved in immunohistochemistry and confocal analysis, and C.H.-F. contributed to conceptual discussions, provided critical input throughout the project, revised the manuscript and acquired funding. All authors reviewed and approved the final manuscript.

## Declaration of interest

The authors declare no competing interests.

## Declaration of generative AI and AI-assisted technologies in the writing process

During the preparation of this work, the authors used Claude AI and DeepL in order to improve the language of the manuscript.

## Notes

### Competing Interest Statement

The authors have declared no competing interest.

