## Supplementary file 2_GeographicDistributionStatistics for "Fruit flies of the subgenera *Drosophila*, *Siphlodora* and *Sophophora* use different mechanisms to adapt their daily activity to long photoperiods"

### Geographical distribution data of the different fly species

**Supplementary file 2A** Descriptive statistics of the latitudinal data for each species' reported collection sites (as recorded in Taxodros).

| **Species** | **N (Geographic coordinates)** | **Mean absolute latitude of distribution [°]** | **Standard deviation** | **Standard error of the mean** | **Trimmed mean (3 %) of the absolute latitude [°]** | **Median of absolute latitudes [°]** | **Latitude of the polarmost report [°]** | **97^th^ percentile (“trimmed polarmost report”) [°]** |
| --- | --- | --- | --- | --- | --- | --- | --- | --- |
| *S. latifasciaeformis* | 269 | 11.90 | 8.70 | 0.53 | 11.77 | 10.00 | 33.73 | 27.63 |
| *S. lebanonensis* | 44 | 36.53 | 5.09 | 0.77 | 36.53 | 37.28 | 46.00 | 44.51 |
| *S. rufifrons* | 269 | 45.92 | 5.51 | 0.34 | 45.90 | 46.13 | 62.55 | 57.10 |
| *H. duncani* | 84 | 37.80 | 4.54 | 0.50 | 37.82 | 38.47 | 47.12 | 45.00 |
| *D. paulistorum* | 288 | 10.63 | 7.69 | 0.45 | 10.51 | 8.62 | 30.26 | 27.60 |
| *D. nebulosa* | 386 | 16.34 | 9.42 | 0.48 | 16.25 | 18.11 | 42.42 | 31.49 |
| *D. prosaltans* | 212 | 14.54 | 8.94 | 0.61 | 14.52 | 13.04 | 31.49 | 29.52 |
| *D. sturtevanti* | 333 | 14.47 | 8.72 | 0.48 | 14.45 | 15.00 | 31.49 | 28.44 |
| *D. melanogaster* | 4560 | 35.60 | 13.06 | 0.19 | 35.75 | 37.59 | 73.24 | 55.30 |
| *D. kikkawai* | 342 | 21.29 | 9.36 | 0.51 | 21.33 | 23.28 | 41.72 | 35.23 |
| *D. ananassae* | 818 | 16.70 | 8.95 | 0.31 | 16.68 | 16.99 | 41.66 | 31.30 |
| *D. obscura* | 833 | 50.54 | 6.04 | 0.21 | 50.45 | 49.00 | 69.95 | 65.02 |
| *D. bifasciata* | 417 | 45.60 | 9.84 | 0.48 | 45.55 | 45.00 | 69.44 | 66.41 |
| *D. subobscura* | 1590 | 46.14 | 7.10 | 0.18 | 46.16 | 46.31 | 64.73 | 58.20 |
| *D. azteca* | 110 | 30.10 | 9.46 | 0.90 | 30.21 | 33.32 | 44.93 | 41.83 |
| *D. busckii* | 1385 | 38.52 | 11.52 | 0.31 | 38.73 | 39.12 | 66.37 | 55.76 |
| *S. pallida* | 1976 | 46.47 | 8.18 | 0.18 | 46.51 | 46.59 | 70.51 | 62.17 |
| *D. robusta* | 413 | 37.80 | 4.80 | 0.24 | 37.80 | 38.40 | 48.35 | 46.34 |
| *D. tsigana* | 35 | 43.10 | 3.92 | 0.66 | 43.10 | 43.15 | 50.32 | 48.30 |
| *D. melanica* | 221 | 34.57 | 4.79 | 0.32 | 34.51 | 33.48 | 48.35 | 45.23 |
| *D. gaucha* | 31 | 26.33 | 11.00 | 1.98 | 26.33 | 29.20 | 37.92 | 37.92 |
| *D. hydei* | 1169 | 37.06 | 12.34 | 0.36 | 37.26 | 38.31 | 73.24 | 55.04 |
| *D. repleta* | 793 | 32.37 | 13.56 | 0.48 | 32.43 | 32.38 | 68.53 | 52.40 |
| *D. mercatorum* | 296 | 21.40 | 11.13 | 0.65 | 21.22 | 22.04 | 56.56 | 48.45 |
| *D. kanekoi* | 28 | 39.69 | 5.86 | 1.11 | 39.69 | 42.16 | 50.32 | 50.32 |
| *D. ezoana* | 83 | 50.10 | 9.60 | 1.05 | 50.05 | 45.00 | 68.54 | 66.48 |
| *D. littoralis* | 319 | 50.98 | 7.37 | 0.41 | 50.94 | 48.43 | 69.51 | 66.48 |
| *D. virilis* | 388 | 35.49 | 8.46 | 0.43 | 35.69 | 35.65 | 55.45 | 49.22 |
| *D. lummei* | 39 | 57.75 | 8.84 | 1.42 | 57.75 | 61.05 | 68.54 | 67.58 |
| *D. novamexicana* | 18 | 35.31 | 2.85 | 0.67 | 35.31 | 34.46 | 39.33 | 39.33 |
| *D. americana texana* | 75 | 32.49 | 2.41 | 0.28 | 32.49 | 32.32 | 37.32 | 36.47 |
| *D. borealis* (eastern) | 28 | 46.31 | 6.29 | 1.19 | 46.31 | 45.32 | 68.16 | 68.16 |
| *D. lacicola* | 17 | 46.66 | 2.46 | 0.60 | 46.66 | 47.12 | 51.37 | 51.37 |
| *Z. indianus* | 606 | 21.28 | 11.55 | 0.47 | 21.23 | 23.11 | 46.58 | 42.07 |
| *Z. camerounensis* | 5 | 7.31 | 3.55 | 1.59 | 7.31 | 6.20 | 13.58 | 13.58 |
| *D. repletoides* | 11 | 29.47 | 4.88 | 1.47 | 29.47 | 29.78 | 35.77 | 35.77 |
| *D. immigrans* | 1683 | 36.28 | 11.20 | 0.27 | 36.49 | 36.42 | 62.91 | 53.38 |
| *D. hypocausta* | 54 | 8.62 | 5.17 | 0.70 | 8.62 | 7.14 | 24.00 | 19.31 |
| *D. albomicans* | 197 | 24.87 | 6.59 | 0.47 | 25.01 | 24.20 | 35.83 | 34.69 |
| *D. sulfurigaster blilimbata* | 98 | 17.29 | 8.77 | 0.89 | 17.31 | 17.05 | 32.41 | 30.30 |
| *D. tripunctata* | 154 | 34.45 | 7.47 | 0.60 | 34.71 | 35.42 | 46.45 | 42.50 |
| *D. polymorpha* | 266 | 18.45 | 9.31 | 0.57 | 18.48 | 21.59 | 34.60 | 31.49 |
| *D. cardini* | 353 | 17.66 | 8.44 | 0.45 | 17.68 | 18.40 | 34.55 | 31.49 |
| *D. funebris* | 1387 | 46.79 | 8.41 | 0.23 | 46.87 | 47.00 | 78.28 | 63.53 |
| *D. multispina* | 20 | 42.38 | 4.03 | 0.90 | 42.38 | 43.00 | 45.15 | 45.15 |
| *D. macrospina* | 131 | 33.82 | 4.49 | 0.39 | 33.79 | 32.41 | 47.12 | 43.03 |
| *D. putrida* | 232 | 37.24 | 4.79 | 0.32 | 37.25 | 37.33 | 46.45 | 45.14 |
| *D. testacea* | 698 | 47.44 | 5.67 | 0.22 | 47.37 | 47.05 | 69.49 | 60.25 |
| *D. brachynephros* | 169 | 38.38 | 4.16 | 0.32 | 38.34 | 37.00 | 51.12 | 45.45 |
| *D. deflecta* | 5 | 38.15 | 5.52 | 2.47 | 38.15 | 40.00 | 43.10 | 43.10 |
| *D. recens* | 48 | 45.26 | 4.61 | 0.67 | 45.26 | 44.14 | 54.50 | 53.83 |
| *D. transversa* | 888 | 51.11 | 6.68 | 0.22 | 51.10 | 50.44 | 69.95 | 65.03 |

**Supplementary file 2B** Coordinates (decimal degrees) representing the original geographic collection sites of the fruit fly strains tested in this study.

| **Species** | **Latitude** | **Longitude** |  | **Species** | **Latitude** | **Longitude** |
| --- | --- | --- | --- | --- | --- | --- |
| *S. latifasciaeformis* | 9.16 | -79.85 |  | *D. virilis* (Japan) | 38.00 | 140.00 |
| *S. lebanonensis* | 33.86 | 33.57 |  | *D. lummei* | 65.95 | 29.17 |
| *S. rufifrons* | 43.58 | 39.72 |  | *D. novamexicana* | 33.92 | -106.87 |
| *H. duncani* | 30.43 | -83.92 |  | *D. americana texana* | 36.47 | -89.32 |
| *D. paulistorum* | -5.00 | -63.00 |  | *D. borealis* (eastern) | 44.10 | -88.90 |
| *D. nebulosa* | 22.77 | -105.67 |  | *D. lacicola* | 51.38 | -96.90 |
| *D. prosaltans* | -24.52 | -48.25 |  | *Z. indianus* | 31.22 | 29.97 |
| *D. sturtevanti* | 10.00 | -84.00 |  | *Z. camerounensis* | -5.08 | 38.67 |
| *D. melanogaster* (Tanzania) | -3.18 | 37.23 |  | *D. repletoides* | 35.77 | 139.10 |
| *D. kikkawai* | 25.02 | 121.59 |  | *D. immigrans* | 49.58 | 10.06 |
| *D. ananassae* | 6.01 | 10.02 |  | *D. hypocausta* | 13.47 | 144.77 |
| *D. obscura* | 65.02 | 25.47 |  | *D. albomicans* | 24.33 | 123.82 |
| *D. bifasciata* | 49.58 | 10.06 |  | *D. sulfurigaster blilimbata* | 13.47 | 144.77 |
| *D. subobscura* (BY) | 49.78 | 9.98 |  | *D. tripunctata* | 33.90 | -83.38 |
| *D. subobscura* (BW) | 49.38 | 8.72 |  | *D. polymorpha* | -27.60 | -48.48 |
| *D. azteca* | 17.17 | -96.72 |  | *D. cardini* | 28.00 | -82.00 |
| *D. busckii* | 49.58 | 10.06 |  | *D. funebris* (Alaska) | 61.52 | -149.90 |
| *S. pallida* (Bavaria 2017) | 49.58 | 10.06 |  | *D. funebris* (Bavaria Q) | 49.59 | 10.00 |
| *S. pallida* (Bavaria 2020) | 49.58 | 10.06 |  | *D. funebris* (Bavaria V) | 49.59 | 10.00 |
| *D. robusta* | 42.00 | -91.97 |  | *D. funebris* (Hokkaido) | 43.43 | 142.72 |
| *D. tsigana* | 43.43 | 142.72 |  | *D. funebris* (Lebanon) | 33.78 | 35.55 |
| *D. melanica* | 30.38 | -97.77 |  | *D. funebris* (Mexico) | 19.58 | -99.12 |
| *D. gaucha* | -36.08 | -60.09 |  | *D. multispina* | 43.43 | 142.72 |
| *D. hydei* | -3.18 | 37.23 |  | *D. macrospina* (South Carolina) | 34.65 | -81.87 |
| *D. repleta* (Bavaria K) | 49.58 | 10.06 |  | *D. macrospina* (Arizona) | 31.48 | -110.85 |
| *D. repleta* (Bavaria S) | 49.59 | 10.00 |  | *D. putrida* | 41.00 | -99.00 |
| *D. mercatorum* | -3.18 | 37.23 |  | *D. testacea* | 46.51 | 6.56 |
| *D. kanekoi* | 43.52 | 143.67 |  | *D. brachynephros* | 33.00 | 133.00 |
| *D. ezoana* | 66.47 | 29.04 |  | *D. deflecta* | 40.36 | -74.67 |
| *D. littoralis* | 69.04 | 20.80 |  | *D. recens* | 43.00 | -74.00 |
| *D. virilis* (Rwanda) | 2.00 | 29.00 |  | *D. transversa* | 59.80 | 17.70 |
