## Supplementary file 3_M-,E-peaks for "Fruit flies of the subgenera *Drosophila*, *Siphlodora* and *Sophophora* use different mechanisms to adapt their daily activity to long photoperiods"

### Peak determination of different fruit fly species in LD 12:12, LD 16:8, and LD 20:4

**Supplementary file 3A** Determination of activity peaks in different fruit fly species under LD 12:12. N is the number of individual flies used to generate the average daily activity profile from which the peaks were determined. Selected morning (M) and evening (E) peaks (in bold) reflect the circadian-driven peaks and indicate the values used to calculate ΔΨ_M-E_ for this photoperiod. Additional peaks found in the daily activity profile are noted with comments.

| **Species (strain)** | N | Selected morning peak (M) | Selected evening peak (E) | ΔΨ_M-E_ in LD 12:12 | Additional peak (ZT) | comment |
| --- | --- | --- | --- | --- | --- | --- |
| *S. latifasciaeformis* | 86 | 0.32 | 12.04 | 11.72 |  |  |
| *S. lebanonensis* | 63 | 4.84 | 10.13 | 5.30 |  |  |
| *S. rufifrons* | 64 | 0.34 | 10.56 | 10.22 |  |  |
| *H. duncani* | 71 | 2.90 | 11.56 | 8.65 |  |  |
| *D. paulistorum* | 107 | 0.40 | 8.25 | 7.86 |  |  |
| *D. nebulosa* | 96 | 0.50 | 11.42 | 10.92 |  |  |
| *D. prosaltans* | 63 | 1.27 | 12.02 | 10.75 |  |  |
| *D. sturtevanti* | 89 | 0.41 | 12.05 | 11.64 |  |  |
| *D. melanogaster* (Tanzania) | 80 | 0.27 | 11.90 | 11.62 |  |  |
| *D. kikkawai* | 62 | 0.52 | 11.91 | 11.39 |  |  |
| *D. ananassae* | 63 | 1.57 | 11.99 | 10.41 |  |  |
| *D. obscura* | 75 | 0.53 | 11.56 | 11.03 |  |  |
| *D. bifasciata* | 127 | 0.42 | 11.77 | 11.36 |  |  |
| *D. subobscura* (BY) | 63 | 0.58 | 10.01 | 9.43 |  |  |
| *D. subobscura* (BW) | 76 | 0.65 | 10.15 | 9.50 |  |  |
| *D. azteca* | 63 | 0.34 | 11.94 | 11.60 |  |  |
| *D. busckii* | 137 | 0.31 | 10.87 | 10.56 | 20.14 | night peak |
| *S. pallida* (2017) | 96 | 1.00 | 11.40 | 10.40 | 19.09 | night peak |
| *S. pallida* (2020) | 60 | 0.46 | 11.70 | 11.23 | 19.73 | night peak |
| *D. robusta* | 62 | 0.48 | 12.20 | 11.72 | 19.21 | night peak |
| *D. tsigana* | 81 | 0.36 | 11.77 | 11.41 |  |  |
| *D. melanica* | 124 | 0.49 | 10.96 | 10.47 |  |  |
| *D. gaucha* | 77 | 0.36 | 11.65 | 11.29 |  |  |
| *D. hydei* | 63 | 0.41 | 11.88 | 11.48 |  |  |
| *D. repleta* (strain K) | 64 | 0.76 | 11.53 | 10.77 |  |  |
| *D. repleta* (strain S) | 102 | 0.93 | 10.70 | 9.77 |  |  |
| *D. mercatorum* | 62 | 0.29 | 11.49 | 11.21 |  |  |
| *D. kanekoi* | 93 | 1.19 | 11.79 | 10.61 |  |  |
| *D. ezoana* | 60 | 0.60 | 11.90 | 11.30 |  |  |
| *D. littoralis* | 64 | 0.83 | 11.74 | 10.91 |  |  |
| *D. virilis* (Rwanda) | 159 | 1.08 | 11.59 | 10.51 |  |  |
| *D. virilis* (Japan) | 62 | 1.10 | 10.77 | 9.67 |  |  |
| *D. lummei* | 63 | 0.35 | 11.94 | 11.60 |  |  |
| *D. novamexicana* | 153 | 0.44 | 11.52 | 11.08 |  |  |
| *D. americana texana* | 152 | 1.13 | 11.19 | 10.06 |  |  |
| *D. borealis* (eastern) | 95 | -0.37 | 11.94 | 12.31 |  |  |
| *D. lacicola* | 160 | 0.38 | 11.79 | 11.41 |  |  |
| *Z. indianus* | 111 | 0.29 | 11.87 | 11.58 |  |  |
| *Z. camerounensis* | 64 | 0.39 | 11.80 | 11.41 |  |  |
| *D. repletoides* | 61 | 0.65 | 10.63 | 9.98 |  |  |
| *D. immigrans* | 80 | 0.16 | 11.86 | 11.70 |  |  |
| *D. hypocausta* | 125 | 2.92 | 11.61 | 8.69 |  |  |
| *D. albomicans* | 63 | 1.00 | 11.61 | 10.61 |  |  |
| *D. sulfurigaster blilimbata* | 108 | 0.34 | 11.71 | 11.37 |  |  |
| *D. tripunctata* | 104 | 0.32 | 11.27 | 10.95 |  |  |
| *D. polymorpha* | 68 | 0.42 | 11.54 | 11.12 |  |  |
| *D. cardini* | 97 | 0.39 | 11.24 | 10.85 |  |  |
| *D. funebris* (Alaska) | 185 | 0.29 | 11.93 | 11.64 |  |  |
| *D. funebris* (Bavaria Q) | 80 | 0.41 | 11.85 | 11.44 |  |  |
| *D. funebris* (Bavaria V) | 100 | 0.37 | 10.64 | 10.27 |  |  |
| *D. funebris* (Hokkaido) | 105 | 1.13 | 11.84 | 10.71 |  |  |
| *D. funebris* (Lebanon) | 85 | 0.46 | 11.66 | 11.20 |  |  |
| *D. funebris* (Mexico) | 119 | 1.24 | 10.12 | 8.89 |  |  |
| *D. multispina* | 132 | 0.49 | 11.97 | 11.48 |  |  |
| *D. macrospina* (SC) | 92 | 0.28 | 11.77 | 11.49 |  |  |
| *D. macrospina* (AZ) | 68 | 1.21 | 11.43 | 10.22 |  |  |
| *D. putrida* | 190 | 0.51 | 6.85 | 6.34 | 6.85 | “afternoon peak” |
| *D. testacea* | 59 | 0.38 | 11.41 | 11.04 |  |  |
| *D. brachynephros* | 93 | 0.65 | 11.14 | 10.49 |  |  |
| *D. deflecta* | 78 | 0.92 | 10.80 | 9.88 |  |  |
| *D. recens* | 99 | 0.34 | 11.79 | 11.45 |  |  |
| *D. transversa* | 74 | 1.05 | 11.94 | 10.88 |  |  |

**Supplementary file 3B** Determination of activity peaks in different fruit fly species under LD 16:8. N is the number of individual flies used to generate the average daily activity profile from which the peaks were determined. Selected morning (M) and evening (E) peaks (in bold) reflect the circadian-driven peaks and indicate the values used to calculate ΔΨM-E for this photoperiod. Additional peaks found in the daily activity profile are noted with comments.

| **Species (strain)** | N | Selected morning peak (M) | Selected evening peak (E) | ΔΨ_M-E_ in LD 16:8 | Additional peak (ZT) | comment |
| --- | --- | --- | --- | --- | --- | --- |
| *S. latifasciaeformis* | 106 | 0.48 | 15.88 | 15.39 |  |  |
| *S. lebanonensis* | 95 | 5.55 | 12.81 | 7.26 |  |  |
| *S. rufifrons* | 108 | 1.51 | 14.60 | 13.09 |  |  |
| *H. duncani* | 100 | 0.40 | 15.68 | 15.28 |  |  |
| *D. paulistorum* | 78 | 0.43 | 11.70 | 11.27 |  |  |
| *D. nebulosa* | 85 | 5.16 | 12.91 | 7.76 |  |  |
| *D. prosaltans* | 74 | 0.66 | 12.89 | 12.23 |  |  |
| *D. sturtevanti* | 80 | 0.40 | 14.34 | 13.94 |  |  |
| *D. melanogaster* (Tanzania) | 94 | 0.27 | 14.15 | 13.89 |  |  |
| *D. kikkawai* | 120 | 0.43 | 13.47 | 13.05 |  |  |
| *D. ananassae* | 105 | 0.41 | 15.63 | 15.22 |  |  |
| *D. obscura* | 117 | 0.44 | 15.70 | 15.26 |  |  |
| *D. bifasciata* | 113 | 0.41 | 14.83 | 14.42 |  |  |
| *D. subobscura* (BY) | 96 | 0.63 | 13.37 | 12.74 |  |  |
| *D. subobscura* (BW) | 80 | 0.65 | 12.78 | 12.13 |  |  |
| *D. azteca* | 98 | 1.34 | 15.87 | 14.54 |  |  |
| *D. busckii* | 90 | 0.37 | 13.51 | 13.14 | 22.66 | advance peak |
| *S. pallida* (2017) | 165 | 0.65 | 14.39 | 13.74 |  |  |
| *S. pallida* (2020) | 80 | 0.90 | 14.25 | 13.35 | 20.25 |  |
| *D. robusta* | 81 | 0.47 | 16.13 | 15.67 | 20.81 |  |
| *D. tsigana* | 91 | 0.65 | 14.56 | 13.91 |  |  |
| *D. melanica* | 80 | 0.52 | 13.70 | 13.18 |  |  |
| *D. gaucha* | 88 | 2.42 | 14.18 | 11.77 |  |  |
| *D. hydei* | 79 | 0.66 | 14.72 | 14.06 |  |  |
| *D. repleta* (strain K) | 63 | 0.83 | 12.78 | 11.95 |  |  |
| *D. repleta* (strain S) | 76 | 1.06 | 12.28 | 11.22 | 6.01 |  |
| *D. mercatorum* | 80 | 4.69 | 14.25 | 9.56 |  |  |
| *D. kanekoi* | 79 | 0.25 | 14.68 | 14.43 |  |  |
| *D. ezoana* | 91 | 0.36 | 15.73 | 15.37 |  |  |
| *D. littoralis* | 77 | 0.80 | 15.52 | 14.72 |  |  |
| *D. virilis* (Rwanda) | 98 | 1.14 | 14.32 | 13.18 | 8.94 |  |
| *D. virilis* (Japan) | 74 | 1.06 | 13.82 | 12.76 |  |  |
| *D. lummei* | 63 | 0.46 | 15.67 | 15.21 |  |  |
| *D. novamexicana* | 95 | 0.50 | 14.00 | 13.50 |  |  |
| *D. americana texana* | 92 | 1.04 | 13.62 | 12.58 |  |  |
| *D. borealis* (eastern) | 63 | 1.00 | 14.18 | 13.18 |  |  |
| *D. lacicola* | 101 | 0.75 | 13.66 | 12.91 |  |  |
| *Z. indianus* | 77 | 0.65 | 14.28 | 13.63 |  |  |
| *Z. camerounensis* | 87 | 4.23 | 15.63 | 11.40 |  |  |
| *D. repletoides* | 80 | 0.79 | 13.06 | 12.28 |  |  |
| *D. immigrans* | 95 | 0.74 | 14.40 | 13.65 |  |  |
| *D. hypocausta* | 104 | 1.28 | 15.61 | 14.33 | 4.03 |  |
| *D. albomicans* | 85 | 0.67 | 15.42 | 14.75 |  |  |
| *D. sulfurigaster blilimbata* | 89 | 0.55 | 13.91 | 13.36 |  |  |
| *D. tripunctata* | 110 | 1.08 | 15.05 | 13.97 |  |  |
| *D. polymorpha* | 86 | 4.36 | 15.27 | 10.91 |  |  |
| *D. cardini* | 88 | 0.63 | 12.89 | 12.27 |  |  |
| *D. funebris* (Alaska) | 101 | 0.36 | 14.03 | 13.68 |  |  |
| *D. funebris* (Bavaria Q) | 111 | 0.26 | 15.32 | 15.06 |  |  |
| *D. funebris* (Bavaria V) | 104 | 0.37 | 12.55 | 12.17 |  |  |
| *D. funebris* (Hokkaido) | 78 | 0.48 | 14.24 | 13.76 |  |  |
| *D. funebris* (Lebanon) | 92 | 1.59 | 13.80 | 12.21 |  |  |
| *D. funebris* (Mexico) | 96 | 1.91 | 13.93 | 12.02 |  |  |
| *D. multispina* | 93 | 0.69 | 14.12 | 13.42 |  |  |
| *D. macrospina* (SC) | 96 | 1.62 | 14.76 | 13.14 |  |  |
| *D. macrospina* (AZ) | 99 | 1.12 | 14.39 | 13.27 |  |  |
| *D. putrida* | 143 | 0.70 | 15.76 | 15.06 |  |  |
| *D. testacea* | 96 | 0.41 | 15.70 | 15.29 |  |  |
| *D. brachynephros* | 84 | 0.68 | 13.38 | 12.70 |  |  |
| *D. deflecta* | 80 | 2.95 | 12.87 | 9.92 |  |  |
| *D. recens* | 83 | 0.37 | 14.05 | 13.67 |  |  |
| *D. transversa* | 100 | 1.10 | 13.44 | 12.34 |  |  |

**Supplementary file 3C** Determination of activity peaks in different fruit fly species under LD 20:4. N is the number of individual flies used to generate the average daily activity profile from which the peaks were determined. Selected morning (M) and evening (E) peaks (in bold) reflect the circadian-driven peaks and indicate the values used to calculate ΔΨM-E for this photoperiod. Asterisks (*) mark species for which individuals that did not synchronize to LD 20:4 were excluded from peak determination; peaks from the remaining synchronized individuals were selected for further analysis. Additional peaks found in the daily activity profile are noted with comments.

| **Species (strain)** | N | Selected morning peak (M) | Selected evening peak (E) | ΔΨ_M-E_ in LD 20:4 | Additional peak (ZT) | comment |
| --- | --- | --- | --- | --- | --- | --- |
| *S. latifasciaeformis* | 101 | 4.76 | 14.40 | 9.63 |  |  |
| *S. lebanonensis* | 95 | 4.58 | 16.05 | 11.47 |  |  |
| *S. rufifrons* | 106 | 3.76 | 13.84 | 10.08 |  |  |
| *H. duncani* | 85 | 4.02 | 16.50 | 12.48 |  |  |
| *D. paulistorum* | 77 | 0.52 | 10.51 | 9.99 |  |  |
| *D. nebulosa* | 76 | 0.56 | 11.74 | 11.18 |  |  |
| *D. prosaltans* | 70 | 2.35 | 13.15 | 10.81 |  |  |
| *D. sturtevanti* | 66 | 3.76 | 14.89 | 11.12 |  |  |
| *D. melanogaster* (Tanzania) | 92 | 0.33 | 13.68 | 13.35 |  |  |
| *D. kikkawai** | 38 | 4.26 | 14.78 | 10.53 |  | selected |
| *D. ananassae* | 101 | 3.92 | 14.02 | 10.10 |  |  |
| *D. obscura* | 112 | 0.52 | 17.97 | 17.45 |  |  |
| *D. bifasciata* | 111 | 0.55 | 16.93 | 16.38 |  |  |
| *D. subobscura* (BY) | 96 | 0.61 | 16.06 | 15.44 |  |  |
| *D. subobscura* (BW) | 79 | 0.81 | 14.90 | 14.10 |  |  |
| *D. azteca* | 97 | 4.28 | 16.88 | 12.60 |  |  |
| *D. busckii* | 88 | 0.67 | 15.84 | 15.17 |  |  |
| *S. pallida* (2017) | 85 | 1.22 | 16.46 | 15.24 |  |  |
| *S. pallida* (2020) | 79 | 1.53 | 17.90 | 16.37 | 7.05 |  |
| *D. robusta* | 81 | 0.73 | 15.65 | 14.92 |  |  |
| *D. tsigana* | 90 | 3.90 | 15.91 | 12.01 |  |  |
| *D. melanica* | 80 | 2.92 | 15.06 | 12.14 |  |  |
| *D. gaucha* | 86 | 4.08 | 15.65 | 11.58 |  |  |
| *D. hydei* | 79 | 4.57 | 15.57 | 11.00 |  |  |
| *D. repleta* (strain K) | 57 | 1.39 | 12.53 | 11.14 |  |  |
| *D. repleta* (strain S) | 74 | 1.33 | 12.04 | 10.71 |  |  |
| *D. mercatorum* | 79 | 6.87 | 16.13 | 9.26 |  |  |
| *D. kanekoi** | 16 | 2.62 | 16.34 | 13.72 |  | selected |
| *D. ezoana* | 91 | 3.50 | 18.65 | 15.15 |  |  |
| *D. littoralis* | 77 | 0.49 | 15.76 | 15.27 |  |  |
| *D. virilis* (Rwanda) | 98 | 1.38 | 15.44 | 14.07 |  |  |
| *D. virilis* (Japan) | 73 | 1.25 | 17.07 | 15.82 |  |  |
| *D. lummei* | 75 | 0.62 | 17.10 | 16.48 |  |  |
| *D. novamexicana* | 92 | 3.47 | 15.89 | 12.42 |  |  |
| *D. americana texana* | 86 | 1.18 | 16.34 | 15.16 |  |  |
| *D. borealis* (eastern) | 63 | 1.20 | 16.74 | 15.54 |  |  |
| *D. lacicola* | 99 | 0.77 | 15.99 | 15.22 |  |  |
| *Z. indianus* | 77 | 4.01 | 15.38 | 11.36 |  |  |
| *Z. camerounensis* | 85 | 5.37 | 15.97 | 10.59 |  |  |
| *D. repletoides* | 80 | 0.35 | 14.58 | 14.24 |  |  |
| *D. immigrans* | 87 | 1.03 | 15.92 | 14.89 |  |  |
| *D. hypocausta* | 83 | 3.69 | 13.12 | 9.43 |  |  |
| *D. albomicans* | 84 | 3.54 | 14.69 | 11.15 |  |  |
| *D. sulfurigaster blilimbata** | 38 | 0.93 | 12.15 | 11.22 |  | selected |
| *D. tripunctata* | 98 | 1.06 | 15.34 | 14.28 |  |  |
| *D. polymorpha* | 78 | 4.48 | 15.00 | 10.52 |  |  |
| *D. cardini* | 84 | 4.13 | 14.21 | 10.08 |  |  |
| *D. funebris* (Alaska) | 98 | 0.56 | 16.73 | 16.16 |  |  |
| *D. funebris* (Bavaria Q) | 110 | 0.39 | 16.46 | 16.07 |  |  |
| *D. funebris* (Bavaria V) | 101 | 0.34 | 16.06 | 15.71 |  |  |
| *D. funebris* (Hokkaido) | 78 | 0.46 | 17.82 | 17.35 |  |  |
| *D. funebris* (Lebanon) | 89 | 5.04 | 16.73 | 11.68 |  |  |
| *D. funebris* (Mexico) | 92 | 4.40 | 17.02 | 12.63 |  |  |
| *D. multispina* | 89 | 0.84 | 16.33 | 15.49 |  |  |
| *D. macrospina* (SC) | 93 | 3.95 | 16.00 | 12.05 |  |  |
| *D. macrospina* (AZ) | 94 | 1.26 | 16.60 | 15.34 |  |  |
| *D. putrida* | 78 | 3.63 | 14.01 | 10.38 |  |  |
| *D. testacea* | 82 | 0.88 | 16.80 | 15.92 |  |  |
| *D. brachynephros* | 80 | 0.69 | 14.23 | 13.54 |  |  |
| *D. deflecta** | 25 | 6.85 | 14.32 | 7.47 |  | selected |
| *D. recens* | 81 | 0.60 | 15.34 | 14.74 |  |  |
| *D. transversa* | 89 | 1.28 | 16.07 | 14.79 |  |  |
