## Supplementary file 4_DD_statistics_Lomb-Scargle for "Fruit flies of the subgenera *Drosophila*, *Siphlodora* and *Sophophora* use different mechanisms to adapt their daily activity to long photoperiods"

### Descriptive statistics for constant darkness (DD) behavior of the different drosophilid species

**Supplementary file 4** DD rhythmicity values and Lomb-Scargle (LS) analysis for the different fruit fly species and strains. Not all flies scored as rhythmic showed a significant free-running period in the LS analysis, for the reasons explained in the Materials and Methods section. SD, standard deviation; SE, standard error.

| Species (strain) | N (total flies analyzed) | Rhythmicity clear (%) | Rhythmicity complex (%) | **Total rhythmicity (%)** | **Arrhythmic (%)** | Flies with significant major period | Mean major period (h) | Major period SD (h) | Major period SE (h) | Mean major power | Major power SD | Major power SE | Flies with significant minor period | Mean minor period (h) | Minor period SD (h) | Minor SE (h) | Mean minor power | Minor power SD | Minor power SE |
| --- | --- | --- | --- | --- | --- | --- | --- | --- | --- | --- | --- | --- | --- | --- | --- | --- | --- | --- | --- |
| *S. latifasciaeformis* | 49 | 61.2 | 38.8 | **100.0** | **0.0** | 49 | 23.53 | 0.81 | 0.12 | 118.8 | 107.8 | 15.4 | 0 | na | na | na | na | na | na |
| *S. lebanonensis* | 32 | 93.8 | 6.3 | **100.0** | **0.0** | 32 | 24.02 | 0.28 | 0.05 | 291.3 | 152.4 | 26.9 | 0 | na | na | na | na | na | na |
| *S. rufifrons* | 40 | 97.5 | 2.5 | **100.0** | **0.0** | 40 | 24.24 | 0.37 | 0.06 | 127.1 | 70.6 | 11.2 | 0 | na | na | na | na | na | na |
| *H. duncani* | 34 | 97.1 | 2.9 | **100.0** | **0.0** | 34 | 23.63 | 0.34 | 0.06 | 128.3 | 104.8 | 18.0 | 18 | 21.24 | 0.29 | 0.05 | 27.9 | 14.7 | 2.5 |
| *D. paulistorum* | 67 | 22.4 | 59.7 | **82.1** | **17.9** | 40 | 24.17 | 0.87 | 0.11 | 33.6 | 26.2 | 3.2 | 0 | na | na | na | na | na | na |
| *D. nebulosa* | 32 | 37.5 | 46.9 | **84.4** | **15.6** | 21 | 24.18 | 0.72 | 0.13 | 21.2 | 8.1 | 1.4 | 0 | na | na | na | na | na | na |
| *D. prosaltans* | 30 | 70.0 | 30.0 | **100.0** | **0.0** | 30 | 24.77 | 0.52 | 0.09 | 43.7 | 20.0 | 3.7 | 0 | na | na | na | na | na | na |
| *D. sturtevanti* | 46 | 73.9 | 26.1 | **100.0** | **0.0** | 46 | 23.43 | 0.53 | 0.08 | 193.0 | 176.0 | 25.9 | 13 | 26.52 | 0.68 | 0.10 | 42.0 | 29.1 | 4.3 |
| *D. melanogaster* (Tanzania) | 47 | 80.9 | 19.1 | **100.0** | **0.0** | 47 | 23.75 | 0.76 | 0.11 | 269.9 | 262.1 | 38.2 | 0 | na | na | na | na | na | na |
| *D. kikkawai* | 30 | 70.0 | 30.0 | **100.0** | **0.0** | 28 | 24.42 | 0.96 | 0.18 | 51.4 | 34.4 | 6.3 | 0 | na | na | na | na | na | na |
| *D. ananassae* | 31 | 93.5 | 6.5 | **100.0** | **0.0** | 31 | 23.92 | 0.54 | 0.10 | 114.2 | 74.4 | 13.4 | 0 | na | na | na | na | na | na |
| *D. obscura* | 45 | 64.4 | 31.1 | **95.6** | **4.4** | 38 | 24.75 | 0.48 | 0.07 | 54.7 | 49.6 | 7.4 | 0 | na | na | na | na | na | na |
| *D. bifasciata* | 48 | 29.2 | 68.8 | **97.9** | **2.1** | 47 | 21.32 | 0.37 | 0.05 | 199.1 | 205.0 | 29.6 | 41 | 24.23 | 0.81 | 0.12 | 74.0 | 55.9 | 8.1 |
| *D. subobscura* (BY) | 31 | 83.9 | 16.1 | **100.0** | **0.0** | 31 | 24.64 | 0.71 | 0.13 | 123.9 | 133.1 | 23.9 | 0 | na | na | na | na | na | na |
| *D. subobscura* (BW) | 42 | 11.9 | 81.0 | **92.9** | **7.1** | 30 | 26.27 | 1.10 | 0.17 | 36.4 | 31.1 | 4.8 | 20 | 23.47 | 0.46 | 0.07 | 40.6 | 25.7 | 4.0 |
| *D. azteca* | 37 | 32.4 | 62.2 | **94.6** | **5.4** | 35 | 24.18 | 1.28 | 0.21 | 104.1 | 108.5 | 17.8 | 12 | 23.77 | 2.01 | 0.33 | 40.0 | 22.0 | 3.6 |
| *D. busckii* | 51 | 47.1 | 49.0 | **96.1** | **3.9** | 49 | 23.69 | 0.81 | 0.11 | 215.9 | 213.4 | 29.9 | 17 | 21.55 | 0.40 | 0.06 | 73.4 | 77.8 | 10.9 |
| *S. pallida* (2017) | 75 | 57.3 | 36.0 | **93.3** | **6.7** | 70 | 21.63 | 0.60 | 0.07 | 58.6 | 49.6 | 5.7 | 0 | na | na | na | na | na | na |
| *S. pallida* (2020) | 31 | 19.4 | 77.4 | **96.8** | **3.2** | 29 | 24.19 | 1.19 | 0.21 | 159.3 | 139.4 | 25.0 | 0 | na | na | na | na | na | na |
| *D. robusta* | 38 | 0.0 | 36.8 | **36.8** | **63.2** | 14 | 23.37 | 2.49 | 0.40 | 20.1 | 5.5 | 0.9 | 0 | na | na | na | na | na | na |
| *D. tsigana* | 41 | 87.8 | 12.2 | **100.0** | **0.0** | 38 | 23.67 | 0.28 | 0.04 | 59.0 | 44.9 | 7.0 | 0 | na | na | na | na | na | na |
| *D. melanica* | 40 | 60.0 | 37.5 | **97.5** | **2.5** | 37 | 23.94 | 0.46 | 0.07 | 50.3 | 44.1 | 7.0 | 0 | na | na | na | na | na | na |
| *D. gaucha* | 37 | 45.9 | 45.9 | **91.9** | **8.1** | 32 | 26.22 | 1.00 | 0.16 | 44.2 | 35.6 | 5.8 | 9 | 21.69 | 1.08 | 0.18 | 20.5 | 8.0 | 1.3 |
| *D. hydei* | 31 | 100.0 | 0.0 | **100.0** | **0.0** | 31 | 23.96 | 0.27 | 0.05 | 438.3 | 260.4 | 46.8 | 0 | na | na | na | na | na | na |
| *D. repleta* (strain K) | 30 | 100.0 | 0.0 | **100.0** | **0.0** | 30 | 24.07 | 0.16 | 0.03 | 185.5 | 82.4 | 15.0 | 0 | na | na | na | na | na | na |
| *D. repleta* (strain S) | 36 | 82.4 | 17.6 | **100.0** | **0.0** | 33 | 24.17 | 0.46 | 0.08 | 138.8 | 184.9 | 30.8 | 0 | na | na | na | na | na | na |
| *D. mercatorum* | 31 | 100.0 | 0.0 | **100.0** | **0.0** | 31 | 24.47 | 0.28 | 0.05 | 147.9 | 84.6 | 15.2 | 0 | na | na | na | na | na | na |
| *D. kanekoi* | 56 | 3.6 | 71.4 | **75.0** | **25.0** | 39 | 23.98 | 1.86 | 0.25 | 23.6 | 9.2 | 1.2 | 0 | na | na | na | na | na | na |
| *D. ezoana* | 37 | 0.0 | 62.2 | **62.2** | **37.8** | 21 | 24.19 | 2.00 | 0.33 | 25.2 | 10.0 | 1.6 | 0 | na | na | na | na | na | na |
| *D. littoralis* | 31 | 0.0 | 32.3 | **32.3** | **67.7** | 10 | 23.65 | 2.30 | 0.41 | 20.2 | 6.7 | 1.2 | 0 | na | na | na | na | na | na |
| *D. virilis* (Rwanda) | 63 | 50.8 | 41.3 | **92.1** | **7.9** | 57 | 22.58 | 1.04 | 0.13 | 48.1 | 32.8 | 4.1 | 0 | na | na | na | na | na | na |
| *D. virilis* (Japan) | 31 | 61.3 | 38.7 | **100.0** | **0.0** | 31 | 24.48 | 0.46 | 0.08 | 108.3 | 59.3 | 10.7 | 18 | 21.84 | 0.56 | 0.10 | 34.0 | 23.9 | 4.3 |
| *D. lummei* | 32 | 0.0 | 62.5 | **62.5** | **37.5** | 20 | 23.77 | 2.31 | 0.41 | 28.7 | 14.8 | 2.6 | 0 | na | na | na | na | na | na |
| *D. novamexicana* | 57 | 12.3 | 63.2 | **75.4** | **24.6** | 32 | 24.38 | 2.33 | 0.31 | 29.2 | 23.2 | 3.1 | 0 | na | na | na | na | na | na |
| *D. americana texana* | 58 | 12.1 | 58.6 | **70.7** | **29.3** | 40 | 23.54 | 1.12 | 0.15 | 32.6 | 17.8 | 2.3 | 0 | na | na | na | na | na | na |
| *D. borealis* (eastern) | 60 | 6.7 | 70.0 | **76.7** | **23.3** | 46 | 22.99 | 1.24 | 0.16 | 25.8 | 10.2 | 1.3 | 0 | na | na | na | na | na | na |
| *D. lacicola* | 62 | 1.6 | 69.4 | **71.0** | **29.0** | 35 | 24.24 | 1.56 | 0.20 | 24.5 | 12.8 | 1.6 | 0 | na | na | na | na | na | na |
| *Z. indianus* | 37 | 29.7 | 62.2 | **91.9** | **8.1** | 29 | 23.72 | 1.33 | 0.22 | 59.1 | 68.2 | 11.2 | 0 | na | na | na | na | na | na |
| *Z. camerounensis* | 31 | 96.8 | 3.2 | **100.0** | **0.0** | 31 | 23.80 | 0.26 | 0.05 | 183.7 | 105.6 | 19.0 | 0 | na | na | na | na | na | na |
| *D. repletoides* | 36 | 100.0 | 0.0 | **100.0** | **0.0** | 36 | 24.60 | 0.25 | 0.04 | 359.1 | 226.3 | 37.7 | 0 | na | na | na | na | na | na |
| *D. immigrans* | 46 | 4.3 | 47.8 | **52.2** | **47.8** | 20 | 24.38 | 1.11 | 0.16 | 34.5 | 29.0 | 4.3 | 0 | na | na | na | na | na | na |
| *D. hypocausta* | 39 | 12.8 | 51.3 | **64.1** | **35.9** | 21 | 23.74 | 2.19 | 0.35 | 24.5 | 9.1 | 1.5 | 0 | na | na | na | na | na | na |
| *D. albomicans* | 32 | 50.0 | 50.0 | **100.0** | **0.0** | 25 | 23.20 | 1.21 | 0.21 | 35.8 | 23.2 | 4.1 | 5 | 24.11 | 2.70 | 0.48 | 16.6 | 3.9 | 0.7 |
| *D. sulfurigaster blilimbata* | 27 | 11.1 | 74.1 | **85.2** | **14.8** | 15 | 25.61 | 2.09 | 0.40 | 20.7 | 5.4 | 1.0 | 0 | na | na | na | na | na | na |
| *D. tripunctata* | 40 | 70.0 | 30.0 | **100.0** | **0.0** | 40 | 23.96 | 0.37 | 0.06 | 104.0 | 76.0 | 12.0 | 0 | na | na | na | na | na | na |
| *D. polymorpha* | 44 | 63.6 | 29.5 | **93.2** | **6.8** | 37 | 24.18 | 0.66 | 0.10 | 53.2 | 42.7 | 6.4 | 0 | na | na | na | na | na | na |
| *D. cardini* | 30 | 46.7 | 50.0 | **96.7** | **3.3** | 27 | 24.71 | 0.53 | 0.10 | 61.1 | 33.1 | 6.0 | 0 | na | na | na | na | na | na |
| *D. funebris*  (Alaska) | 43 | 100.0 | 0.0 | **100.0** | **0.0** | 43 | 24.20 | 0.26 | 0.04 | 257.8 | 147.2 | 22.5 | 0 | na | na | na | na | na | na |
| *D. funebris*  (Bavaria Q) | 48 | 83.3 | 16.7 | **100.0** | **0.0** | 48 | 23.66 | 0.59 | 0.08 | 119.3 | 95.1 | 13.7 | 0 | na | na | na | na | na | na |
| *D. funebris*  (Bavaria V) | 38 | 76.3 | 21.1 | **97.4** | **2.6** | 37 | 23.78 | 0.90 | 0.15 | 117.8 | 79.6 | 12.9 | 16 | 21.20 | 0.34 | 0.06 | 23.8 | 8.3 | 1.4 |
| *D. funebris*  (Hokkaido) | 68 | 100.0 | 0.0 | **100.0** | **0.0** | 68 | 25.16 | 0.25 | 0.03 | 450.7 | 244.3 | 29.6 | 58 | 21.82 | 0.70 | 0.09 | 35.0 | 19.0 | 2.3 |
| *D. funebris*  (Lebanon) | 31 | 100.0 | 0.0 | **100.0** | **0.0** | 31 | 23.15 | 0.29 | 0.05 | 123.6 | 68.4 | 12.3 | 0 | na | na | na | na | na | na |
| *D. funebris*  (Mexico) | 55 | 98.2 | 1.8 | **100.0** | **0.0** | 55 | 23.22 | 0.24 | 0.03 | 248.2 | 167.6 | 22.6 | 0 | na | na | na | na | na | na |
| *D. multispina* | 30 | 76.7 | 23.3 | **100.0** | **0.0** | 29 | 24.57 | 0.38 | 0.07 | 123.9 | 87.1 | 15.9 | 0 | na | na | na | na | na | na |
| *D. macrospina* (SC) | 36 | 88.9 | 11.1 | **100.0** | **0.0** | 36 | 22.58 | 0.74 | 0.12 | 122.7 | 71.0 | 11.8 | 0 | na | na | na | na | na | na |
| *D. macrospina* (AZ) | 42 | 64.3 | 28.6 | **92.9** | **7.1** | 37 | 21.97 | 0.38 | 0.06 | 59.3 | 35.6 | 5.5 | 0 | na | na | na | na | na | na |
| *D. putrida* | 50 | 30.0 | 60.0 | **90.0** | **10.0** | 44 | 24.16 | 1.08 | 0.15 | 98.1 | 81.7 | 11.6 | 0 | na | na | na | na | na | na |
| *D. testacea* | 46 | 37.0 | 60.9 | **97.8** | **2.2** | 43 | 22.88 | 0.90 | 0.13 | 50.2 | 33.1 | 4.9 | 0 | na | na | na | na | na | na |
| *D. brachynephros* | 60 | 5.0 | 66.7 | **71.7** | **28.3** | 42 | 23.73 | 1.39 | 0.18 | 27.1 | 14.9 | 1.9 | 0 | na | na | na | na | na | na |
| *D. deflecta* | 39 | 10.3 | 74.4 | **84.6** | **15.4** | 33 | 24.73 | 1.16 | 0.19 | 36.1 | 20.7 | 3.3 | 0 | na | na | na | na | na | na |
| *D. recens* | 32 | 21.9 | 68.8 | **90.6** | **9.4** | 30 | 22.37 | 1.15 | 0.20 | 33.9 | 20.3 | 3.6 | 0 | na | na | na | na | na | na |
| *D. transversa* | 40 | 70.0 | 30.0 | **100.0** | **0.0** | 40 | 21.62 | 0.29 | 0.05 | 194.4 | 140.8 | 22.3 | 37 | 24.63 | 0.93 | 0.15 | 77.1 | 70.4 | 11.1 |
