## Supplementary file 5_Period_AvLatitude_Corr for "Fruit flies of the subgenera *Drosophila*, *Siphlodora* and *Sophophora* use different mechanisms to adapt their daily activity to long photoperiods"

**Supplementary file 5** Correlations between mean free-running period length in DD, mean latitudinal distribution and original latitude of the strain in different sample groups.

| Sample (Size) | Variable 1 | Variable 2 | **Spearman's correlation ρ** | **Spearman's correlation p-value** |
| --- | --- | --- | --- | --- |
| selected species strains (52) | Period τ in DD | Mean latitudinal distr. (3 % trimmed) | **-0.137** | **0.373** |
| selected species strains (52) | Period τ in DD | Original latitude of tested strain | **0.037** | **0.793** |
| *Sophophora* (11) | Period τ in DD | Mean latitudinal distr. (3 % trimmed) | **0.182** | **0.593** |
| *Sophophora* (11) | Period τ in DD | Original latitude of tested strain | **0.527** | **0.096** |
| *Siphlodora* (16) | Period τ in DD | Mean latitudinal distr. (3 % trimmed) | **-0.341** | **0.196** |
| *Siphlodora* (16) | Period τ in DD | Original latitude of tested strain | **-0.029** | **0.914** |
| *Drosophila* (16) | Period τ in DD | Mean latitudinal distr. (3 % trimmed) | **-0.365** | **0.165** |
| *Drosophila* (16) | Period τ in DD | Original latitude of tested strain | **-0.157** | **0.560** |
