## Supplementary file 6_Strain_Information for "Fruit flies of the subgenera *Drosophila*, *Siphlodora* and *Sophophora* use different mechanisms to adapt their daily activity to long photoperiods"

### Fly strains, rearing information, and experimental specifications

**Supplementary file 6A** Fruit fly species and strains used in this study. Biogeographic origins of speciation are derived from Russo et al. (2013) and Izumitani et al. (2016). Current distributions were taken from TaxoDros: AF, Africa; PA, Palearctic (Eurasia); NT, Neotropic (South America); NA, Nearctic (North America); SA, Saharo-Arabian; OR Oriental (India, Southeast Asia); SJ, Sino-Japanese; APC, Australasian and Pacific; HW, Hawaiian; CMP, Cosmopolitan (worldwide). Question marks (?) indicate biogeographic origins that remain under debate. NDSSC, National *Drosophila* Species Stock Center at Cornell University. Strains labeled “collected” were newly caught in the wild for this study.

| **Fruit fly species (strain)** | **Genus (subgenus)** | **Group (subgroup)** | **Biogeographic origin (speciation)** | **Current distribution** | **source of strain** | **collection site (if known)** | **geographic coordinates (decimal)** | **collection year (if known)** |
| --- | --- | --- | --- | --- | --- | --- | --- | --- |
| *Scaptodrosophila latifasciaeformis* | *Scaptodrosophila* |  | PA | CMP (Tropics) | NDSSC | Barro Colorado Island, Canal Zone, Panama | ≈ 9.1580556,  -79.84808333 | 1955 |
| *S. lebanonensis* | *Scaptodrosophila* |  | PA | PA, SA | NDSSC | Beirut, Lebanon | ≈ 33.8597222, 35.57027778 | 1948 |
| *S. rufifrons* | *Scaptodrosophila* |  | PA | PA | NDSSC | Sochi, Krasnodar Krai, Russia | 43.583333,  39.71666667 |  |
| *Hirtodrosophila duncani* | *Collesia*‑*H. duncani* clade | *duncani* | NA? | NA | Kyorin-Fly | Florida, USA | ≈ 30.43, -83.92 | 2000 |
| *Drosophila paulistorum* | *Drosophila* (*Sophophora*) | *willistoni* (*willistoni*) | NT | NT | NDSSC | Amazonas, Brazil | ≈ -5, -63 |  |
| *D. nebulosa* | *Drosophila* (*Sophophora*) | *willistoni* (*bocainensis*) | NT | NT | NDSSC | Tecualilla, Sinaloa, Mexico | 22.7675,  -105.6705556 | 2004 |
| *D. prosaltans* | *Drosophila* (*Sophophora*) | *saltans* (*saltan*s) | NT | NT | NDSSC | Eldorado, São Paulo, Brazil | ≈ -24.5166667,  -48.25 |  |
| *D. sturtevanti* | *Drosophila* (*Sophophora*) | *saltans* (*sturtevanti*) | NT | NT | NDSSC | Costa Rica | ≈ 10, -84 | 2014 |
| *D. melanogaster* | *Drosophila* (*Sophophora*) | *melanogaster* (*melanogaster*) | AF | CMP | Menegazzi et al., 2017 | Tanzania | ≈ -3.1833333, 37.23333333 | 2014 |
| *D. kikkawai* | *Drosophila* (*Sophophora*) | *melanogaster* (*montium*) | PA, OR | CMP (Tropic) | NDSSC | Taipei, Taiwan | 25.0175, 121.5908333 | 2009 |
| *D. ananassae* | *Drosophila* (*Sophophora*) | *melanogaster* (*ananassae*) | PA, APC | CMP (Tropic) | NDSSC | Mbengwi, Cameroon | ≈ 6.0077778, 10.02472222 | 2004 |
| *D. obscura* | *Drosophila* (*Sophophora*) | *obscura* (*obscura*) | PA | PA | Ehime-Fly | Oulu, Finland | ≈ 65.0166667, 25.46666667 | 2012 |
| *D. bifasciata* | *Drosophila* (*Sophophora*) | *obscura* (*obscura*) | PA | PA | collected (Peter Deppisch) | Oellingen, Bayern, Germany | 49.5788889, 10.06166667 | 2017 |
| *D. subobscura* | *Drosophila* (*Sophophora*) | *obscura* (*obscura*) | PA | CMP (Temperate) | collected (Pamela Menegazzi) | Gerbrunn, Bayern, Germany | 49.7783333, 9.983611111 | 2020 |
| *D. subobscura* | *Drosophila* (*Sophophora*) | *obscura* (*obscura*) | PA | CMP (Temperate) | NDSSC | Heidelberg, BW, Germany | 49.3833333, 8.716666667 | 2005 |
| *D. azteca* | *Drosophila* (*Sophophora*) | *obscura* (*affinis*) | NA | NA | NDSSC | San Pablo Etla, Oaxaca, Mexico | ≈ 17.1666667,  -96.71666667 | 2003 |
| *D. busckii* | *Drosophila* (*Dorsilopha*) | *busckii* | PA | CMP | collected (Peter Deppisch) | Oellingen, Bayern, Germany | 49.5788889, 10.06166667 | 2017 |
| *Scaptomyza pallida* (2017) | *Scaptomyza* (*Parascaptomyza*) |  | CMP, HW | CMP | collected (Peter Deppisch) | Oellingen, Bayern, Germany | 49.5788889, 10.06166667 | 2017 |
| *S. pallida* (2020) | *Scaptomyza* (*Parascaptomyza*) |  | CMP, HW | CMP | collected (Peter Deppisch) | Oellingen, Bayern, Germany | 49.5788889, 10.06166667 | 2020 |
| *Drosophila robusta* | *Drosophila* (*Siphlodora*) | *robusta* (*robusta*) | PA | NA | NDSSC | Iowa River, Iowa, USA | ≈ 42,  -91.96666667 | 2003 |
| *D. tsigana* | *Drosophila* (*Siphlodora*) | *melanica* | PA | PA | Ehime-Fly | Hokkaidō, Japan | ≈ 43.4333333, 142.7166667 | 2004 |
| *D. melanica* | *Drosophila* (*Siphlodora*) | *melanica* | NA | NA | NDSSC | Austin, Texas, USA | ≈ 30.3833333, -97.76666667 |  |
| *D. gaucha* | *Drosophila* (*Siphlodora*) | *mesophragmatica* | NT | NT | NDSSC | Buenos Aires, Argentina | ≈ -36.0847222, -60.08555556 | 2006 |
| *D. hydei* | *Drosophila* (*Siphlodora*) | *repleta* (*hydei*) | NT | CMP | Beauchamp et al., 2018 | Nkweseko,  Machame, Tanzania | ≈ -3.1833333, 37.23333333 | 2014 |
| *D. repleta* (K) | *Drosophila* (*Siphlodora*) | *repleta* (*repleta*) | NT, CMP | CMP | collected (P. Deppisch) | Oellingen, Bayern, Germany | 49.5788889, 10.06166667 | 2017 |
| *D. repleta* (S) | *Drosophila* (*Siphlodora*) | *repleta* (*repleta*) | NT, CMP | CMP | collected (Peter Deppisch) | Sonderhofen, Bayern, Germany | 49.5936111, 10.00194444 | 2017 |
| *D. mercatorum* | *Drosophila* (*Siphlodora*) | *repleta* (*mercatorum*) | NT, CMP | CMP | Beauchamp et al., 2018 | Nkweseko,  Machame, Tanzania | ≈ -3.1833333, 37.23333333 | 2014 |
| *D. kanekoi* | *Drosophila* (*Siphlodora*) | *virilis* | PA | PA (SJ) | Kyorin-Fly | Rikubetsu, Hokkaido, Japan | ≈ 43.5188889, 143.675 | 2017 |
| *D. ezoana* | *Drosophila* (*Siphlodora*)*)* | *virilis* | PA | PA | Hannele Kauranen,  Lilli Bähr | Oulanka, Finland | ≈ 66.4653333, 29.03677778 | 2017 |
| *D. littoralis* | *Drosophila* (*Siphlodora*) | *virilis* | PA | PA | Pekka Lankinen | Kilpisjärvi,  Finland | ≈ 69.0442222, 20.80344444 | 1970 |
| *D. virilis* | *Drosophila* (*Siphlodora*) | *virilis* | PA | CMP | NDSSC | Gikongoro, Rwanda | ≈ -2.06, 29.96 | 2009 |
| *D. virilis* | *Drosophila* (*Siphlodora*) | *virilis* | PA | CMP | Ehime-Fly | Yamagata, Japan | ≈ 38, 140 | 2006 |
| *D. lummei* | *Drosophila* (*Siphlodora*) | *virilis* | PA | PA | Anneli Hoikkala | Kuusamo, Oulanka, Finland | ≈ 65.95, 29.16666667 | 2013 |
| *D. novamexicana* | *Drosophila* (*Siphlodora*) | *virilis* | NA | NA | NDSSC | San Antonio, New Mexico, USA | ≈ 33.9166667,  -106.8666667 | 1947 |
| *D. americana texana* | *Drosophila* (*Siphlodora*) | *virilis* | NA | NA | NDSSC | Brewer Basin, Tennessee, USA | ≈ 36.4666667,  -89.315067 | 2005 |
| *D. borealis* (eastern) | *Drosophila* (*Siphlodora*) | *virilis* | NA | NA | NDSSC | Poygan Marsh, Wisconsin, USA | ≈ 44.1, -88.9 | 2005 |
| *D. lacicola* | *Drosophila* (*Siphlodora*) | *virilis* | NA | NA | NDSSC | Beaver Creek, Manitoba, Canada | ≈ 51.3833333,  -96.93 | 1949 |
| *Zaprionus indianus* | *Zaprionus*-clade (*Zaprionus*) |  | CMP, AF | CMP | NDSSC | Alexandria, Egypt | ≈ 31.2166667, 29.96666667 |  |
| *Z. camerounensis* | *Zaprionus*-clade (*Zaprionus*) | *vittiger* subcomplex | AF | AF | NDSSC | East Usambara Mountains, Amani, Tanzania | ≈ -5.0833333, 38.66666667 |  |
| *Drosophila repletoides* | *Zaprionus*-clade | *tumiditarsus* | PA | PA (SJ) | Kyorin-Fly | Tokyo, Japan | ≈ 35.7713889, 139.1002778 | 1965 |
| *D. immigrans* | *Drosophila* (*Drosophila*) | *immigrans* (*immigrans*) | OR | CMP | collected (Peter Deppisch) | Oellingen, Bayern, Germany | 49.5788889, 10.06166667 | 2017 |
| *D. hypocausta* | *Drosophila* (*Drosophila*) | *immigrans* (*hypocausta*) | OR, APC | OR, APC | Kyorin-Fly | Guam, USA | ≈ 13.4666667, 144.7666667 | 2012 |
| *D. albomicans* | *Drosophila* (*Drosophila*) | *immigrans* (*nasuta*) | PA, OR | PA, SJ, OR, APC | Kyorin-Fly | Iriomote, Japan | ≈ 24.3333333, 123.8166667 | 2017 |
| *D. sulfurigaster blilimbata* | *Drosophila* (*Drosophila*) | *immigrans* (*nasuta*) | OR, APC | OR, APC, HW | Kyorin-Fly | Guam, USA | ≈ 13.4666667, 144.7666667 | 2012 |
| *D. tripunctata* | *Drosophila* (*Drosophila*) | *tripunctata* (*tripunctata* IV) | NA | NA, NT | Clare Scott-Chialvo | Athens, Georgia,  USA  (strain IsoF 1) | ≈ 33.9,  -83.38333333 | 2007 |
| *D. polymorpha* | *Drosophila* (*Drosophila*) | *cardini* (*cardini*) | NT | NT | NDSSC | Morro da Lagoa, Florianópolis, SC, Brazil | ≈ -27.6,  -48.48333333 | 2012 |
| *D. cardini* | *Drosophila* (*Drosophila*) | *cardini* (*cardini*) | NT | NT, HW, (NA) | Clare Scott-Chialvo | Florida,  USA  (strain F55) | ≈ 28, -82 | 2004 |
| *D. funebris* (Alaska) | *Drosophila* (*Drosophila*) | *funebris* (*funebris*) | PA (NA) | CMP | NDSSC | Big Lake, Alaska, USA | ≈ 61.5166667,  -149.9 | 1957 |
| *D. funebris* (Bavaria Q) | *Drosophila* (*Drosophila*) | *funebris* (*funebris*) | PA (NA) | CMP | collected (Peter Deppisch) | Sonderhofen, Bayern, Germany | 49.5936111, 10.00194444 | 2017 |
| *D. funebris* (Bavaria V) | *Drosophila* (*Drosophila*) | *funebris* (*funebris*) | PA (NA) | CMP | collected (Peter Deppisch) | Sonderhofen, Bayern, Germany | 49.5936111, 10.00194444 | 2017 |
| *D. funebris* (Mexico) | *Drosophila* (*Drosophila*) | *funebris* (*funebris*) | PA (NA) | CMP | NDSSC | Mexico City, Mexico | ≈ 19.5833333,  -99.11666667 |  |
| *D. funebris* (Lebanon) | *Drosophila* (*Drosophila*) | *funebris* (*funebris*) | PA (NA) | CMP | NDSSC | Ain Anoub, Lebanon | ≈ 33.7808333, 35.54638889 |  |
| *D. funebris* (Hokkaido) | *Drosophila* (*Drosophila*) | *funebris* (*funebris*) | PA (NA) | CMP | Kyorin-Fly | Hokkaidō, Japan | ≈ 43.4333333, 142.7166667 | 1987 |
| *D. multispina* | *Drosophila* (*Drosophila*) | *funebris* | PA | PA, SJ | Kyorin-Fly | Hokkaidō, Japan | ≈ 43.4333333, 142.7166667 | 2007 |
| *D. macrospina* (Arizona) | *Drosophila* (*Drosophila*) | *funebris* (*macrospina*) | NA | NA | NDSSC | Sonoita creek, Patagonia, Arizona, USA | ≈ 31.4833333,  -110.85 | 1958 |
| *D. macrospina* (South Carolina) | *Drosophila* (*Drosophila*) | *funebris* (*macrospina*) | NA | NA | NDSSC | Cross Anchor, South Carolina, USA | ≈ 34.65,  -81.86666667 |  |
| *D. putrida* | *Drosophila* (*Drosophila*) | *testacea* | NA | NA | Kyorin-Fly | Nebraska, USA | ≈ 41, -99 | 1950 |
| *D. testacea* | *Drosophila* (*Drosophila*) | *testacea* | PA | CMP (PA + NA) | Steve Perlman | St. Sulpice, Vaud, Switzerland | ≈ 46.5088889, 6.560277778 | 2012 |
| *D. brachynephros* | *Drosophila* (*Drosophila*) | *quinaria* | PA | PA, SJ | Kyorin-Fly | Ehime, Japan | ≈ 33, 133 | 2004 |
| *D. deflecta* | *Drosophila* (*Drosophila*) | *quinaria* | NA | NA | Kyorin-Fly | New Jersey, USA | ≈ 40.36, -74.67 | 2000 |
| *D. recens* | *Drosophila* (*Drosophila*) | *quinaria* | NA | NA | Clare Scott-Chialvo | New York,  USA  (mixed strain) | ≈ 43, -74 | 2009 |
| *D. transversa* | *Drosophila* (*Drosophila*) | *quinaria* | NA | NA, PA | Clare Scott-Chialvo | Uppsala,  Schweden  (strain isoF 5) | ≈ 59.8, 17.7 | 2005 |

**Supplementary file 6B** Data for maintenance and rearing of the different fruit fly species. Flies were kept in plastic vials (45 mm in diameter, 104 mm in height) filled to a depth of approximately 2 cm with either standard or banana food. Mushroom-feeding species were additionally supplied with ≈ 4 to 5 g of defrosted, sliced button mushrooms. Additionally, in order to facilitate pupation and to control food consistency and moisture, most food vials were also equipped with cotton paper strips (40 × 30 × 1 mm) and/or round coffee filters (Ø 45 mm).

| **Fruit fly species** | **Food** | **Food supply (cotton paper (C), filter (F))** | **Rearing temperature** | **Rearing light regime** |
| --- | --- | --- | --- | --- |
| *S. latifasciaeformis* | Banana | 1 C + 1 F | 25 °C | LD 12:12 |
| *S. lebanonensis* | Banana | 1 C + 1 F | 18 °C | LD 12:12 |
| *S. rufifrons* | Banana | 1 C + 2 F | 25 °C | LD 12:12 |
| *H. duncani* | Banana | 3 F | 25 °C | LD 12:12 |
| *D. paulistorum* | Standard | 1C + 1 F | 18 °C | LD 12:12 |
| *D. nebulosa* | Standard | 2 C | 18 °C | LD 12:12 |
| *D. prosaltans* | Standard | 1 C + 1 F | 18 °C | LD 12:12 |
| *D. sturtevanti* | Standard | 2 C | 25 °C | LD 12:12 |
| *D. melanogaster* | Standard | - | 18 °C | LD 12:12 |
| *D. kikkawai* | Banana | 1 C + 1 F | 18 °C | LD 12:12 |
| *D. ananassae* | Standard | 1 C + 1 F | 25 °C | LD 12:12 |
| *D. obscura* | Standard | 3 F | 21 °C | LD 20:4 |
| *D. bifasciata* | Standard | 3 F | 18 °C | LD 12:12 |
| *D. subobscura* | Standard | 1 C + 2 F | 18 °C | LD 12:12 |
| *D. azteca* | Mushroom | 2 F | 21 °C | LD 20:4 |
| *D. busckii* | Standard | 2 F | 18 °C | LD 12:12 |
| *Scaptomyza pallida* | Standard | 1 C + 1 F | 18 °C | LD 12:12 |
| *D. robusta* | Banana | 1 C + 1 F | 25 °C | LD 12:12 |
| *D. tsigana* | Banana | 3 F | 21 °C | LD 20:4 |
| *D. melanica* | Standard | 2 C | 18 °C | LD 12:12 |
| *D. gaucha* | Banana | 1 C + 1 F | 21 °C | LD 20:4 |
| *D. hydei* | Standard | 2 C | 18 °C | LD 12:12 |
| *D. repleta* | Standard | 1 C + 1 F | 25 °C | LD 12:12 |
| *D. mercatorum* | Standard | 2 C | 18 °C | LD 12:12 |
| *D. kanekoi* | Standard | 1 C + 1 F | 25 °C | LD 12:12 |
| *D. ezoana* | Standard | 2 C | 21 °C | LD 20:4 |
| *D. littoralis* | Standard | 2 C | 21 °C | LD 20:4 |
| *D. virilis* | Standard | 2 C | 18 °C | LD 12:12 |
| *D. lummei* | Standard | 2 C | 21 °C | LD 20:4 |
| *D. novamexicana* | Standard | 2 C | 18 °C | LD 12:12 |
| *D. americana texana* | Standard | 2 C | 18 °C | LD 12:12 |
| *D. borealis* (eastern) | Standard | 2 C | 21 °C | LD 20:4 |
| *D. lacicola* | Standard | 2 C | 21 °C | LD 20:4 |
| *Z. indianus* | Banana | 2 C | 18 °C | LD 12:12 |
| *Z. camerounensis* | Banana | 2 C | 18 °C | LD 12:12 |
| *D. repletoides* | Standard | 2 C | 25 °C | LD 12:12 |
| *D. immigrans* | Standard | 1 C + 1 F | 18 °C | LD 12:12 |
| *D. hypocausta* | Banana | 1 C + 1 F | 25 °C | LD 12:12 |
| *D. albomicans* | Standard | 1C + 1 F | 25 °C | LD 12:12 |
| *D. sulfurigaster blilimbata* | Standard | 1 C + 1 F | 25 °C | LD 12:12 |
| *D. tripunctata* | Mushroom | 1 C + 1 F | 21 °C | LD 20:4 |
| *D. polymorpha* | Standard | 1 C + 2 F | 18 °C | LD 12:12 |
| *D. cardini* | Mushroom | 1 C + 1 F | 21 °C | LD 20:4 |
| *D. funebris* | Standard | 2 C | 18 °C | LD 12:12 |
| *D. multispina* | Mushroom | 1 C + 1 F | 21 °C | LD 20:4 |
| *D. macrospina* | Standard | 1 C + 1 F | 18 °C | LD 12:12 |
| *D. putrida* | Banana | 3 F | 25 °C | LD 12:12 |
| *D. testacea* | Mushroom | 1 C + 1 F | 21 °C | LD 20:4 |
| *D. brachynephros* | Mushroom | 1 C + 1 F | 18 °C | LD 12:12 |
| *D. deflecta* | Mushroom | 1 C + 1 F | 18 °C | LD 12:12 |
| *D. recens* | Mushroom | 1 C + 1 F | 18 °C | LD 12:12 |
| *D. transversa* | Mushroom | 1 C + 1 F | 18 °C | LD 12:12 |

**Supplementary file 6C** List of locomotor activity tube sizes, fixation times, and antibodies used for immunohistochemical stainings in the dfferent fly species. Some species were recorded in both 5 mm and 7 mm tubes for practical reasons, since two different species were sometimes housed in a single monitor.The standard antibody used for PDF staining was a monoclonal mouse anti-PDF (C7 line; Developmental Studies Hybridoma Bank; 1:2000). Where this antibody did not stain PDF sufficiently, polyclonal rabbit anti-β-PDH (1:400; Dircksen et al., 1987) was used instead.

| **Fruit fly species** | **DAM (TriKinetics) tube diameter** | **Fixation time for IHC** | **Antibody against PDF** |
| --- | --- | --- | --- |
| *S. latifasciaeformis* | 5 mm | 2.5 h | PDF c7 |
| *S. lebanonensis* | 7 mm | 3.0 h | PDF c7 |
| *S. rufifrons* | 5 mm, 7 mm | 3.0 h | PDF c7 |
| *H. duncani* | 7 mm | 3.0 h | PDF c7 |
| *D. paulistorum* | 5 mm | 2.5 h | PDF c7 |
| *D. nebulosa* | 7 mm | 3.0 h | PDF c7 |
| *D. prosaltans* | 5 mm, 7 mm | 2.5 h | PDF c7 |
| *D. sturtevanti* | 5 mm, 7 mm | 3.0 h | PDF c7 |
| *D. melanogaster* | 5 mm | 2.5 h | PDF c7 |
| *D. kikkawai* | 5 mm | 3.0 h | PDF c7 |
| *D. ananassae* | 5 mm | 2.5 h | PDF c7 |
| *D. obscura* | 5 mm, 7 mm | 3.0 h | β-PDH |
| *D. bifasciata* | 5 mm, 7 mm | 3.0 h | β-PDH |
| *D. subobscura* | 7 mm | 3.0 h | β-PDH |
| *D. azteca* | 7 mm | 2.75 h | β-PDH |
| *D. busckii* | 5 mm | 2.5 h | β-PDH |
| *Scaptomyza pallida* | 7 mm | 2.75 h | β-PDH |
| *D. robusta* | 7 mm | 3.25 h | PDF c7 |
| *D. tsigana* | 7 mm | 3.25 h | PDF c7 |
| *D. melanica* | 7 mm | 3.25 h | PDF c7 |
| *D. gaucha* | 7 mm | 3.25 h | β-PDH/PDF c7 |
| *D. hydei* | 7 mm | 3.25 h | β-PDH |
| *D. repleta* | 7 mm | 3.25 h | β-PDH |
| *D. mercatorum* | 7 mm | 3.25 h | β-PDH |
| *D. kanekoi* | 7 mm | 3.25 h | PDF c7 |
| *D. ezoana* | 7 mm | 3.25 h | PDF c7 |
| *D. littoralis* | 7 mm | 3.25 h | PDF c7 |
| *D. virilis* | 7 mm | 3.25 h | PDF c7 |
| *D. lummei* | 7 mm | 3.25 h | PDF c7 |
| *D. novamexicana* | 7 mm | 3.25 h | PDF c7 |
| *D. americana texana* | 7 mm | 3.25 h | PDF c7 |
| *D. borealis* (eastern) | 7 mm | 3.25 h | PDF c7 |
| *D. lacicola* | 7 mm | 3.25 h | PDF c7 |
| *Z. indianus* | 7 mm | 3.25 h | PDF c7 |
| *Z. camerounensis* | 7 mm | 3.25 h | PDF c7 |
| *D. repletoides* | 7 mm | 3.0 h | β-PDH/PDF c7 |
| *D. immigrans* | 7 mm | 3.25 h | PDF c7 |
| *D. hypocausta* | 7 mm | 3.25 h | PDF c7 |
| *D. albomicans* | 7 mm | 3.25 h | PDF c7 |
| *D. sulfurigaster blilimbata* | 7 mm | 3.25 h | PDF c7 |
| *D. tripunctata* | 7 mm | 3.0 h | PDF c7 |
| *D. polymorpha* | 5 mm, 7 mm | 3.0 h | β-PDH |
| *D. cardini* | 7 mm | 3.0 h | β-PDH |
| *D. funebris* | 7 mm | 3.25 h | PDF c7 |
| *D. multispina* | 7 mm | 3.25 h | PDF c7 |
| *D. macrospina* | 7 mm | 3.25 h | PDF c7 |
| *D. putrida* | 7 mm | 3.0 h | PDF c7 |
| *D. testacea* | 7 mm | 2.5 h | PDF c7 |
| *D. brachynephros* | 7 mm | 3.25 h | PDF c7 |
| *D. deflecta* | 7 mm | 3.25 h | PDF c7 |
| *D. recens* | 7 mm | 3.0 h | PDF c7 |
| *D. transversa* | 7 mm | 3.25 h | PDF c7 |
