## Supplementary material for "Fruit flies of the subgenera *Drosophila*, *Siphlodora* and *Sophophora* use different mechanisms to adapt their daily activity to long photoperiods": Main Figures in High Resolution

bioRxiv, 2026

### Figures from the main article in high resolution

**Figure 1: Simplified clock network of the brain (PDF and CRY expression scheme)**

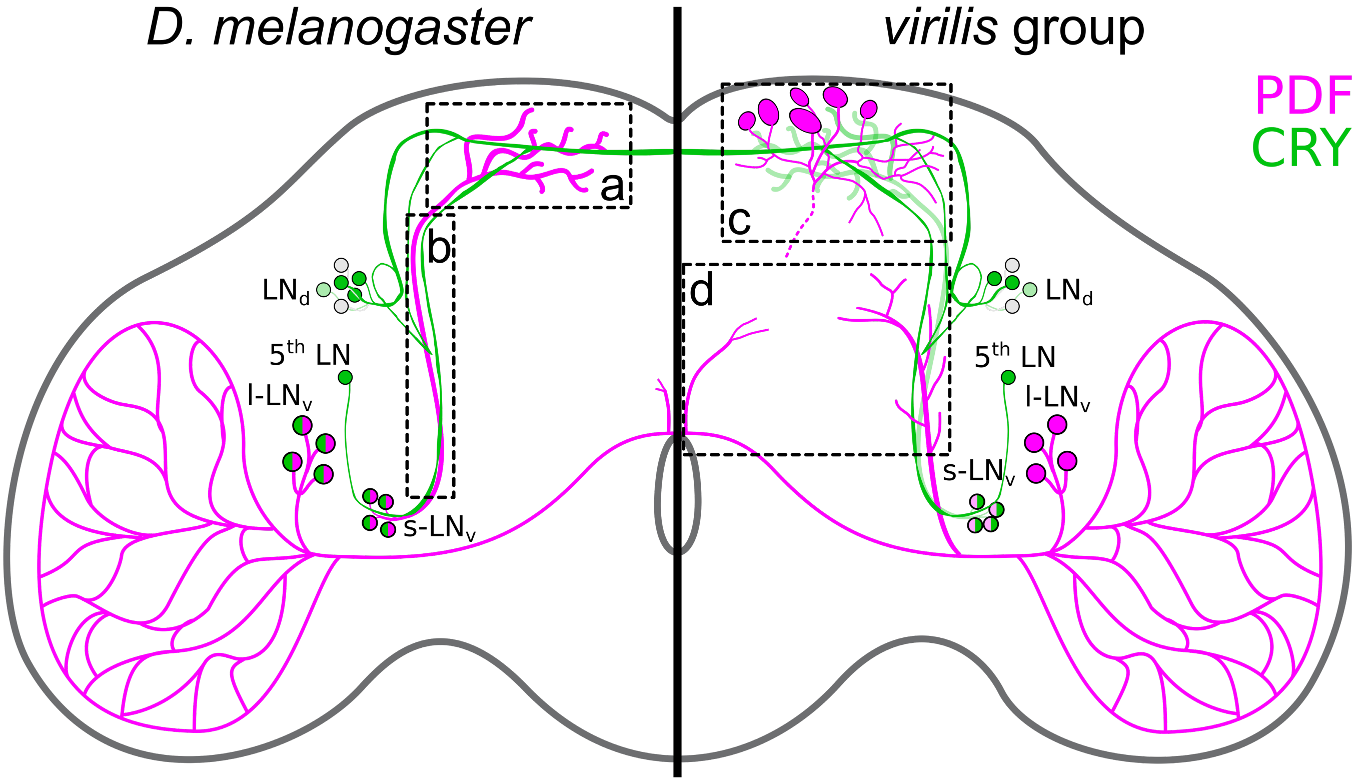

**Figure 1.** Simplified clock network in the brain of *Drosophila melanogaster* (left) and *Drosophila* species from the *virilis* group (right). Only a subset of clock neurons is shown: neurons associated with morning activity (small ventrolateral neurons, s-$LN_v$) and evening activity (the fifth lateral neuron, $5^{th}$ LN, and dorsolateral neurons, $LN_d$). PDF and CRY expression are shown in magenta and green, respectively. The *virilis* group species display markedly weaker PDF expression in the s-$LN_v$, contain extra PDF-positive cells in the dorsolateral protocerebrum, extra PDF fibers in the central brain, and lack CRY expression in the large ventrolateral neurons (l-$LN_v$). Stippled rectangles indicate the brain areas used to assess PDF staining across the different *Drosophila* species in this study: a, PDF staining intensity in the s-$LN_v$ terminals; b, presence of PDF-positive varicosities along the s-$LN_v$ axons; c, PDF-fiber density of non-clock neurons in the vicinity of the s-$LN_v$ terminals; d, PDF-fiber density in the centrolateral brain.

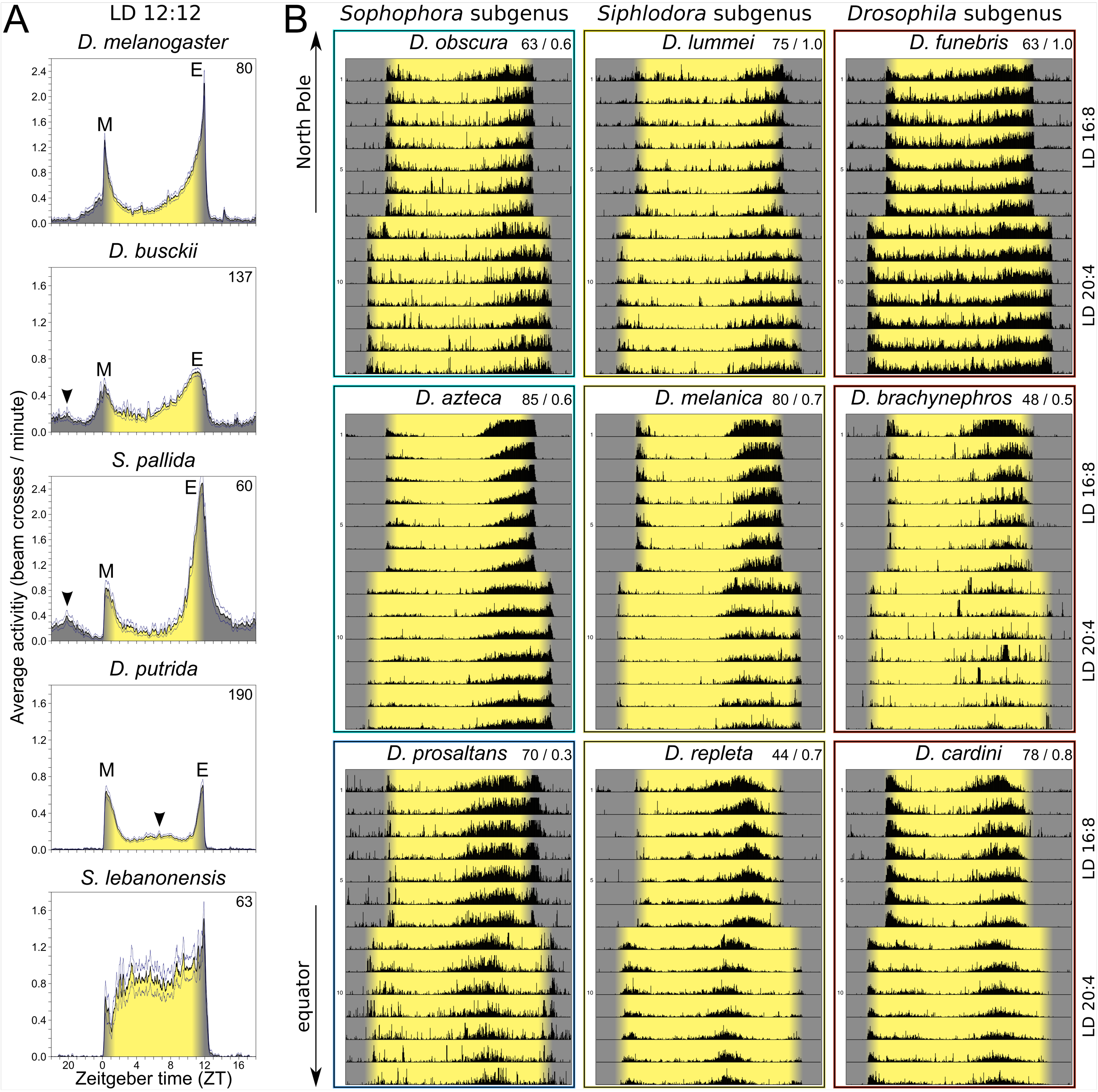

**Figure 2.** Phylogenetic relationship of the 52 fruit fly species used in this study. The divergence of the species is displayed as a chronogram with a timescale in million years ago (MYA). Relationships and divergence events were inferred from the studies of Morales-Hojas et al. (2011), Oliveira et al. (2012), Russo et al. (2013), Izumitani et al. (2016), Finet et al. (2021), Suvorov et al. (2022), and Tanaka et al. (2022). Note that the nodes are estimates and become less certain the further back in time one goes. The symbols * and # indicate divergence times that are controversial between different studies.

**Figure 3: Latitudinal distribution pattern of the 52 tested fruit fly species**

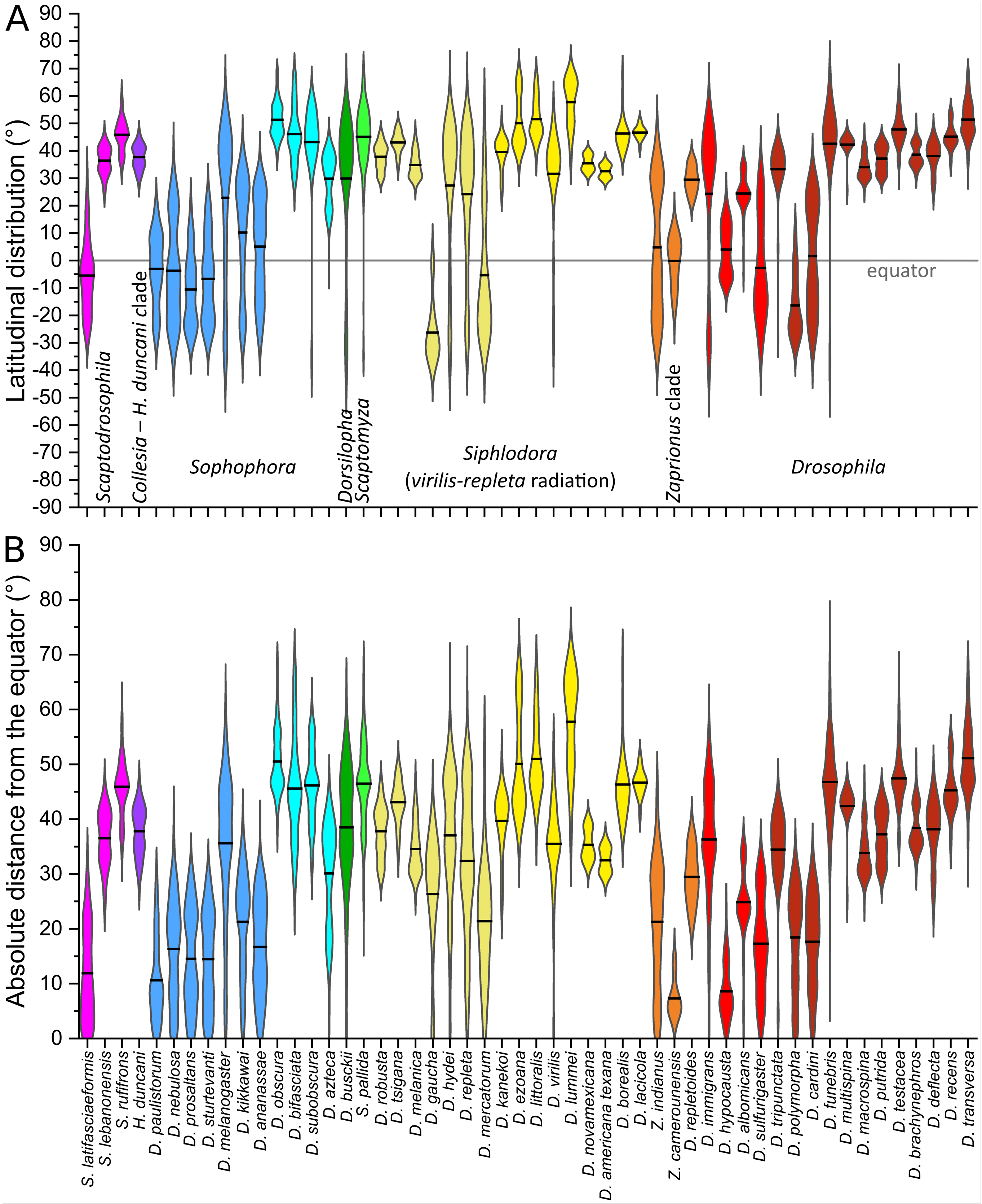

**Figure 3.** Latitudinal distribution pattern of the 52 *Drosophila* species investigated in this study. Reported collection sites were extracted from Gerhard Bächli's Taxodros website. Species are arranged by phylogenetically related group, using the same color code as in Figure 2). **(A)** Violin plots of the global latitudinal distribution of all species with arithmetic means (thick lines). **(B)** Violin plots of the absolute distance of each species' latitudinal coordinates from the equator.

**Figure 4: LD 12:12 daily activity profiles and average actograms (LD 16:8, LD 20:4)**

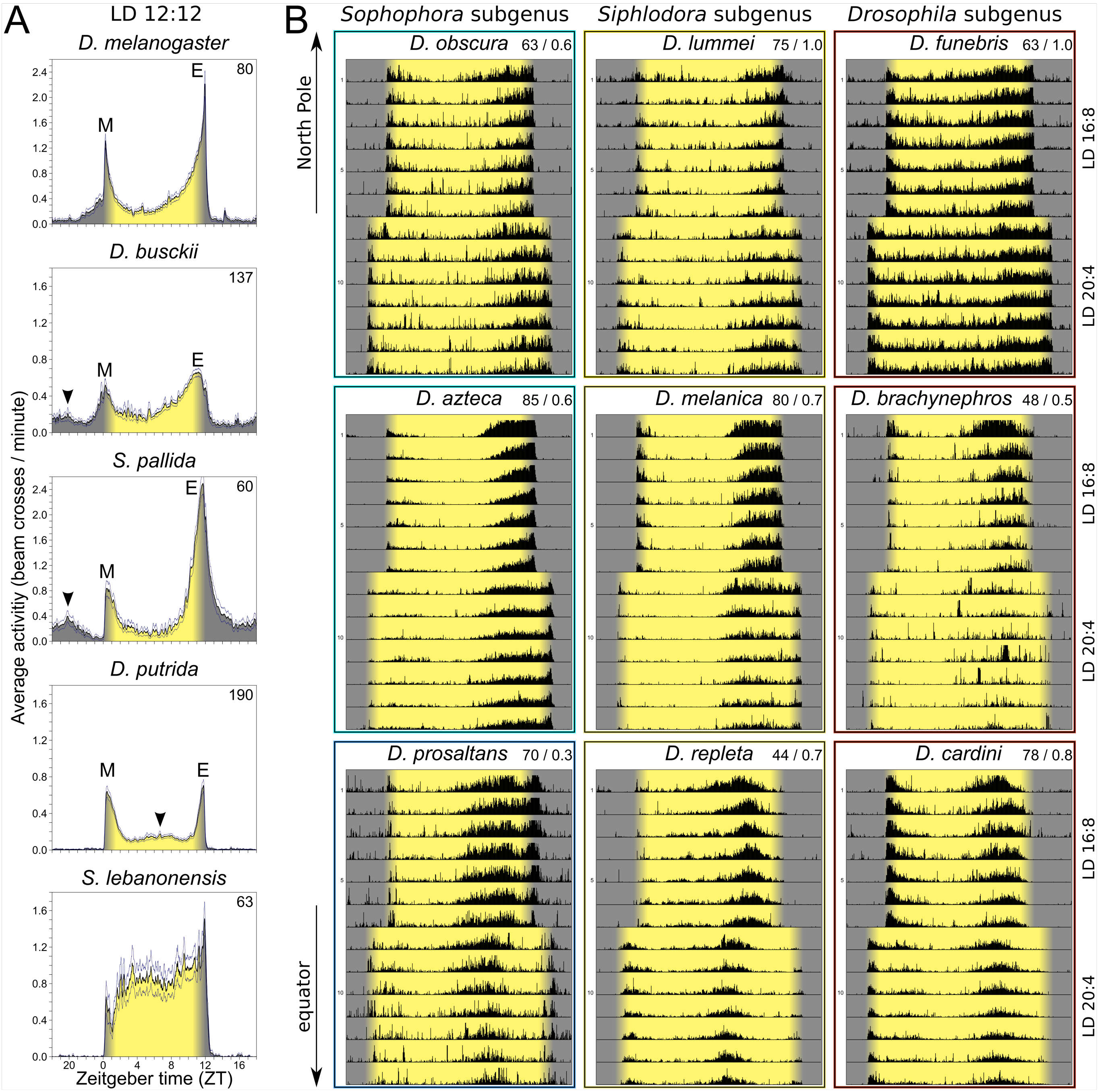

**Figure 4.** Average activity profiles under LD 12:12, and average actograms under LD 16:8 and 20:4. **(A)** Average activity profile of *D. melanogaster* and four species with slightly different activity patterns. Besides the typical morning (M) and evening (E) activity bouts, *D. busckii* and *S. pallida* showed nocturnal activity bouts, and *D. putrida* displayed midday activity (arrowheads). *S. lebanonensis* exhibited high activity throughout the day, lacking a siesta. The number of flies averaged for each profile is indicated in the top right corner. Thin lines above and below the mean indicate the standard error of the mean. **(B)** Average actograms of representative species from the three main *Drosophila* subgenera. The first row shows high latitude species, the second row species from temperate climate zones, and the bottom row species from low latitudes; only *D. repleta* is cosmopolitan and also occuring in temperate regions. The first seven days show recordings under LD 16:8, and the last seven days recordings under LD 20:4. The light phase is highlighted with a yellow background, the light-ramping (dawn/dusk simulation) is highlighted with a beige background, and the dark phase with a grey background. The frame colors refer to the phylogenetic color-coding from Figure 2. The number of flies and the upper activity threshold (lower for less active species and higher for highly active species) are given in the top right corner of each actogram. Each black line in the actograms represents an averaged 3-minute bin.

#### Figure 5: Average daily activity profiles under LD 20:4

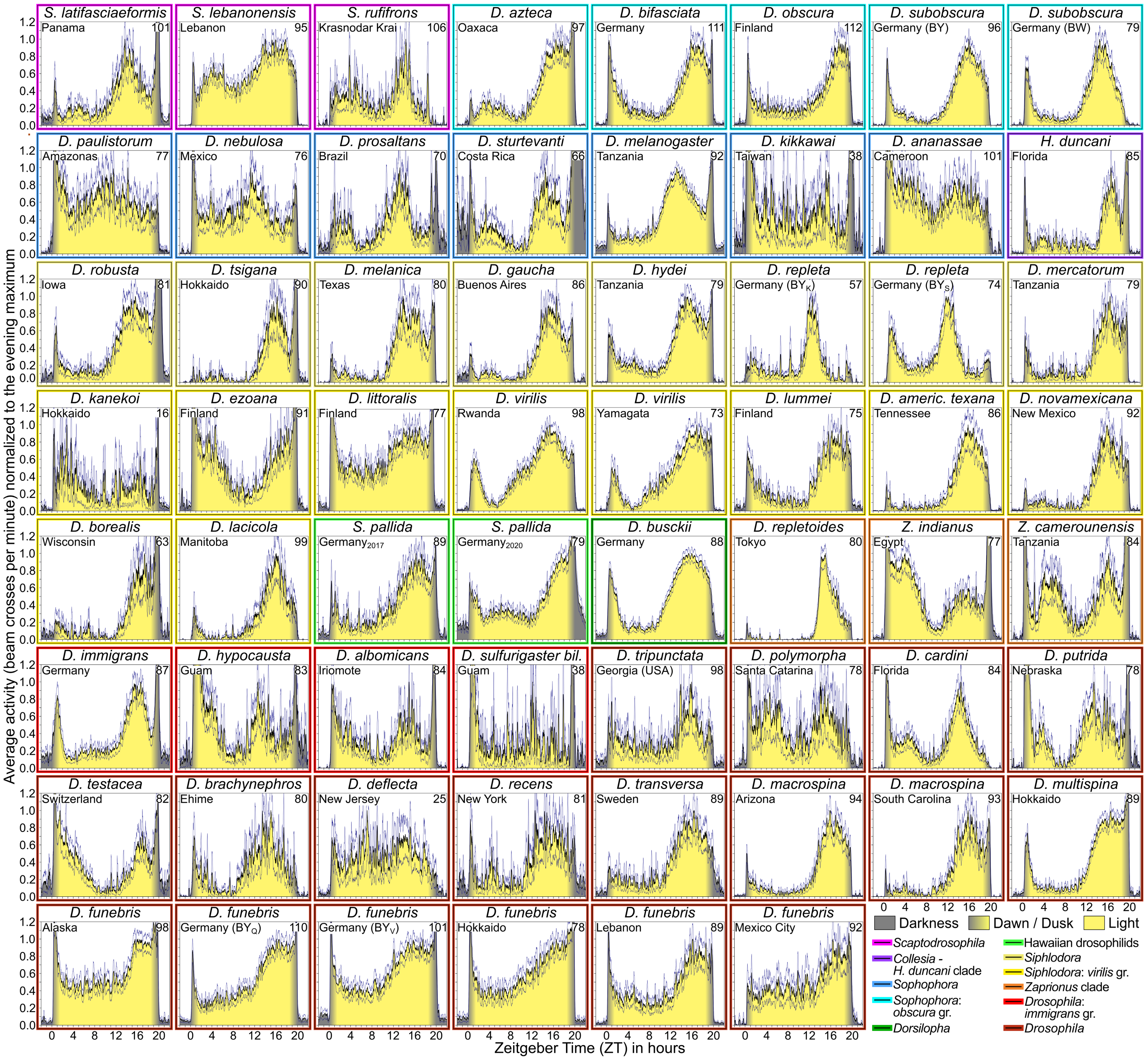

**Figure 5**. Average daily activity profiles of all tested species and strains in LD 20:4. Curves are normalized to the maximum amplitude of the circadian-driven morning or evening peaks. The origin of the species (or strain) is indicated in the upper left corner of each diagram, and the number of flies included is indicated in the upper right corner. Individuals that free-ran under LD 20:4 were omitted from the calculation of the daily activity profile (concerning the species *D. kikkawai*, *D. kanekoi*, *D. sulfurigaster bilimbata*, and *D. deflecta*). Frame colors refer to the phylogenic color-coding from Figure 2 (also shown in the lower right corner). During normalization, maximal activity during the evening peak was set to 1. This was done to improve visualization, since some species were very inactive. The magnitude of startle responses to dawn and dusk events was disregarded (e.g., the lights-off startle response was unusually high and long-lasting in *D. sturtevanti*, second row).

#### Figure 6: Correlation between $\Delta\Psi_{M\text{-}E}$ in LD 20:4 and average latitude

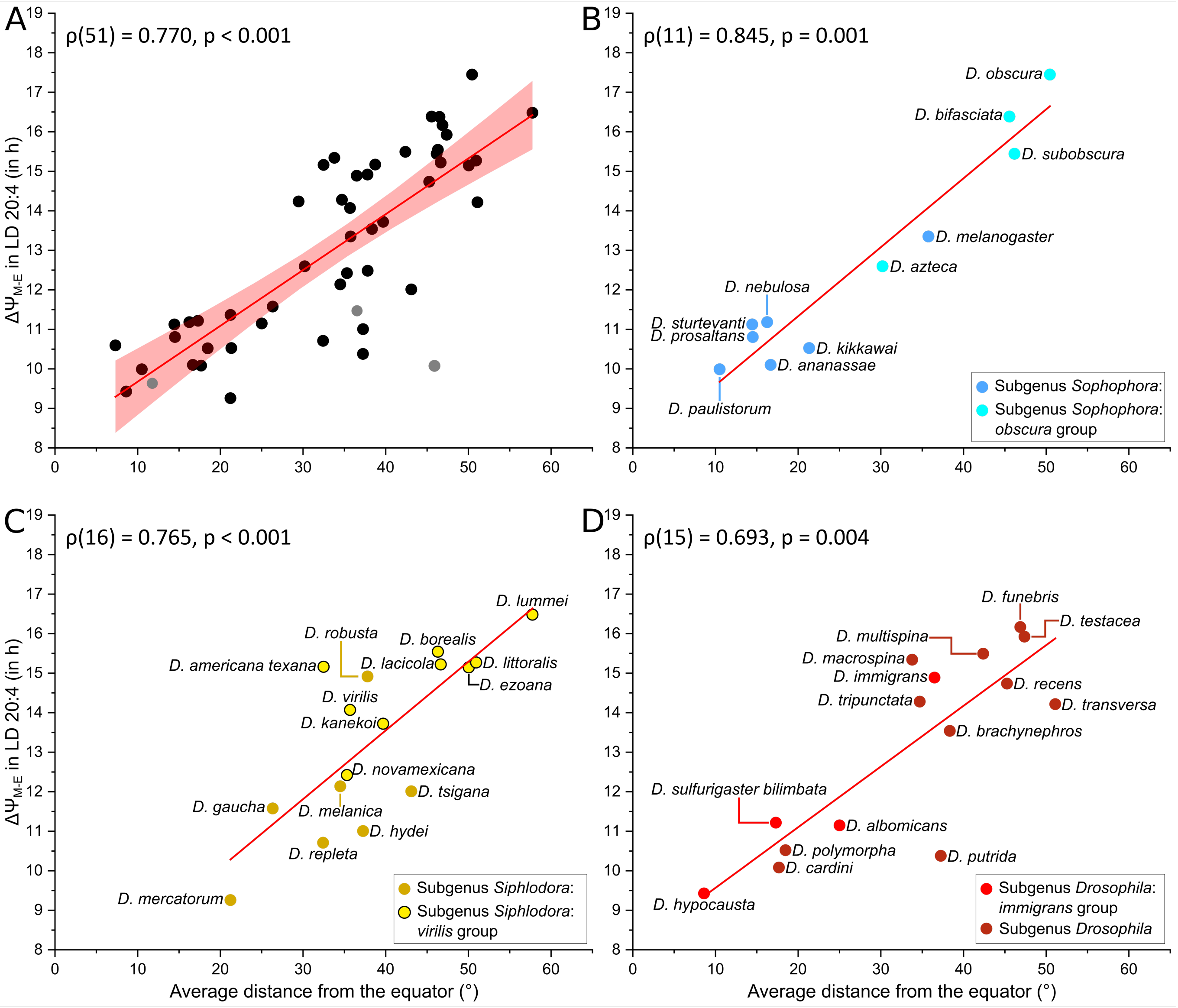

**Figure 6.** Correlation between $\Delta\Psi_{M\text{-}E}$ in LD 20:4 and average latitudinal distance from the equator in all *Drosophila* species **(A)** and the three different *Drosophila* subgenera **(B-D)** (trimmed by 3 % to exclude potential outlier reports, i.e., false extreme latitudinal values). To avoid distorting the correlation coefficient with duplicate values for species with multiple tested strains, only one strain per species is included (51 species total): *D. subobscura* from Bavaria, *S. pallida* collected in 2020, *D. repleta* from Bavaria (strain S), *D. virilis* from Rwanda, *D. funebris* from Alaska, and *D. macrospina* from Arizona. The three *Scaptodrosophila* species, which fall outside the main *Drosophila* radiation (see Figure 2), are shown in grey. **(B)** *Sophophora* subgenus, **(C)** *Drosophila* subgenus, **(D)** *Siphlodora* subgenus; Regression lines are indicated in red. The 95 % confidence band is illustrated in **(A)**. Spearman's rank correlation coefficients are indicated in the upper left corner of each panel.

#### Figure 7: CRY expression in l-LN$_v$ and correlation of CRY expression with $\Delta\Psi_{M\text{-}E}$

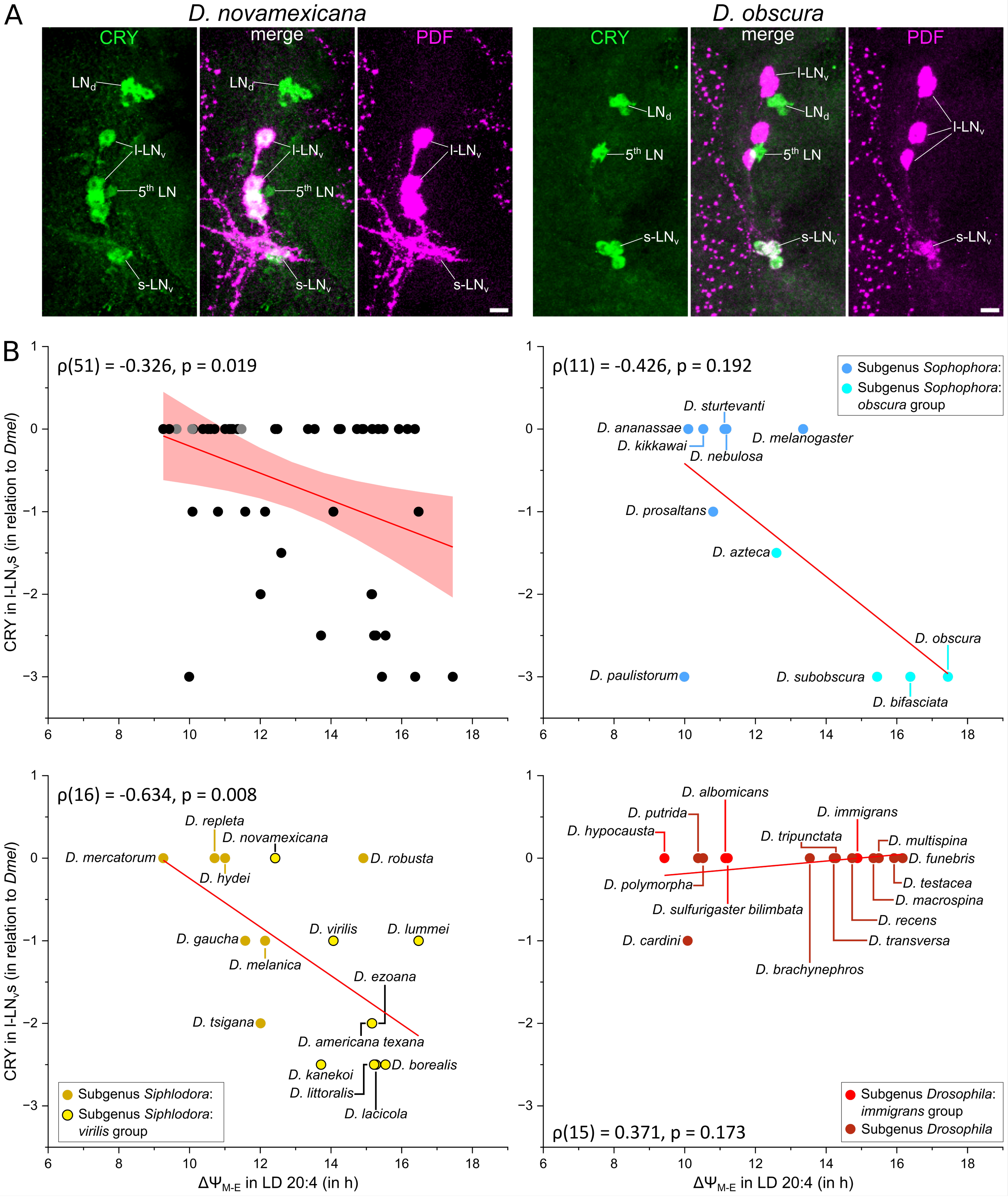

**Figure 7.** CRY expression in the l-LN$_v$ of two *Drosophila* species and correlation of CRY expression with $\Delta\Psi_{M\text{-}E}$ across species. **(A)** Confocal images of the lateral clock neurons of *D. novamexicana* (subgenus *Siphlodora) and D. obscura* (subgenus *Sophophora*), immunostained with anti-CRY (green) and anti-PDF (magenta); middle panels show an overlay of both channels. The dorsolateral neurons (LN$_d$) and the fifth lateral neuron (5$^{th}$ LN) belong to the evening neurons, while the small ventrolateral neurons (s-LN$_v$) are morning neurons. Note that *D. novamexicana* retains CRY in the four l-LN$_v$, while *D. obscura* lacks CRY in these neurons. Scale bar: 10 µm. **(B)** Correlation between CRY presence in the l-LN$_v$s and average $\Delta\Psi_{M\text{-}E}$ in all *Drosophila* species and within three main subgenera. The same 51 strains as in Figure 6 are included here. Labeling as in Figure 6.

**Figure 8: PDF expression pattern in the left brain hemisphere of selected species**

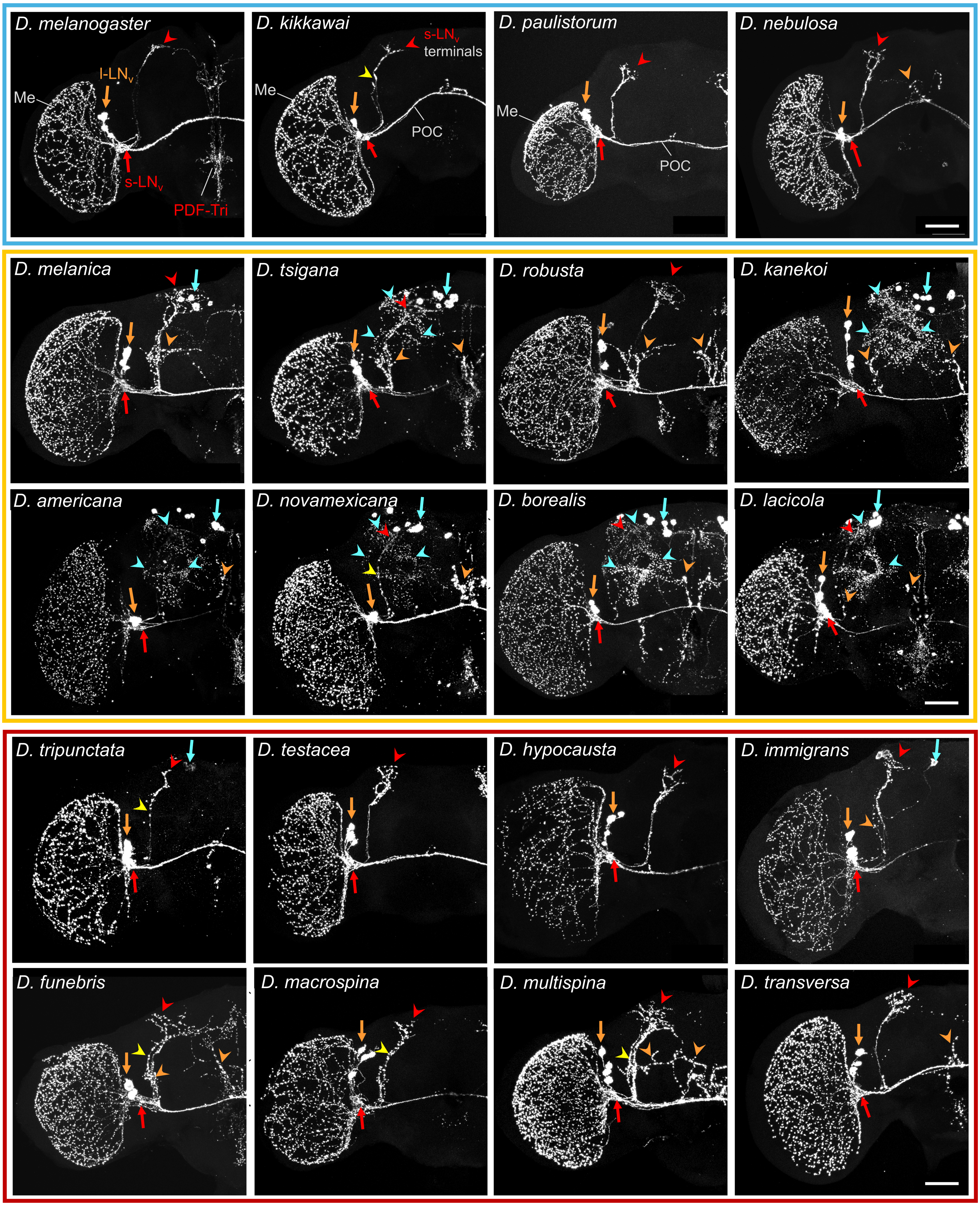

**Figure 8.** PDF expression pattern in the left brain hemisphere of selected species of the *Sophophora* (blue frame), *Siphlodora* (yellow frame) and *Drosophila* (red frame) subgenera. Orange arrows point to the l-LNv; orange arrowheads indicate l-LNv fibers leaving the posterior optic commissure (POC) and projecting into the centrolateral brain. Red arrows mark the s-LNv; red arrowheads mark their dorsal terminals. Yellow arrowheads indicate varicose projections of the s-LNv toward the dorsal brain. Cyan arrows mark putative PDF-positive corazonin cell clusters; cyan arrowheads mark their projections in the brain. Scale bar: 50 µm.

#### Figure 9: Correlations between PDF in the centrolateral brain area and $\Delta\Psi_{M-E}$

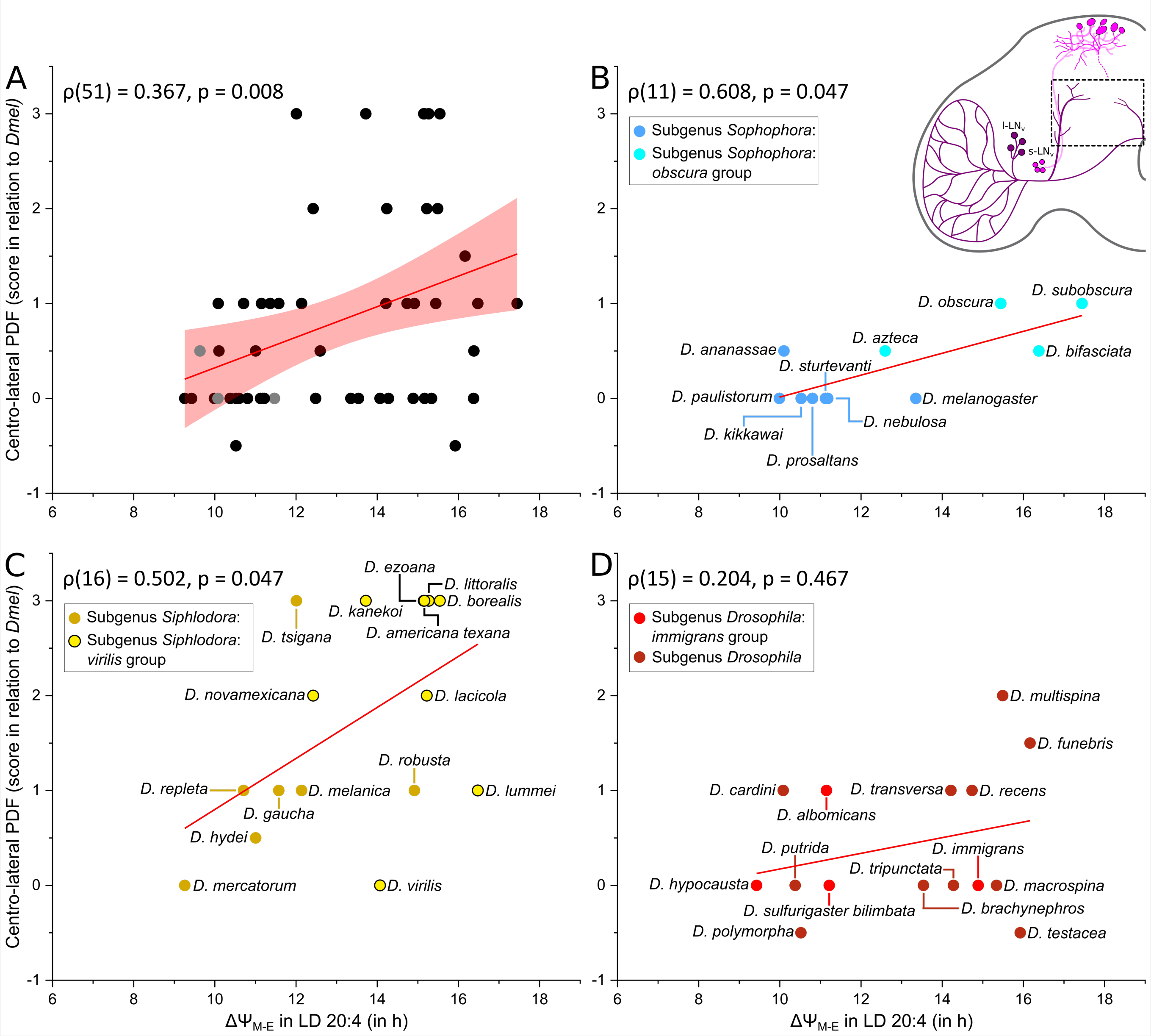

**Figure 9.** Correlations between the relative abundance of PDF in the centrolateral brain area and $\Delta\Psi_{M-E}$ in all *Drosophila* species **(A)** and within the three different *Drosophila* subgenera **(B-D)**. The inset in the upper right corner indicates the quantified brain region. The same 51 strains as in Figure 6 are included here. **(B)** *Sophophora* subgenus, **(C)** *Drosophila* subgenus, **(D)** *Siphlodora* subgenus. Labeling as in Figures 6 and 7.

**Figure 10: Correlations between PDF in *Siphlodora* and *Sohphophora***

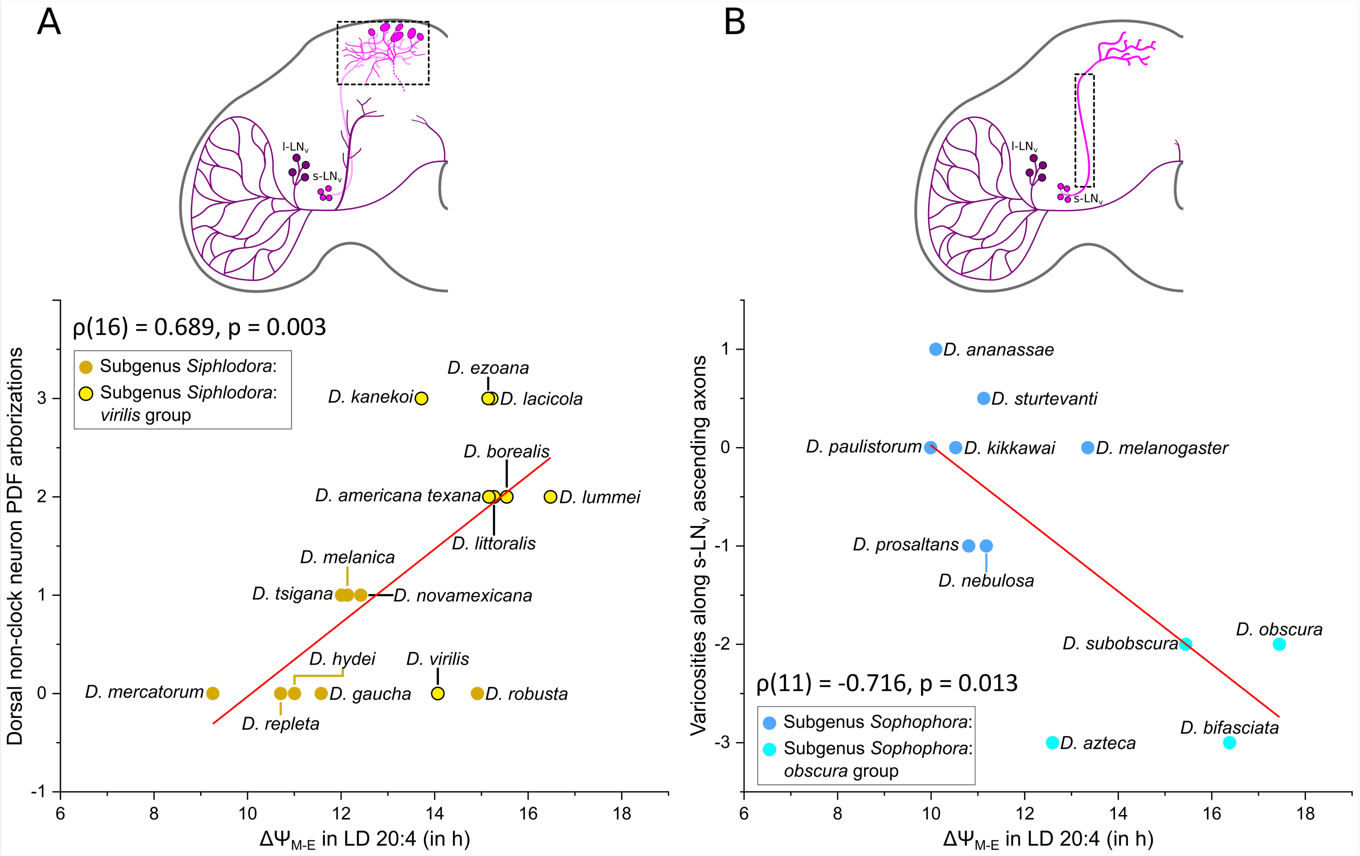

**Figure 10.** Correlations between the relative abundance of PDF in the dorsal protocerebrum and average $\Delta\Psi_{M\text{-}E}$ within the *Siphlodora* subgenus **(A)**, and between the PDF abundance in the ascending s-$LN_v$ axons and average $\Delta\Psi_{M\text{-}E}$ within the *Sophophora* subgenus **(B).**

**Figure 11: Correlation between PDF in the s-LNᵥ terminals and ΔΨ$_{M-E}$**

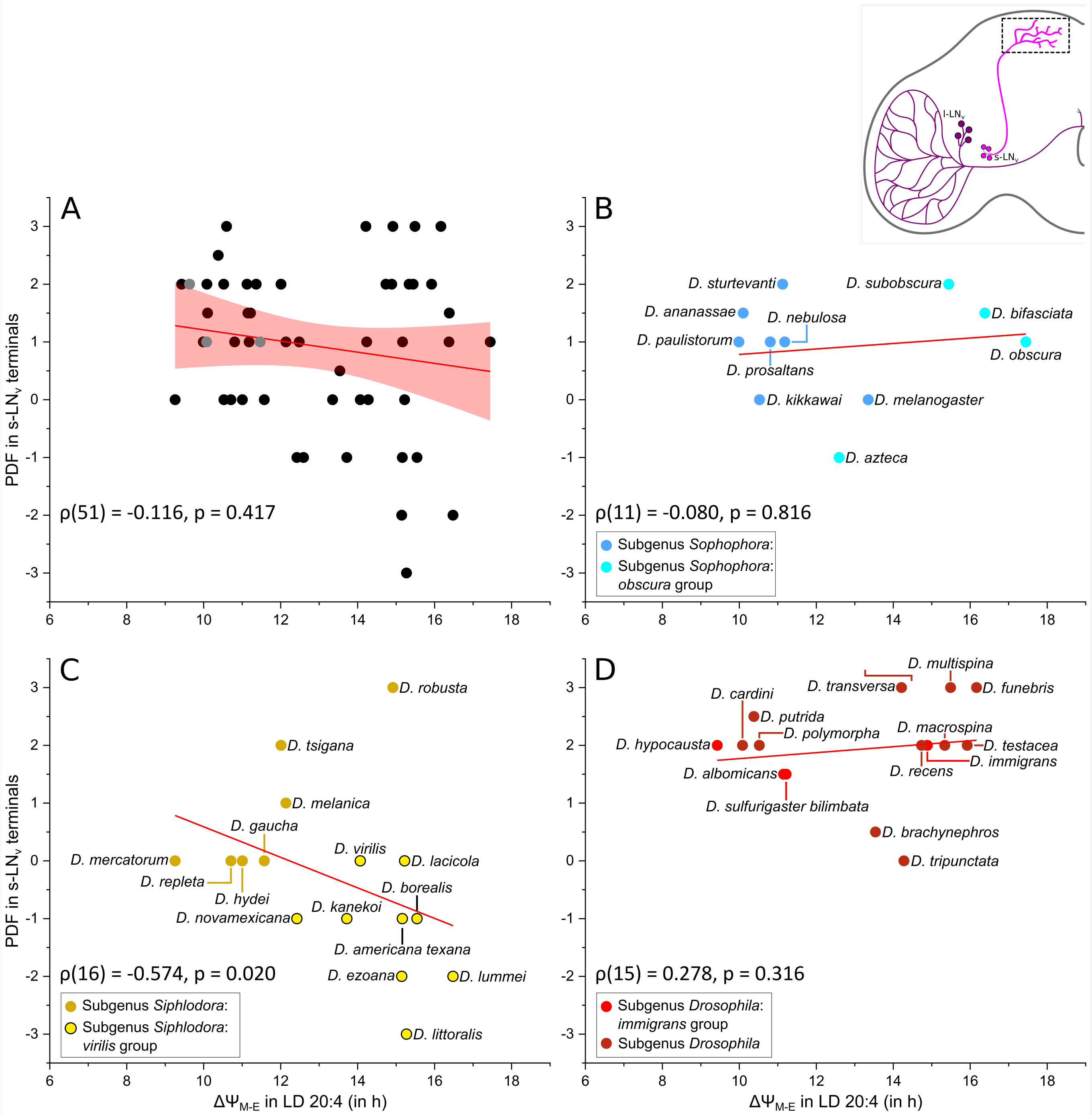

**Figure 11.** Correlation between PDF in the s-LNᵥ terminals and ΔΨ$_{M-E}$, shown across all *Drosophila* species **(A)** and within the three main subgenera **(B–D)**. PDF staining intensity showed no clear relationship with ΔΨM-E in the *Sophophora* subgenus **(B),** whereas it correlated negatively with ΔΨ$_{M-E}$ in the *Siphlodora* subgenus **(C)** and trended positively in the *Drosophila* subgenus **(D)**. The inset in the upper right corner indicates the quantified brain region. Labeling as in Figure 6.

**Figure 12: Summary of the correlations between PDF staining and latitude / $\Delta\Psi_{M\text{-}E}$**

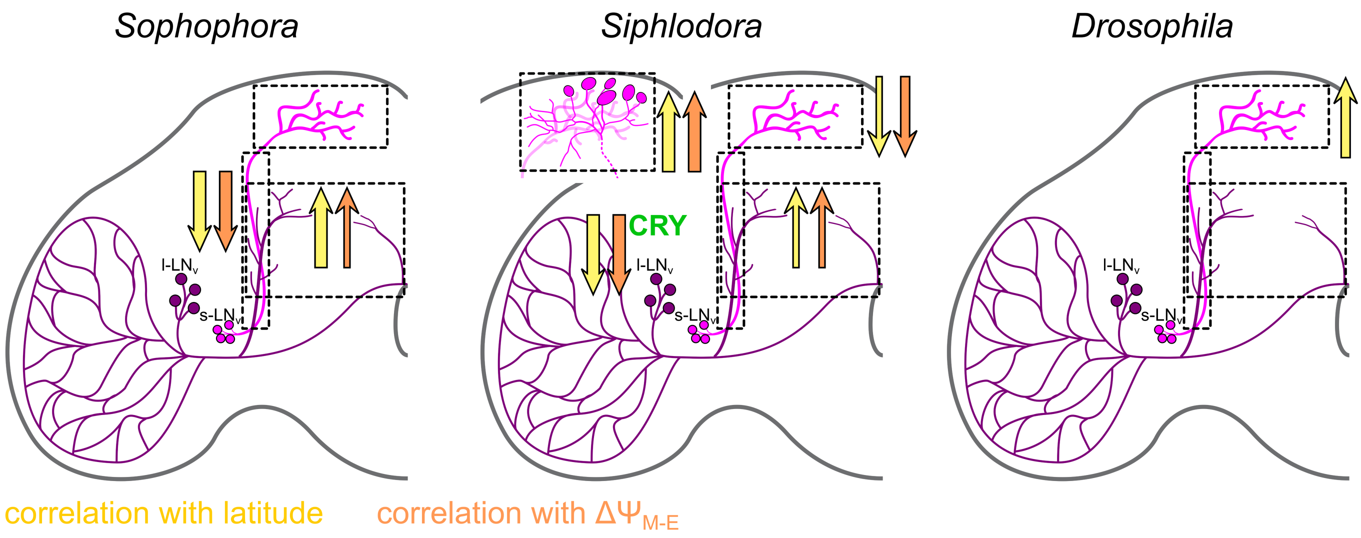

**Figure 12.** Summary of the correlations between PDF staining (magenta) and latitude/ $\Delta\Psi_{M\text{-}E}$ across the three different *Drosophila* subgenera. Arrow thickness indicates correlation strength; arrow direction indicates the direction of the correlation. Yellow arrows show correlations between PDF and latitude; orange arrows show correlations between PDF and $\Delta\Psi_{M\text{-}E}$. Within the *Siphlodora* subgenus, the expression of CRY in the l-$LN_v$ neurons also correlates negatively with $\Delta\Psi_{M\text{-}E}$ and latitude.

#### Figure 13: Power of rhythmicity in DD as a function of latitudinal distribution

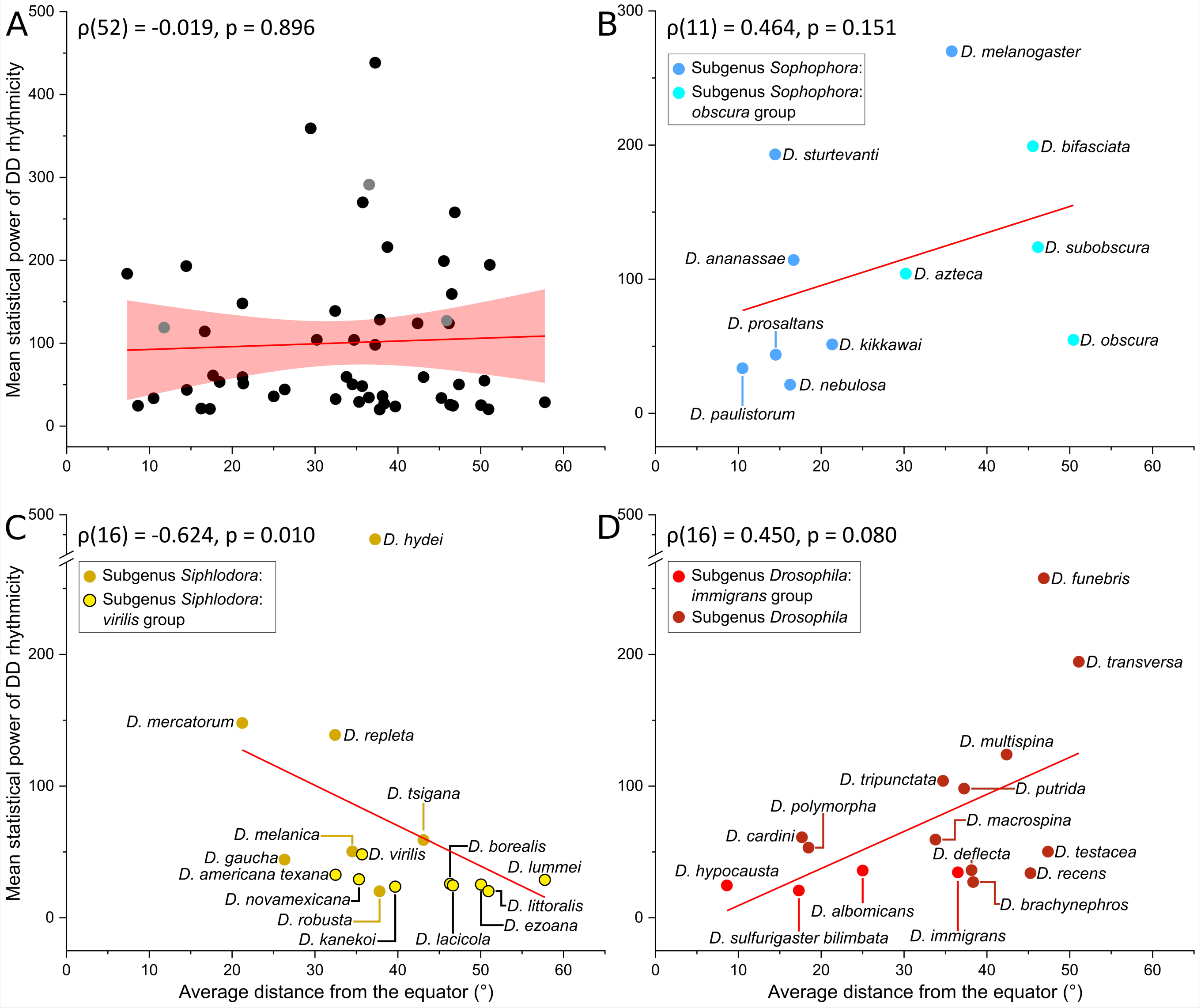

**Figure 13.** Power of rhythmicity in DD as a function of the mean latitudinal distribution (trimmed by 3 % to exclude potential outlier reports, i.e., false extreme latitudinal values). Spearman's rank correlation coefficient (ρ) is given in each panel. **(A)** Geographic range does not influence rhythm power across Drosophilidae as a whole. **(B)** and **(D)** No significant correlations between species range and rhythm strength were found in the subgenera *Sophophora* and *Drosophila*. **(C)** Within the *Siphlodora* subgenus, there was a significant negative relationship between latitudinal distribution and rhythm power.

**Figure 14: Power of rhythmicity in DD as a function of PDF expression features**

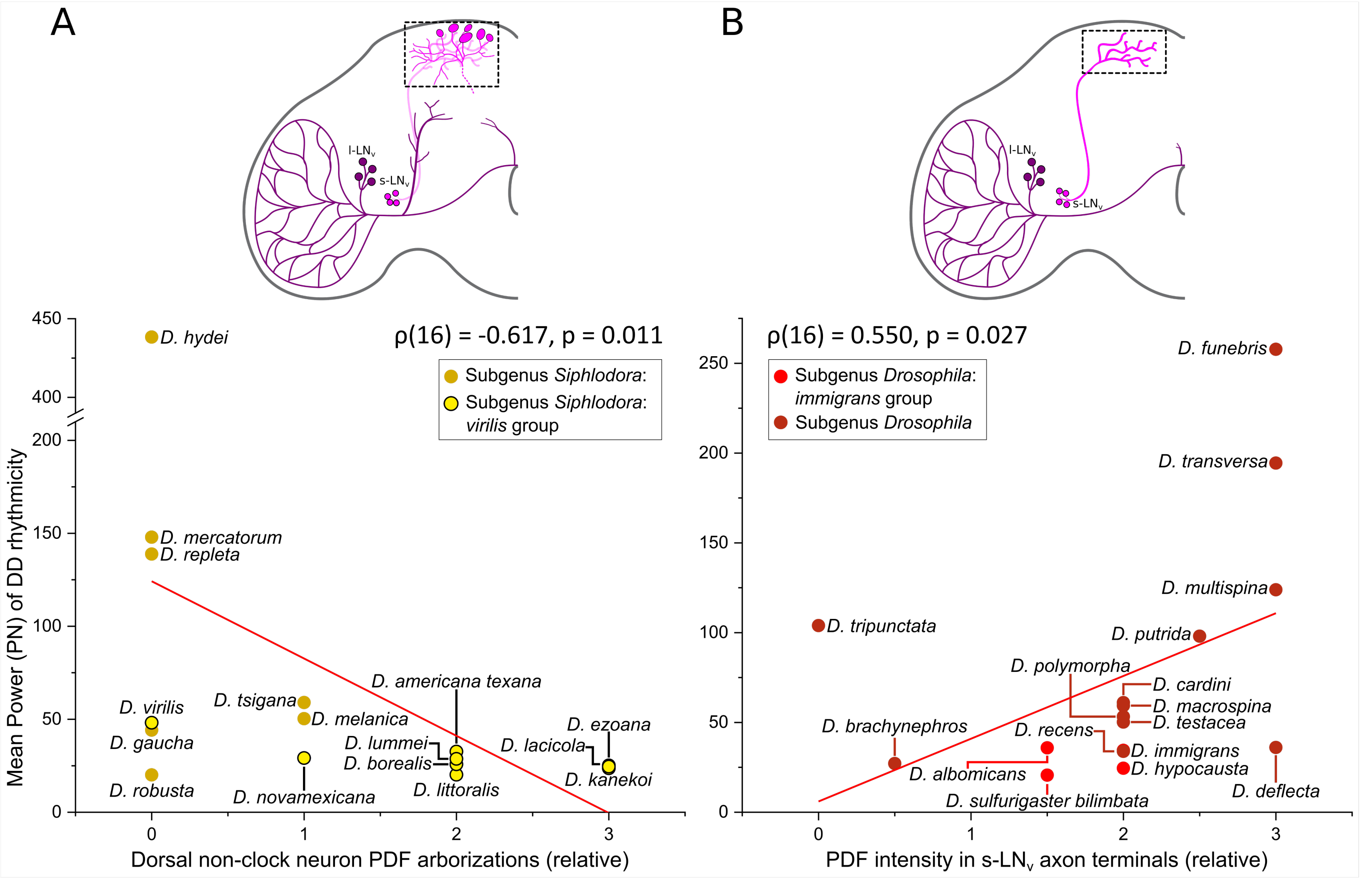

**Figure 14.** Power of rhythmicity in DD as a function of PDF expression features in the superior protocerebrum of the *Siphlodora* and *Drosophila* subgenera. Spearman's rank correlation coefficient (ρ) is given in each panel. The insets at the top indicate the quantified brain region. **(A)** A moderate negative correlation between mean power and the PDF intensity in dorsal non-clock CRZ neurons in the *Siphlodora* subgenus. **(B)** A moderate correlation between mean power and PDF intensity in the s-LNv terminals within the *Drosophila* subgenus.
