## Supplementary figures for "Fruit flies of the subgenera *Drosophila*, *Siphlodora* and *Sophophora* use different mechanisms to adapt their daily activity to long photoperiods"

bioRxiv, 2026

**Supplementary figures**

**Figure 3—figure supplements**

➔ See next pages

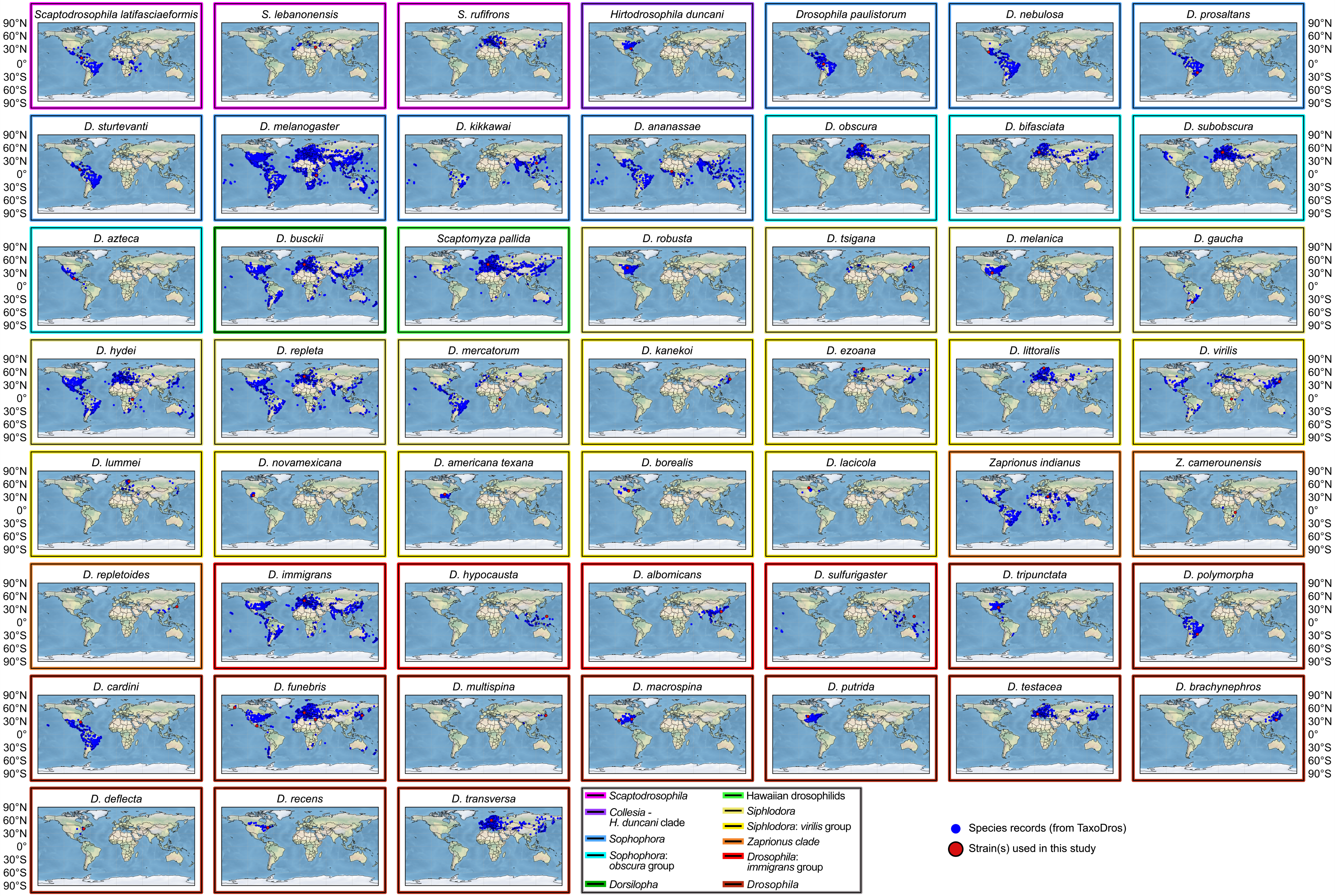
Scaptodrosophila latifasciaeformis
S. lebanonensis
S. rufifrons
Hirtodrosophila duncani
Drosophila paulistorum
D. nebulosa
D. prosaltans
D. sturtevanti
D. melanogaster
D. kikkawai
D. ananassae
D. obscura
D. bifasciata
D. subobscura
D. azteca
D. busckii
Scaptomyza pallida
D. robusta
D. tsigana
D. melanica
D. gaucha
D. hydei
D. repleta
D. mercatorum
D. kanekoi
D. ezoana
D. littoralis
D. virilis
D. lummei
D. novamexicana
D. americana texana
D. borealis
D. lacicola
Zaprionus indianus
Z. camerounensis
D. repletoides
D. immigrans
D. hypocausta
D. albomicans
D. sulfurigaster
D. tripunctata
D. polymorpha
D. cardini
D. funebris
D. multispina
D. macrospina
D. putrida
D. testacea
D. brachynephros
D. deflecta
D. recens
D. transversa
90°N
60°N
30°N
0°
30°S
60°S
90°S
Scaptodrosophila
Collesia - H. duncani clade
Sophophora
Sophophora: obscura group
Dorsilopha
Hawaiian drosophilids
Siphlodora
Siphlodora: virilis group
Zaprionus clade
Drosophila: immigrans group
Drosophila
Species records (from TaxoDros)
Strain(s) used in this study

**Figure 3—figure supplement 1.** Geographical distribution of the species (from TaxoDros) including the origin of the strains used in the present study (red dots).

**Extended legend:**
Within genus *Scaptodrosophila*, *S. latifasciaeformis* is a true equatorial species, whereas *S. lebanonensis* is mostly distributed across arid, temperate regions of the Mediterranean and the Middle East. *S. rufifrons* is restricted to the temperate Palearctic, with its northernmost findings coming from southern Sweden and Siberia. Within the subgenus *Sophophora*, which also includes the model organism *D. melanogaster*, distribution patterns differ clearly among species groups. Representatives of the *willistoni* group (*D. nebulosa*, *D. paulistorum*) and the *saltans* group (*D. prosaltans*, *D. sturtevanti*) are neotropic and do not occur beyond the 35$^{th}$ parallels of the New World. *D. ananassae* and *D. kikkawai*, two *melanogaster* group species originally from the Old World, are likewise restricted to equatorial regions but have since spread to America, Asia, Australia, and Oceania, making them cosmopolitan species of the tropics. *D. melanogaster* itself is a true cosmopolitan, occurring even in temperate regions. In contrast to these seven species of the *willistoni*, *saltans* and *melanogaster* groups, only *D. azteca* among the four *obscura* group species occurs in tropical climate zones; the other three, *D. bifasciata*, *D. obscura*, and *D. subobscura* are restricted to temperate and subarctic regions, with *D. bifasciata* and *D. obscura* extending as far north as Scandinavia. *Hirtodrosophila duncani* belongs to an outgroup of the main *Sophpohora* subgenus (see Figure 2) and is restricted to the eastern United States. *D. busckii*, the sole representant of the subgenus *Dorsilopha*, is native to North America but is now found worldwide, having spread into tropical and temperate regions, with recorts also documented in southern Scandinavia.

The main *Drosophila* cluster is divided into the subgenera *Siphlodora* and *Drosophila*, with the Hawaiian clade and the *Zaprionus* clade positioned outside the main subgenera (see Figure 2). Of the Hawaiian clade, we investigated only *Scaptomyza pallida,* which has spread far north, occurring at the North Cape, in Iceland, and in northern Siberia. Of the two species from the *Zaprionus* subgenus, *Z. camerounensis* is limited to equatorial Africa, while *Z. indianus* is a tropical cosmopolite. The phylogenetic position of *D. repletoides* remains controversial: recent studies allocate them with the *Zaprionus* clade (Russo et al., 2013; Yassin, 2013), although its geographic distribution is restricted to East and Southeast Asia.

The subgenus *Siphlodora* is composed of species that belong to the *virilis-repleta* radiation (O'Grady and DeSalle, 2018). Most species investigated belong to the *virilis* group: *D. kanekoi* is limited to East Asia (China, Japan); *D. borealis* and *D. lacicola* are native to the northern United States and Canada; *D. americana texana* and *D. novamexicana* occur in more southern, arid regions of the US; *D. virilis* is found worldwide, mainly in temperate but occasionally tropical regions (two strains of *D. virilis* were tested; red dots in the map); and *D. ezoana*, *D. littoralis*, and *D. lummei* are Palearctic species found mostly in northern Europe, Siberia, and northern Japan. The investigated *repleta* group species *D. hydei*, *D. mercatorum*, and *D. repleta* all share a cosmopolitan distribution, though most records — particularly for *D. mercatorum* — come from tropical regions. *D. gaucha*, from the *mesophragmatica* subgroup, has a well-defined range, occuring exclusively on South American plains and thus basically restricted to the southern hemisphere. The remaining *Siphlodora* representatives from the *melanica* group (*D. melanica*, *D. tsigana*) and the *robusta* group (*D. robusta*) share a common latitudinal range of 30°N – 50°N.

Geographical distribution of species within the *Drosophila* subgenus is highly variable. *D. sulfurigaster* and *D. hypocausta* from the *immigrans* group are tropical species home to Southeast Asia and the Pacific region, while *D. albomicans* occurs somewhat further north, mainly in China and Japan. *D. immigrans*, by contrast, has spread all over the world and is also found in temperate regions and few boreal areas. Two species belong to the *testacea* group: *D. putrida* is restricted to North America, especially the eastern US, whereas *D. testacea* also occurs in the Palearctic, extending to Scandinavia. The mushroom-feeding flies of the *quinaria* group are largely restricted to temperate and boreal regions of North America (*D. recens*, *D. deflecta*) or to Japan and the Korean Peninsula (*D. brachynephros*), while *D. transversa* is a Nearctic and Palearctic cosmopolite capable of reaching far northern latitudes (68°N). Species from the *cardini* group, *D. cardini* and *D. polymorpha*, are restricted to the Neotropics, and *D. tripunctata*, from the *tripunctata* group, occurs in North and Central America. The final group within the *Drosophila* subgenus is the *funebris* group: *D. macrospina* is distributed mainly in the southern United States and Mexico, while *D. multispina* is native to East Asia, especially Japan. *D. funebris* is a cosmopolite, found predominantly in temperate regions and, noteworthy, in arctic areas (Alaska, Iceland, Svalbard), but occasionally also in tropical locations. Four strains of *D. funebris* and two strains of *D. macrospina* were tested (red dots).

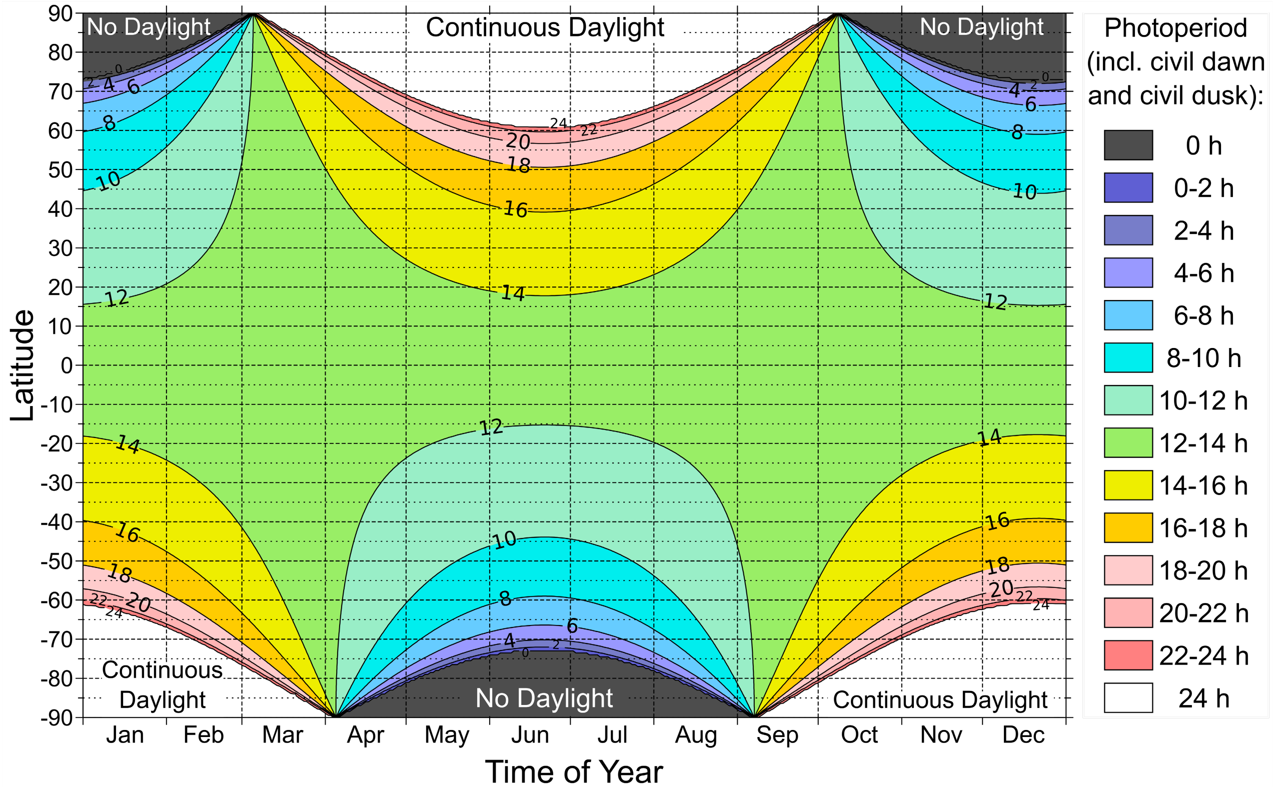

**Figure 3—figure supplement 2.** Variation of photoperiod in dependence of latitude and season. The light phases in this diagram include the civil twilight phases before sunrise and after sunset. At civil twilight, the sun is 6° below the horizon. Note that the light phases in the northern and southern hemispheres are not identical and that the length of the day (including civil twilight) is longer than 12 hours in summer, even at the equator (Modified from an original illustration of Stephen Hudson).

### Figure 5—figure supplements

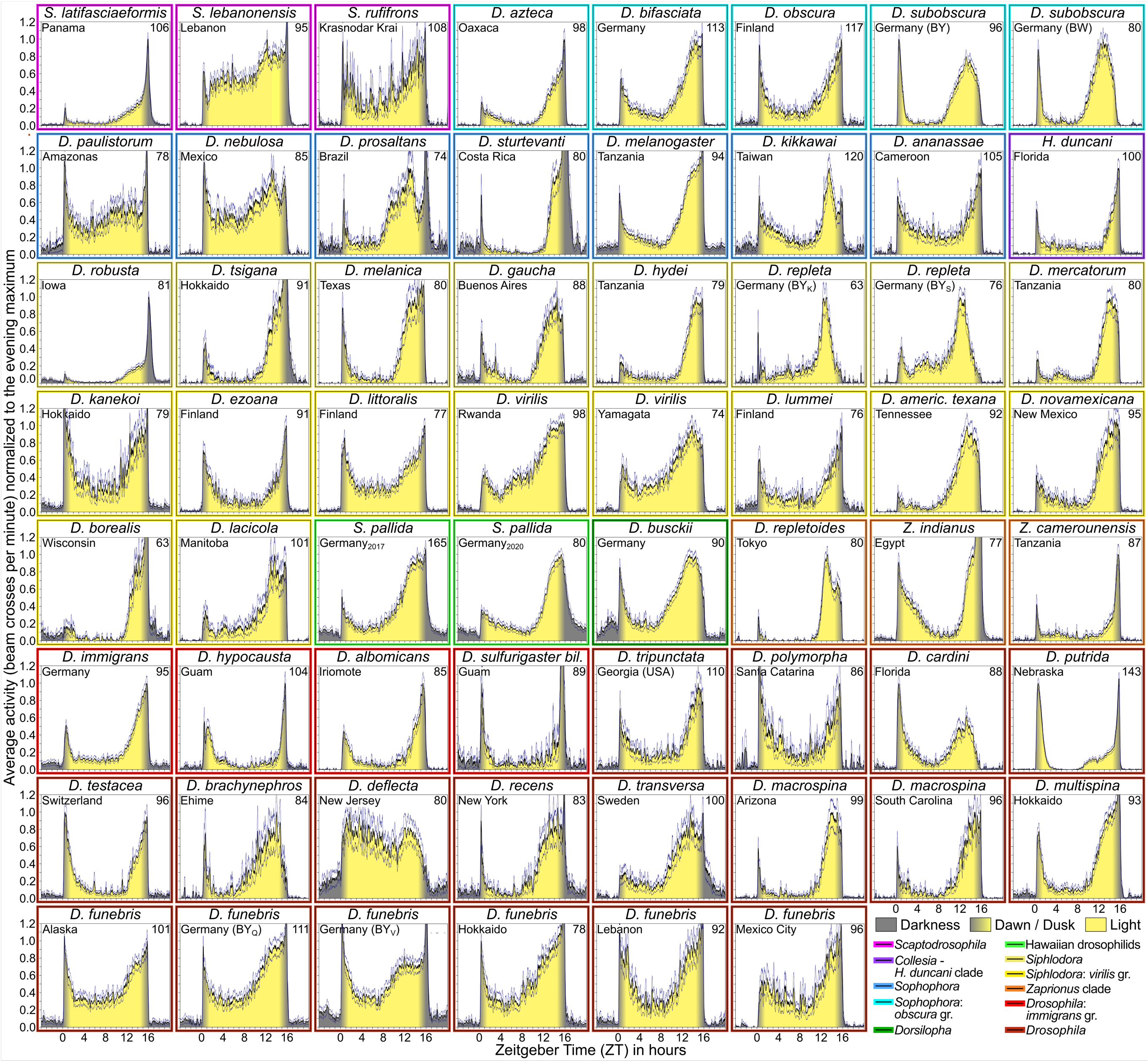

**Figure 5—figure supplement 1.** Average daily activity profiles of all tested species and strains in LD 16:8. Curves are normalized to the maximum amplitude of the circadian-driven morning or evening peaks. The origin of the species (or strain) is indicated in the upper left corner of each diagram, and the number of flies included is indicated in the upper right corner. Frame colors refer to the phylogenic color-coding shown in the lower right from Figure 2 (also shown in the lower right corner). During normalization, maximal activity during the circadian evening peak was set to 1. This was done to improve visualization, since some species were very inactive. The magnitude of startle responses to dawn and dusk events was disregarded (e.g., the lights-off startle response was unusually high and long-lasting in *D. sturtevanti*, second row).

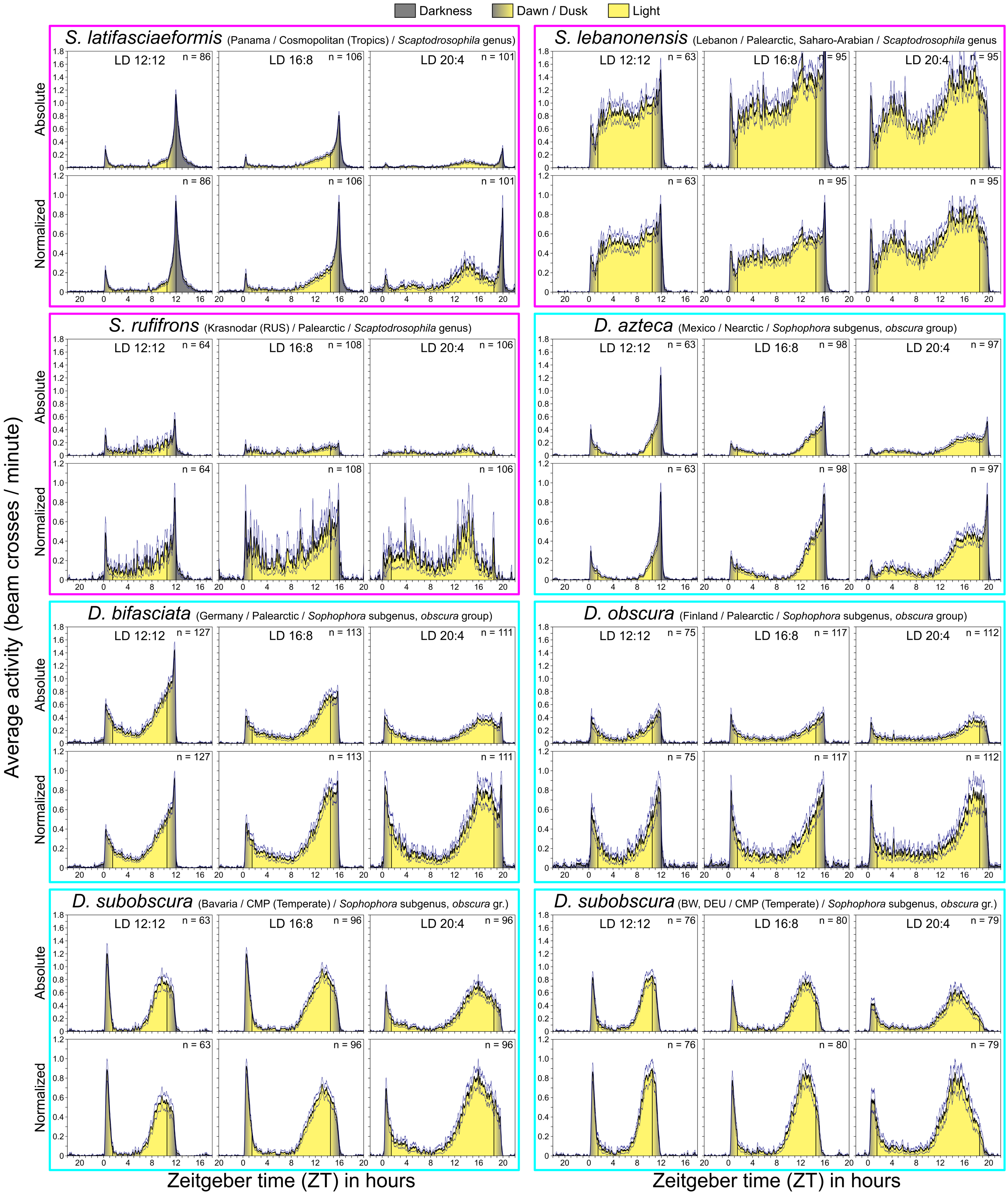

**Figure 5—figure supplement 2.** Compilation of average activity profiles in LD 12:12, LD 16:8, and LD 20:4 for all strains of the *Scaptodrosophila* genus (magenta) and the *obscura* group within the *Sophophora* subgenus (cyan). Shown are the average day profiles calculated out of absolute values, and the profiles normalized to the highest average activity, often occurring at the lights-off event. Thin blue curves indicate the standard error. Information about the strain origin, the current geographical distribution, and the phylogeny (from left to right) is stated after the species name in parentheses.

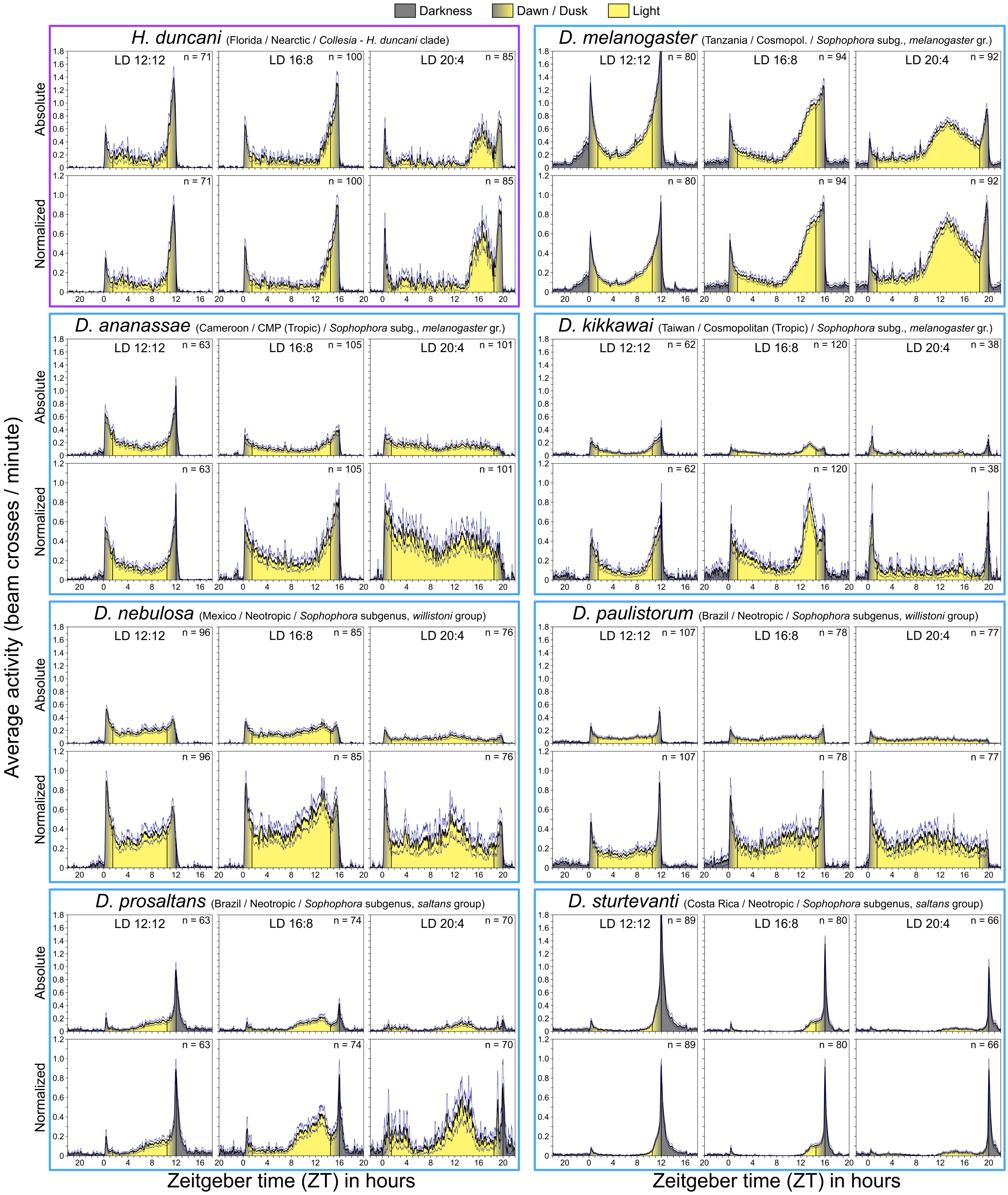

**Figure 5—figure supplement 3.** Compilation of average activity profiles in LD 12:12, LD 16:8, and LD 20:4 for the remaining the *Sophophora* strains (blue) and *H. duncani* (purple). Shown are the average day profiles calculated out of absolute values, and the profiles normalized to the highest average activity, often occurring at the lights-off event. Thin blue curves indicate the standard error. Information about the strain origin, the current geographical distribution, and the phylogeny (from left to right) is stated after the species name in parentheses.

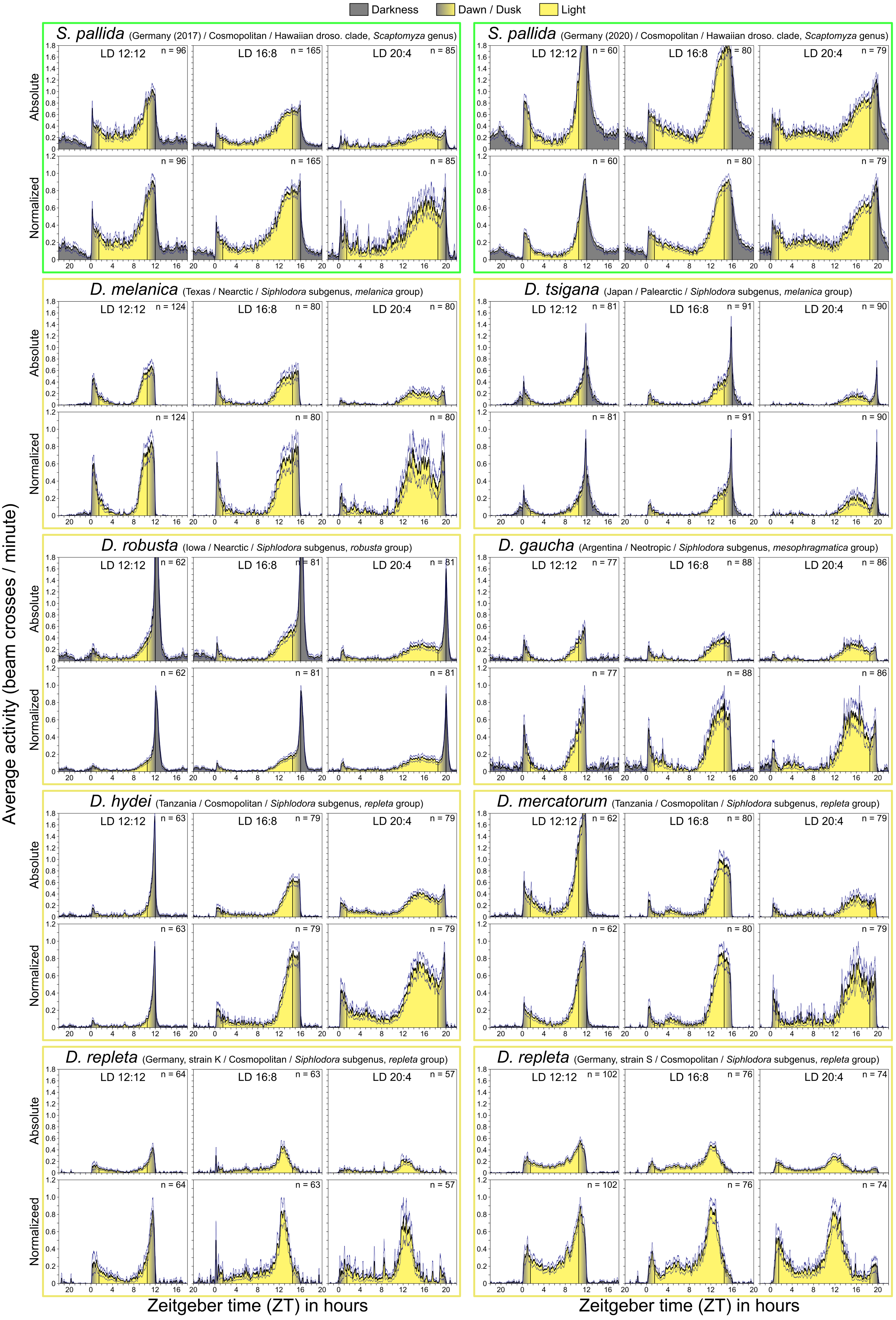

Darkness
Dawn / Dusk
Light
S. pallida (Germany (2017) / Cosmopolitan / Hawaiian droso. clade, Scaptomyza genus)
S. pallida (Germany (2020) / Cosmopolitan / Hawaiian droso. clade, Scaptomyza genus)
D. melanica (Texas / Nearctic / Siphlodora subgenus, melanica group)
D. tsigana (Japan / Palearctic / Siphlodora subgenus, melanica group)
D. robusta (Iowa / Nearctic / Siphlodora subgenus, robusta group)
D. gaucha (Argentina / Neotropic / Siphlodora subgenus, mesophragmatica group)
D. hydei (Tanzania / Cosmopolitan / Siphlodora subgenus, repleta group)
D. mercatorum (Tanzania / Cosmopolitan / Siphlodora subgenus, repleta group)
D. repleta (Germany, strain K / Cosmopolitan / Siphlodora subgenus, repleta group)
D. repleta (Germany, strain S / Cosmopolitan / Siphlodora subgenus, repleta group)
LD 12:12
LD 16:8
LD 20:4
Absolute
Normalized
Normalizeed
Average activity (beam crosses / minute)
Zeitgeber time (ZT) in hours

**Figure 5—figure supplement 4.** Compilation of average activity profiles in LD 12:12, LD 16:8, and LD 20:4 for all strains of the two *S. pallida* strains (light green) and strains from the *Siphlodora* subgenus (light gold). Shown are the average day profiles calculated out of absolute values, and the profiles normalized to the highest average activity, often occurring at the lights-off event. Thin blue curves indicate the standard error. Information about the strain origin, the current geographical distribution, and the phylogeny (from left to right) is stated after the species name in parentheses.

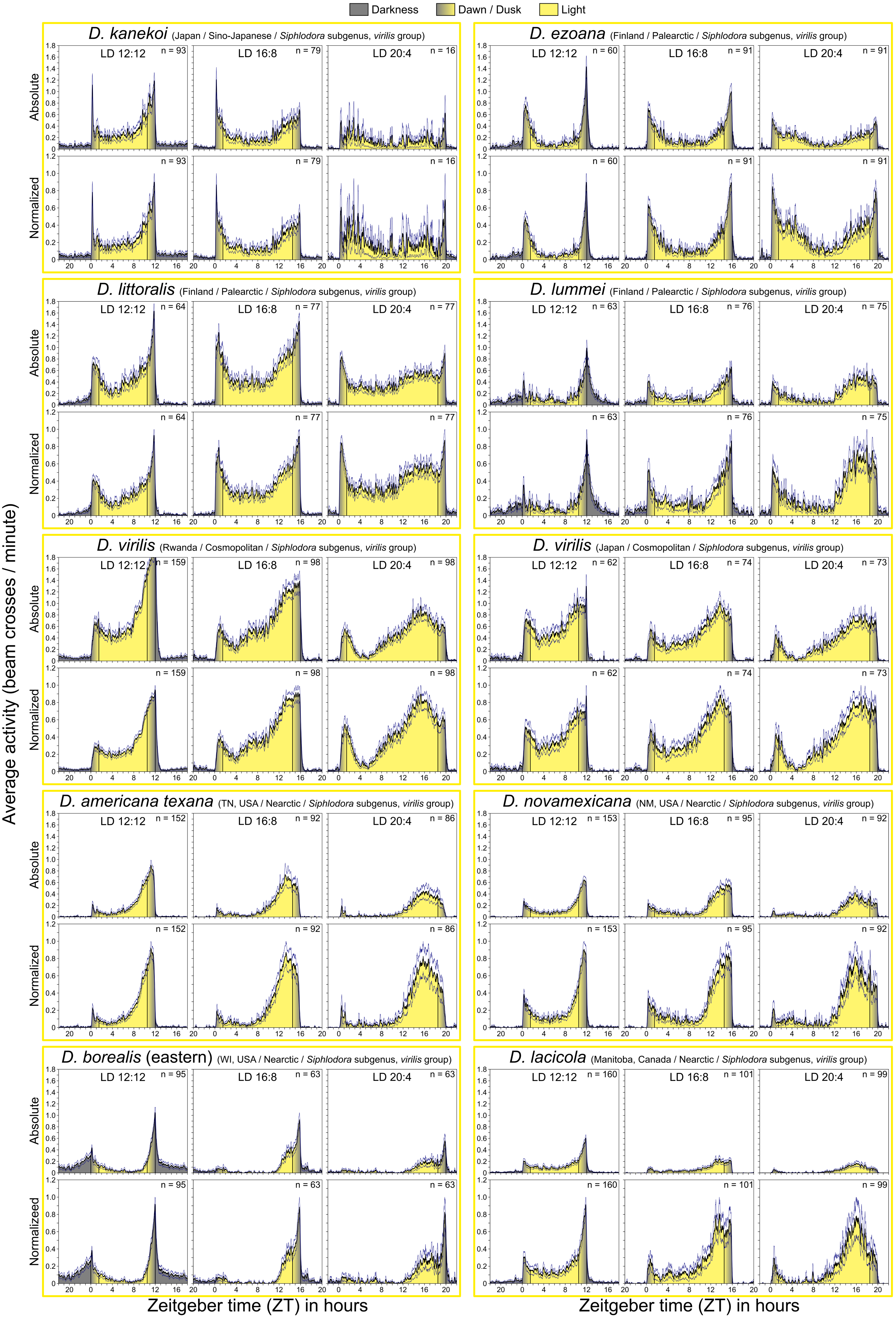

Darkness
Dawn / Dusk
Light
D. kanekoi (Japan / Sino-Japanese / Siphlodora subgenus, virilis group)
D. ezoana (Finland / Palearctic / Siphlodora subgenus, virilis group)
D. littoralis (Finland / Palearctic / Siphlodora subgenus, virilis group)
D. lummei (Finland / Palearctic / Siphlodora subgenus, virilis group)
D. virilis (Rwanda / Cosmopolitan / Siphlodora subgenus, virilis group)
D. virilis (Japan / Cosmopolitan / Siphlodora subgenus, virilis group)
D. americana texana (TN, USA / Nearctic / Siphlodora subgenus, virilis group)
D. novamexicana (NM, USA / Nearctic / Siphlodora subgenus, virilis group)
D. borealis (eastern) (WI, USA / Nearctic / Siphlodora subgenus, virilis group)
D. lacicola (Manitoba, Canada / Nearctic / Siphlodora subgenus, virilis group)
LD 12:12
LD 16:8
LD 20:4
Absolute
Normalized
Normalizeed
Average activity (beam crosses / minute)
Zeitgeber time (ZT) in hours

**Figure 5—figure supplement 5.** Compilation of average activity profiles in LD 12:12, LD 16:8, and LD 20:4 for all tested *virilis* group strains within the *Siphlodora* subgenus (yellow). Shown are the average day profiles calculated out of absolute values, and the profiles normalized to the highest average activity, often occurring at the lights-off event. Thin blue curves indicate the standard error. Information about the strain origin, the current geographical distribution, and the phylogeny (from left to right) is stated after the species name in parentheses.

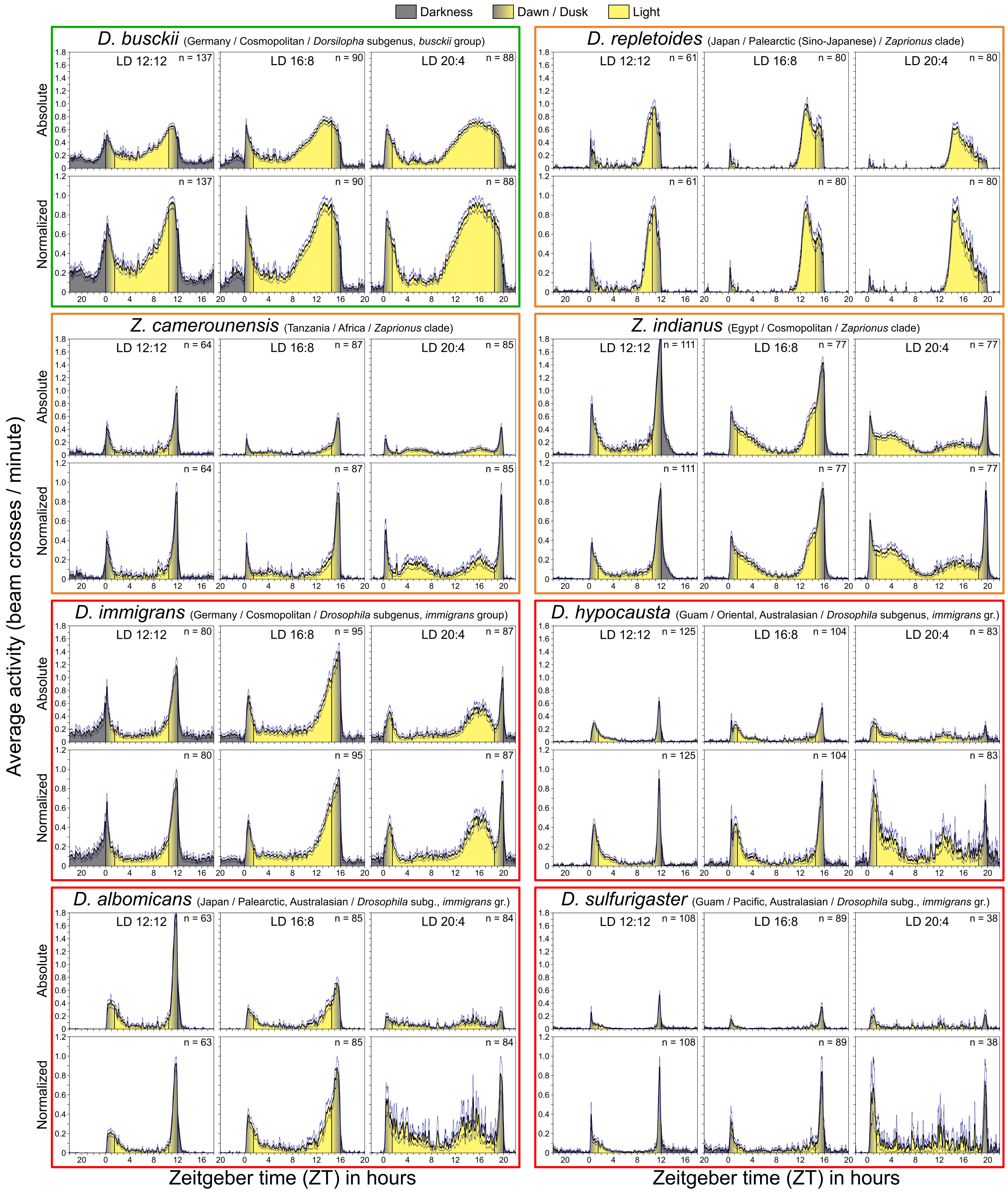

**Figure 5—figure supplement 6.** Compilation of average activity profiles in LD 12:12, LD 16:8, and LD 20:4 for all strains of the *Zaprionus* clade (orange), the *immigrans* species group (light red), and *D. busckii* (dark green). Shown are the average day profiles calculated out of absolute values, and the profiles normalized to the highest average activity, often occurring at the lights-off event. Thin blue curves indicate the standard error. Information about the strain origin, the current geographical distribution, and the phylogeny (from left to right) is stated after the species name in parentheses.

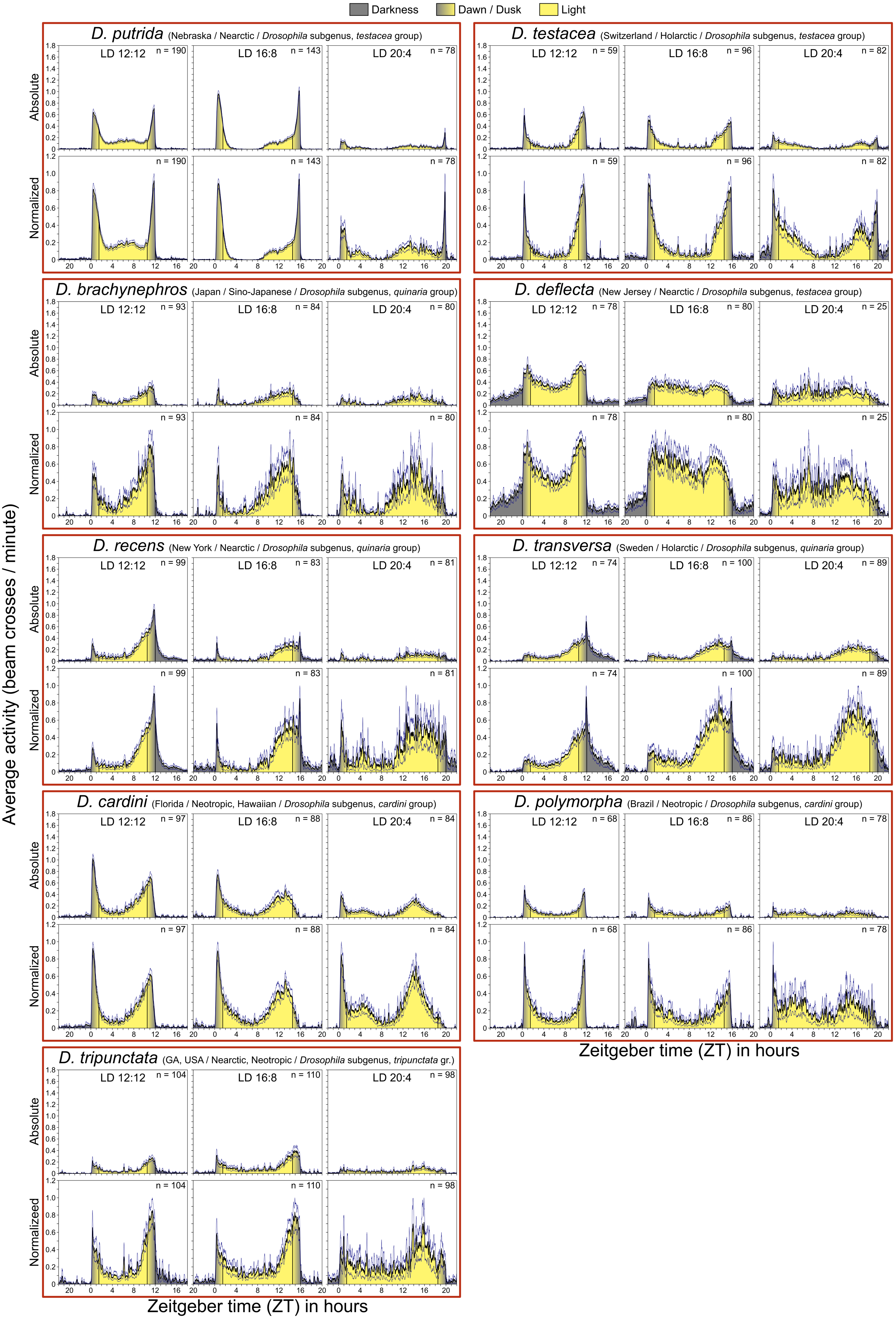

Darkness
Dawn / Dusk
Light
D. putrida (Nebraska / Nearctic / Drosophila subgenus, testacea group)
D. testacea (Switzerland / Holarctic / Drosophila subgenus, testacea group)
D. brachynephros (Japan / Sino-Japanese / Drosophila subgenus, quinaria group)
D. deflecta (New Jersey / Nearctic / Drosophila subgenus, testacea group)
D. recens (New York / Nearctic / Drosophila subgenus, quinaria group)
D. transversa (Sweden / Holarctic / Drosophila subgenus, quinaria group)
D. cardini (Florida / Neotropic, Hawaiian / Drosophila subgenus, cardini group)
D. polymorpha (Brazil / Neotropic / Drosophila subgenus, cardini group)
D. tripunctata (GA, USA / Nearctic, Neotropic / Drosophila subgenus, tripunctata gr.)
LD 12:12
LD 16:8
LD 20:4
Absolute
Normalized
Normalizeed
Average activity (beam crosses / minute)
Zeitgeber time (ZT) in hours

**Figure 5—figure supplement 7.** Compilation of average activity profiles in LD 12:12, LD 16:8, and LD 20:4 for strains of the *Drosophila* subgenus (red). Shown are the average day profiles calculated out of absolute values, and the profiles normalized to the highest average activity, often occurring at the lights-off event. Thin blue curves indicate the standard error. Information about the strain origin, the current geographical distribution, and the phylogeny (from left to right) is stated after the species name in parentheses.

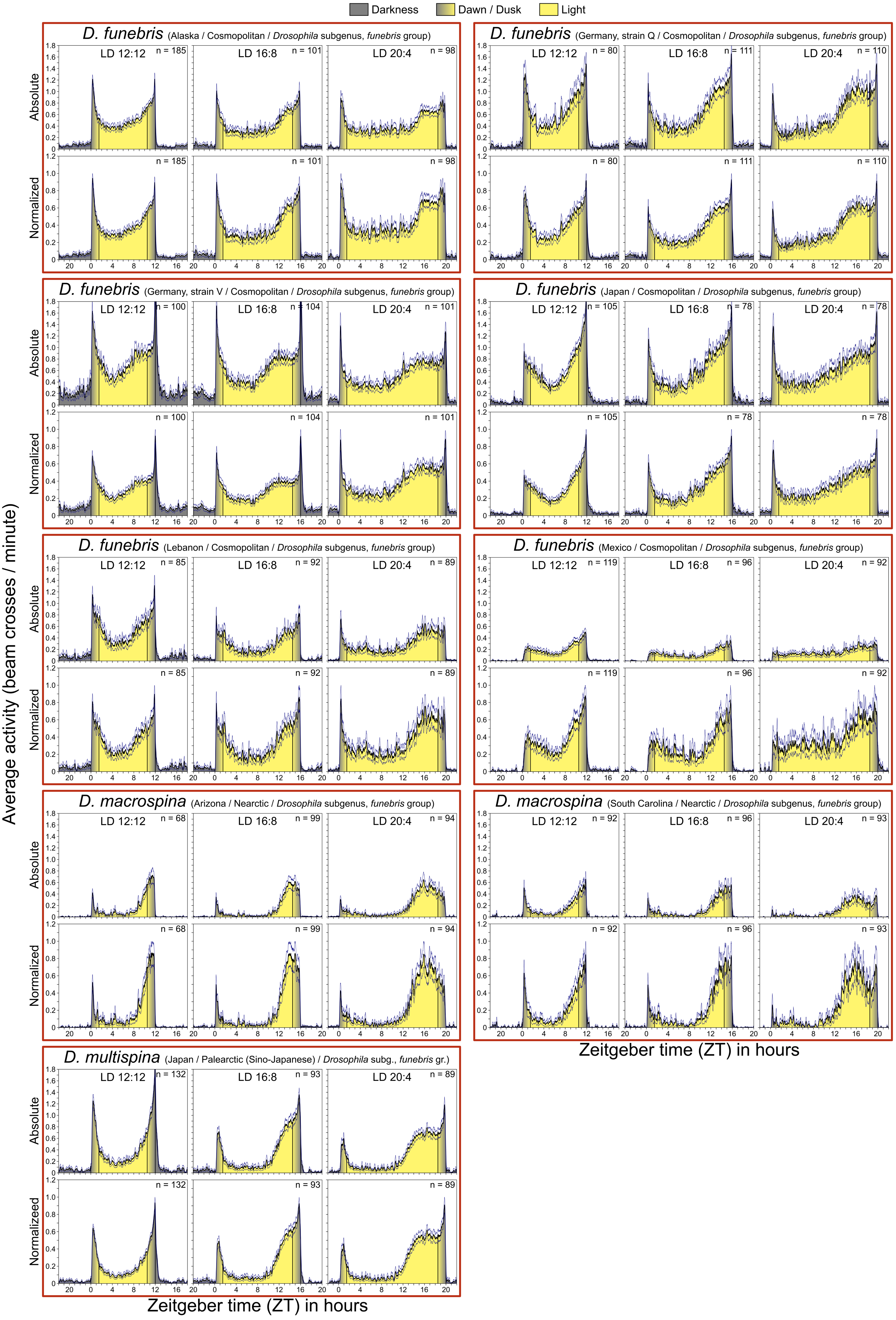

Darkness
Dawn / Dusk
Light
D. funebris (Alaska / Cosmopolitan / Drosophila subgenus, funebris group)
D. funebris (Germany, strain Q / Cosmopolitan / Drosophila subgenus, funebris group)
D. funebris (Germany, strain V / Cosmopolitan / Drosophila subgenus, funebris group)
D. funebris (Japan / Cosmopolitan / Drosophila subgenus, funebris group)
D. funebris (Lebanon / Cosmopolitan / Drosophila subgenus, funebris group)
D. funebris (Mexico / Cosmopolitan / Drosophila subgenus, funebris group)
D. macrospina (Arizona / Nearctic / Drosophila subgenus, funebris group)
D. macrospina (South Carolina / Nearctic / Drosophila subgenus, funebris group)
D. multispina (Japan / Palearctic (Sino-Japanese) / Drosophila subg., funebris gr.)
LD 12:12
LD 16:8
LD 20:4
Absolute
Normalized
Normalizeed
n = 185
n = 101
n = 98
n = 80
n = 111
n = 110
n = 100
n = 104
n = 101
n = 105
n = 78
n = 78
n = 85
n = 92
n = 89
n = 119
n = 96
n = 92
n = 68
n = 99
n = 94
n = 92
n = 96
n = 93
n = 132
n = 93
n = 89
Average activity (beam crosses / minute)
Zeitgeber time (ZT) in hours

**Figure 5—figure supplement 8.** Compilation of average activity profiles in LD 12:12, LD 16:8, and LD 20:4 for all strains of the *funebris* species group. Shown are the average day profiles calculated out of absolute values, and the profiles normalized to the highest average activity, often occurring at the lights-off event. Thin blue curves indicate the standard error. Information about the strain origin, the current geographical distribution, and the phylogeny (from left to right) is stated after the species name in parentheses.

### Figure 6—figure supplements

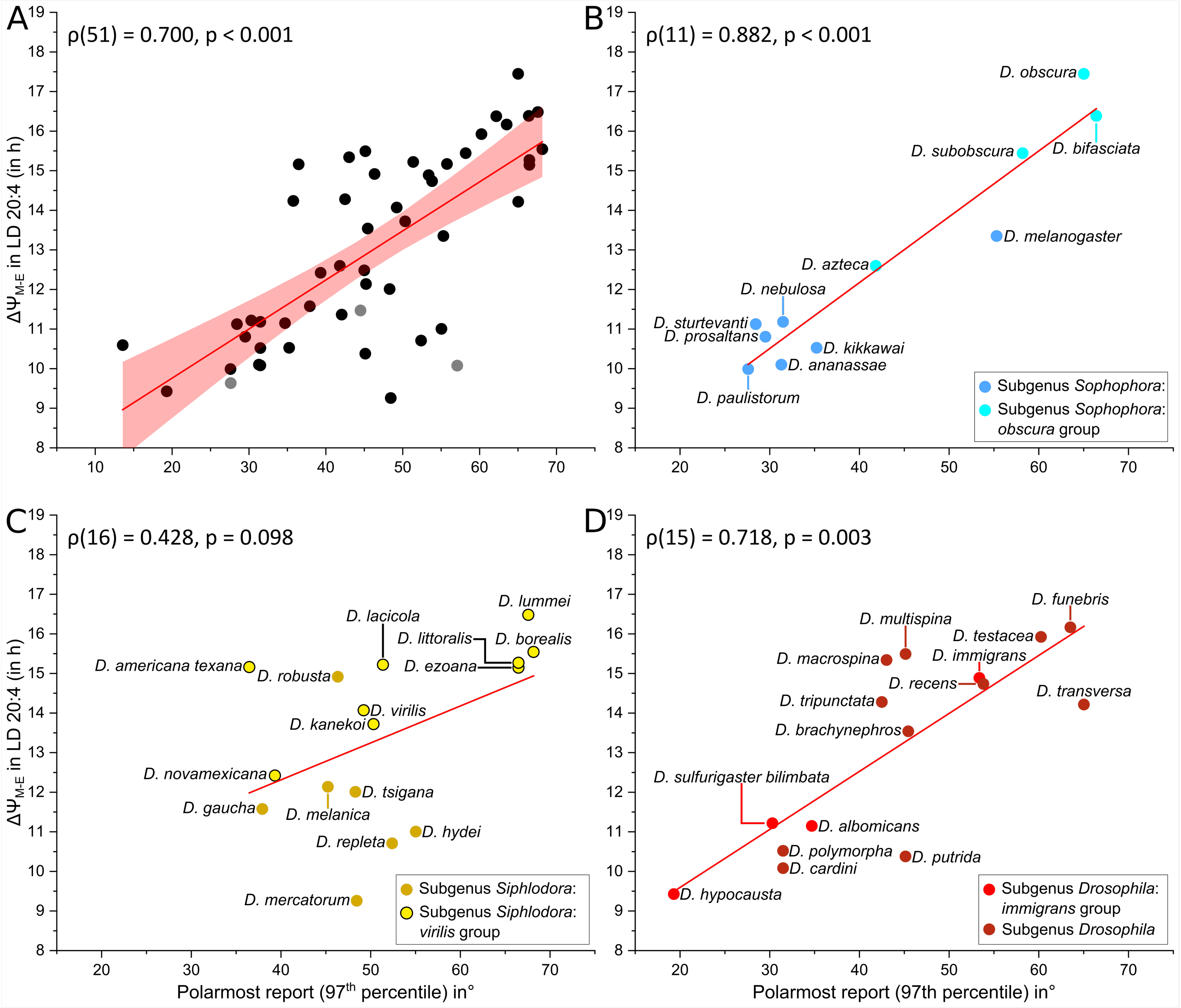

**Figure 6—figure supplement 1.** Correlation between $\Delta\Psi_{M-E}$ in LD 20:4 and most poleward report of all *Drosophila* species **(A)** and the three different *Drosophila* subgenera **(B-D)**. To avoid distorting the correlation coefficient with duplicate values for species with multiple tested strains, only one strain per species is included (51 species total): *D. subobscura* from Bavaria, *S. pallida* collected in 2020, *D. repleta* from Bavaria (strain S), *D. virilis* from Rwanda, *D. funebris* from Alaska, and *D. macrospina* from Arizona. The three *Scaptodrosophila* species, which fall outside the main *Drosophila* radiation (see Figure 2), are shown in grey. **(B)** *Sophophora* subgenus, **(C)** *Drosophila* subgenus, **(D)** *Siphlodora* subgenus; Regression lines are indicated in red. The 95 % confidence band is illustrated in **(A)**. Spearman's rank correlation coefficients are indicated in the upper left corner of each panel.

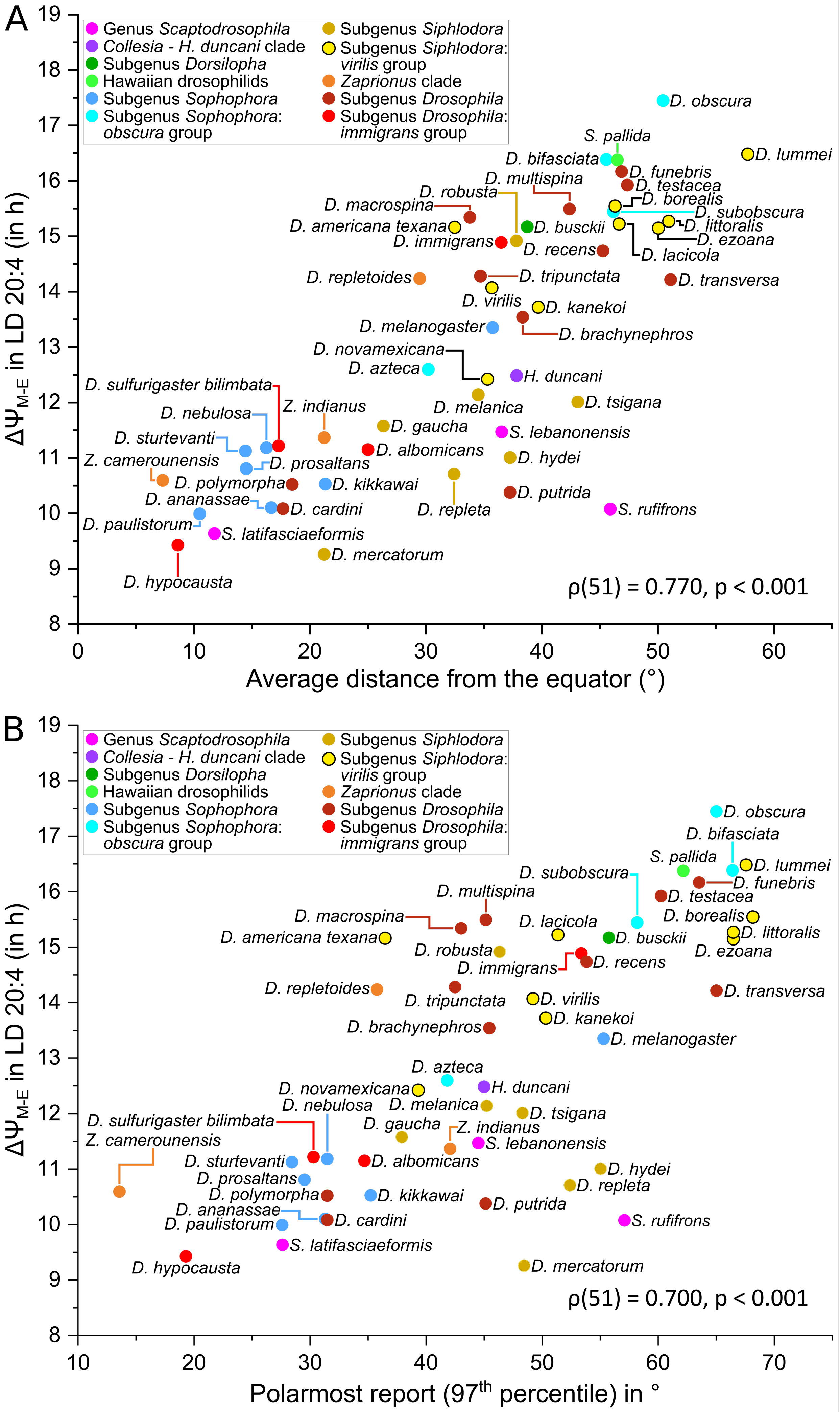
A
Genus Scaptodrosophila
Collesia - H. duncani clade
Subgenus Dorsilopha
Hawaiian drosophilids
Subgenus Sophophora
Subgenus Sophophora: obscura group
Subgenus Siphlodora
Subgenus Siphlodora: virilis group
Zaprionus clade
Subgenus Drosophila
Subgenus Drosophila: immigrans group
ρ(51) = 0.770, p < 0.001
ΔΨM-E in LD 20:4 (in h)
Average distance from the equator (°)
B
ρ(51) = 0.700, p < 0.001
Polarmost report (97th percentile) in °

**Figure 6—figure supplement 2.** Correlation between $\Delta\Psi_{M-E}$ in LD 20:4 and latitudinal distribution of all *Drosophila* species. **A**: Correlation with the average distance from the equator (trimmed by 3 % to exclude potential outlier reports, i.e., false extreme latitudinal values). **B**: Correlation with polarmost report.

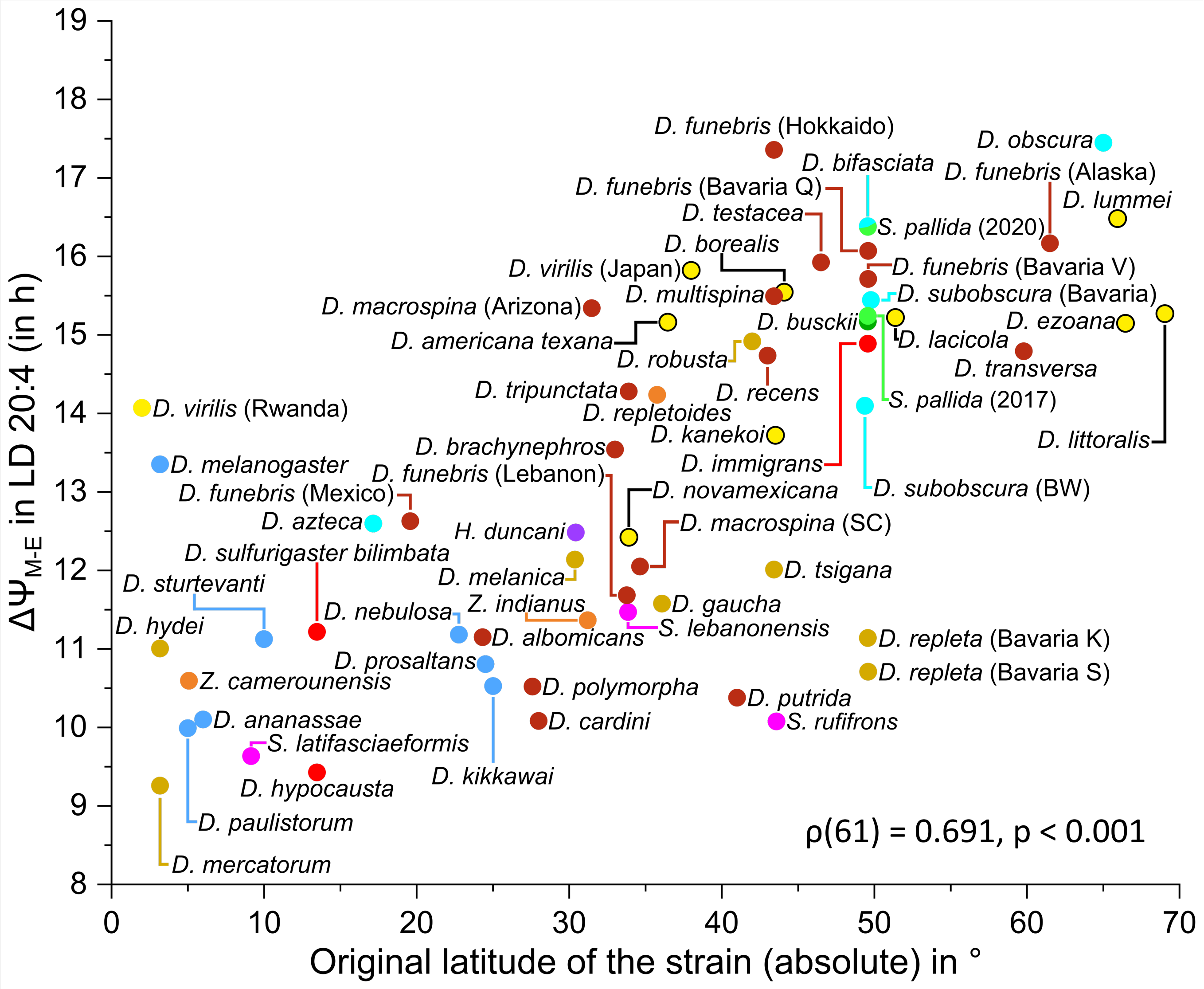

**Figure 6—figure supplement 3.** Correlation between $\Delta\Psi_{M-E}$ in LD 20:4 and the original latitude of all tested strains (four strains of *D. funebris*, and two strains of *S. pallida*, *D. virilis*, *D. repleta* and *D. macrospina,* respectively). The four *D. funebris* strains originated from very different latitudes and those stemming from higher latitudes showed a larger $\Delta\Psi_{M-E}$ than those from lower latitudes, suggesting that $\Delta\Psi_{M-E}$ increases with latitude even within a single species.

### Figure 7—figure supplements

➔ See next pages

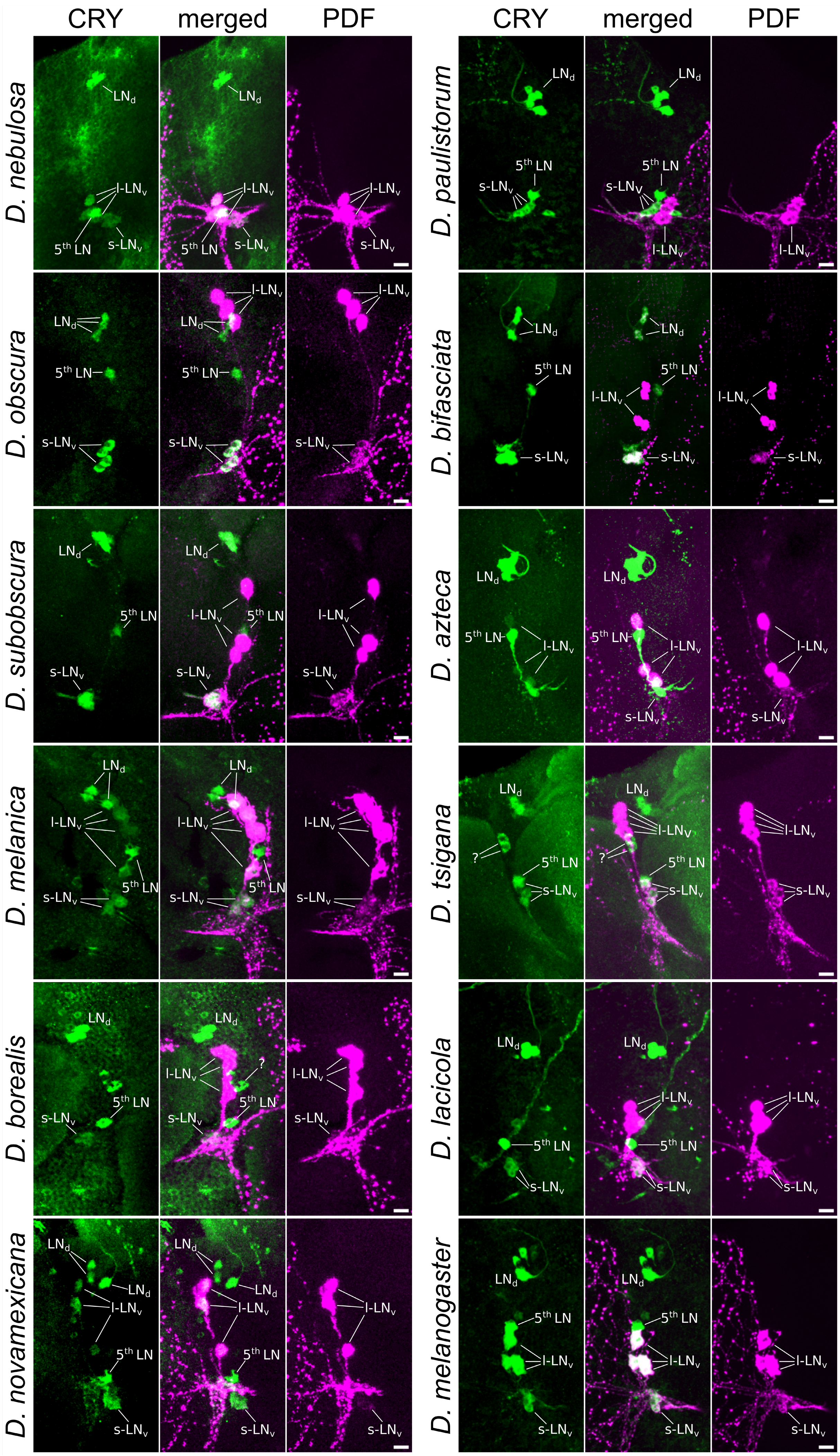
CRY
merged
PDF
D. nebulosa
D. obscura
D. subobscura
D. melanica
D. borealis
D. novamexicana
D. paulistorum
D. bifasciata
D. azteca
D. tsigana
D. lacicola
D. melanogaster
LN$_d$
l-LN$_v$
s-LN$_v$
5$^{th}$ LN
?

**Figure 7—figure supplement 1.** Expression patterns of CRY and PDF in the lateral clock neuron clusters of different species. The focus in this figure is on species with an expression pattern differing from that of *D. melanogaster* (row 6, lower right). Shown are maximum intensity projections of confocal images of the lateral clock neuron clusters, immunostained with anti-CRY (green) and anti-PDF (magenta); middle panels show an overlay of both channels. The dorsolateral neurons (LN$_d$) and the fifth lateral neuron (5$^{th}$ LN) belong to the evening neurons, whereas the small ventrolateral neurons (s-LN$_v$) are morning neurons. The top row shows two species of the *willistoni* group (subgenus Sophophora): *D. nebulosa* has CRY in the l-LNv, whereas its close relative *D. paulistorum* lacks it in this cluster. While *D. nebulosa* features CRY in the l-LN$_v$ neurons, its close relative *D. paulistorum* lacks it in this cluster. Rows 3 and 4 show all investigated species of the *obscura* group; the three northern species *D. obscura*, *D. bifasciata*, and *D. subobscura* lack CRY in all l-LNv, whereas it is weakly expressed in the more southern *D. azteca*. The *melanica* group species differ in the l-LN$_v$ expression of CRY, which is weakly present in *D. melanica* but absent in *D. tsigana* (row 4). Rows 5 and 6 show three newly described virilis group species; all share weak PDF expression in the s-LNv, but in contrast to the boreal *D. borealis* and *D. lacicola*, the arid-adapted *D. novamexicana* from the south-western United States showed CRY-positive signals in the l-LNv cluster. Scale bar: 10 µm.

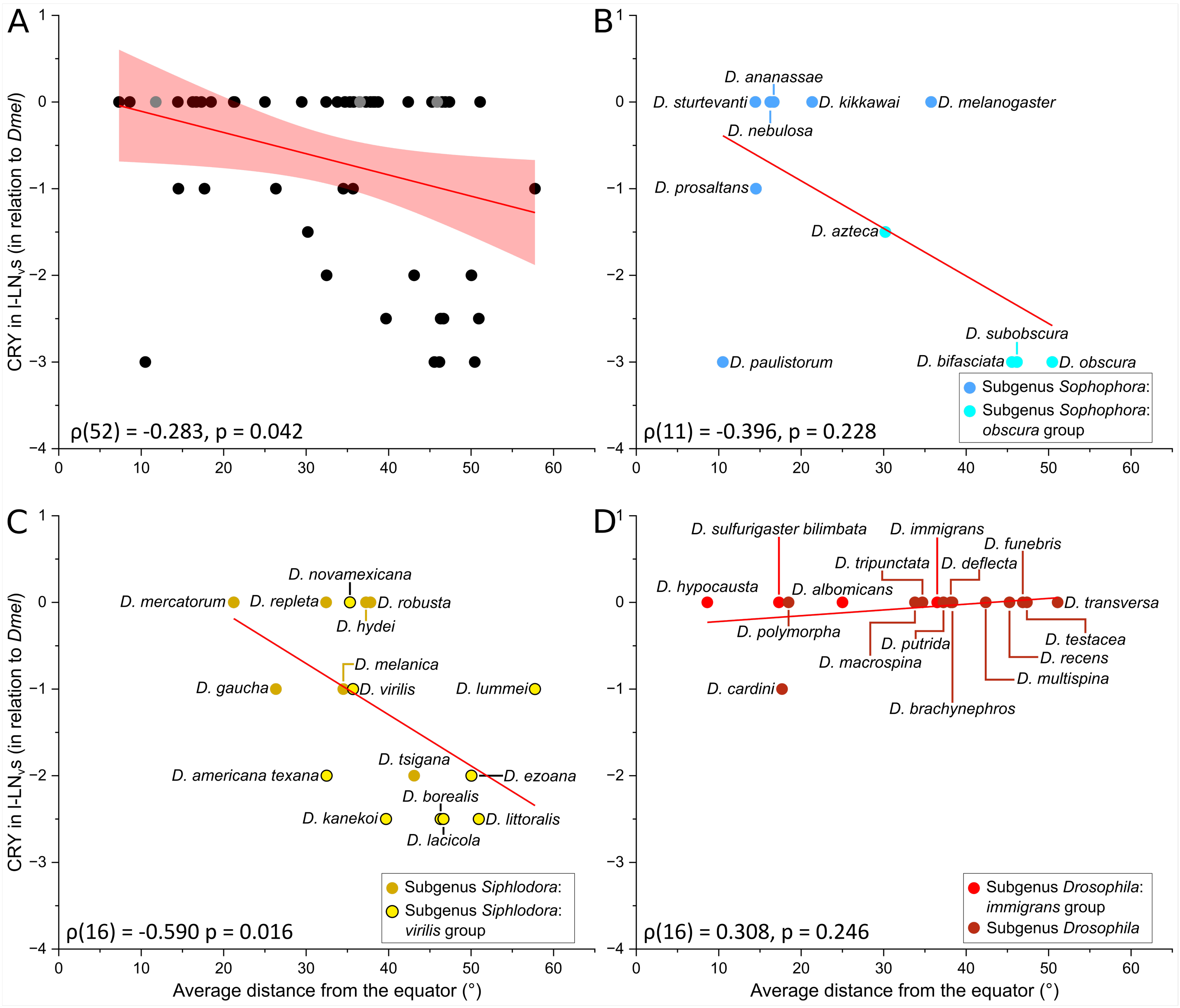

**Figure 7—figure supplement 2.** Correlation between CRY expression in the l-LNv cluster and latitude (average distance from the equator, trimmed by 3 % to exclude potential outlier records, i.e., extreme latitudinal values), shown across all *Drosophila* species **(A)** and within the three main subgenera **(B–D)**. The three *Scaptodrosophila* species in (A), which fall outside the main *Drosophila* radiation (see Figure 2), are shown in grey.

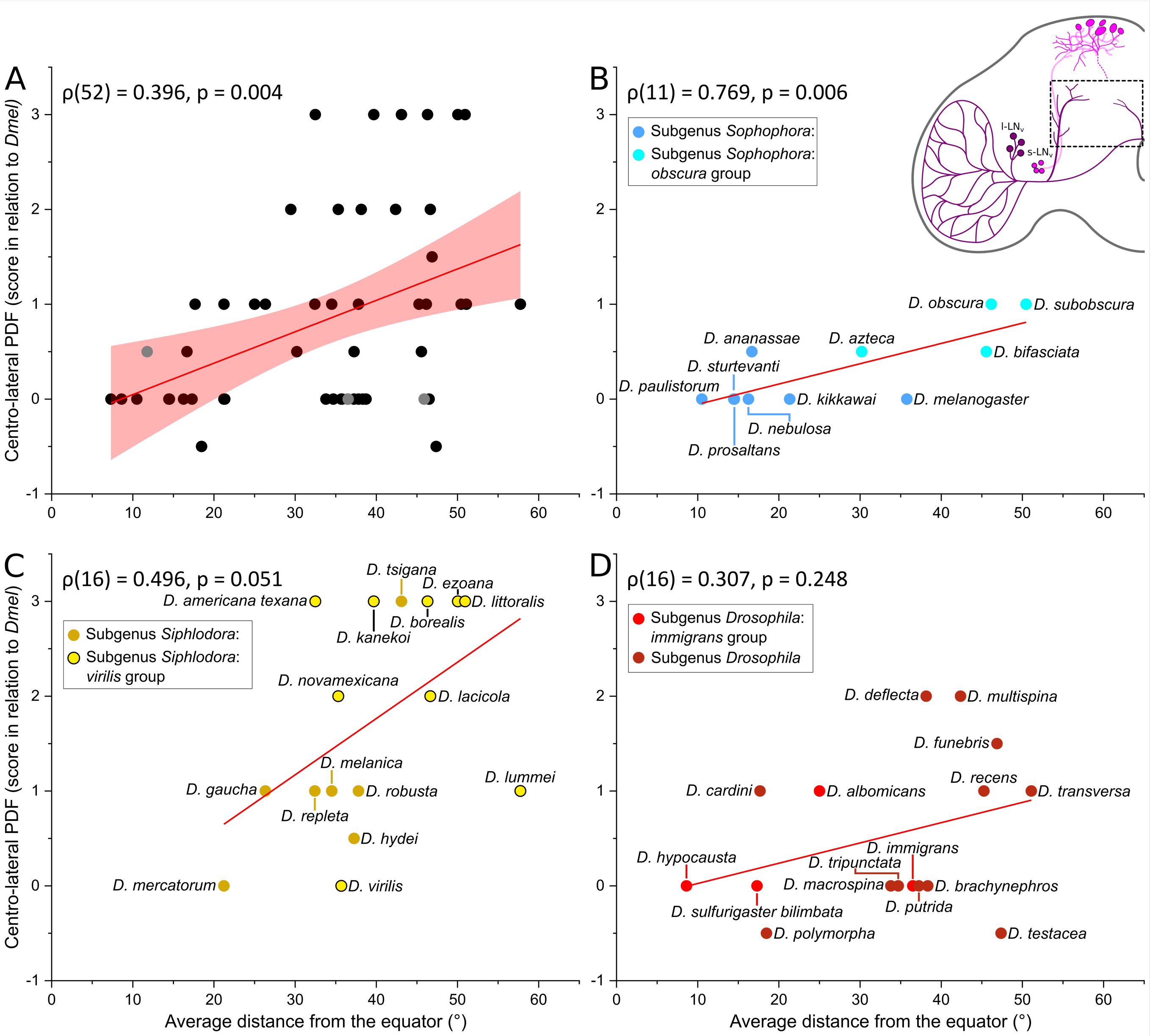

**Figure 9—figure supplement 1.** Correlations between the relative abundance of PDF in the centrolateral brain area and latitude in all *Drosophila* species **(A)** and within the three different *Drosophila* subgenera **(B-D)**. The inset displayed in the upper right indicates the quantified brain region. **(B)** *Sophophora* subgenus, **(C)** *Drosophila* subgenus, **(D)** *Siphlodora* subgenus. The average latitudinal distance from the equator was calculated from a 3 %-trimmed dataset to exclude potential outlier reports, i.e., false extreme latitudinal values.

**Figure 10—figure supplement**

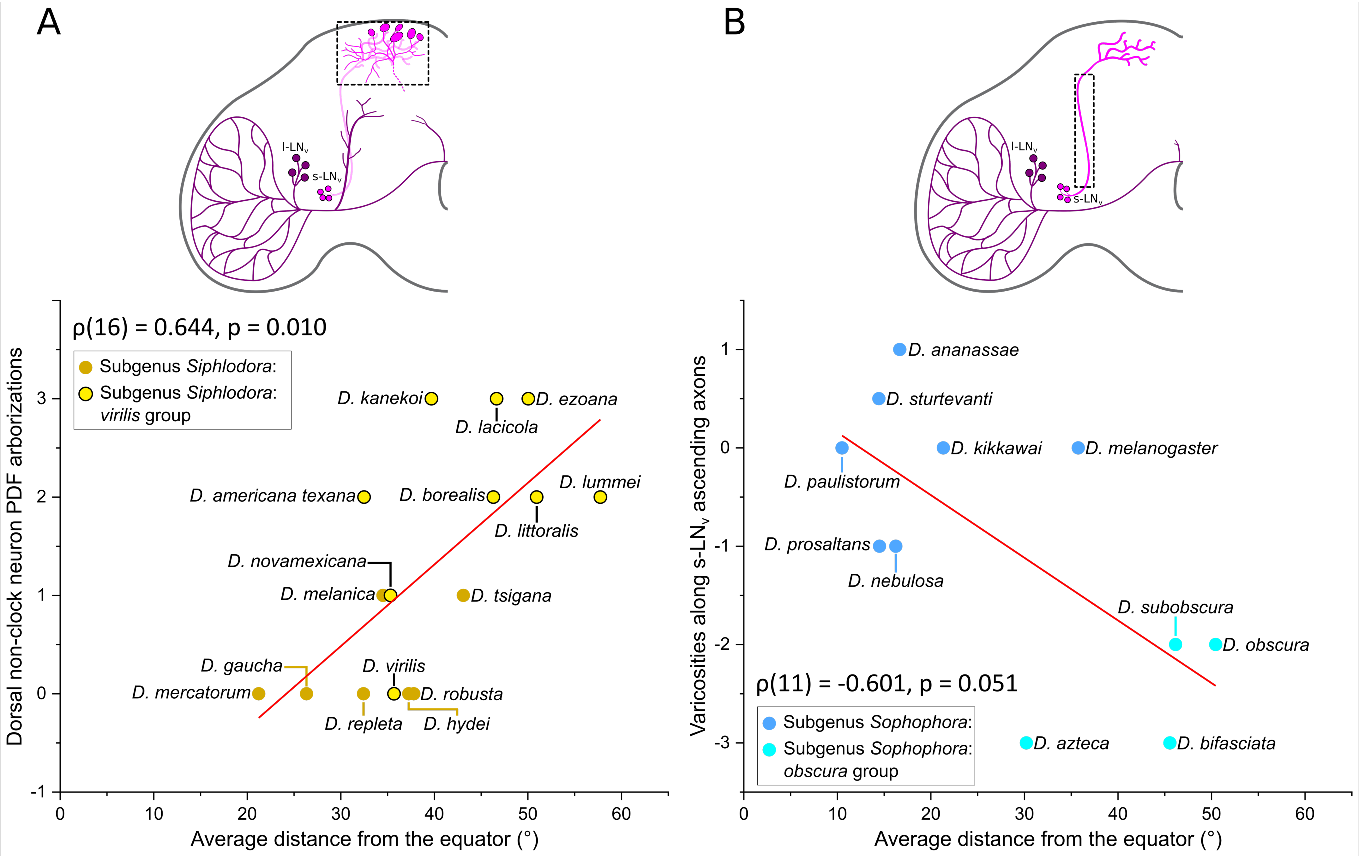

**Figure 10—figure supplement 1.** Correlations between the relative abundance of PDF in the dorsal protocerebrum and latitude (average distance from the equator, trimmed by 3 % to exclude potential outlier reports, i.e., false extreme latitudinal values) within the *Siphlodora* subgenus **(A)**, and between PDF abundance in the ascending s-$LN_v$ axons and latitude within the *Sophophora* subgenus **(B).**

**Figure 11—figure supplement**

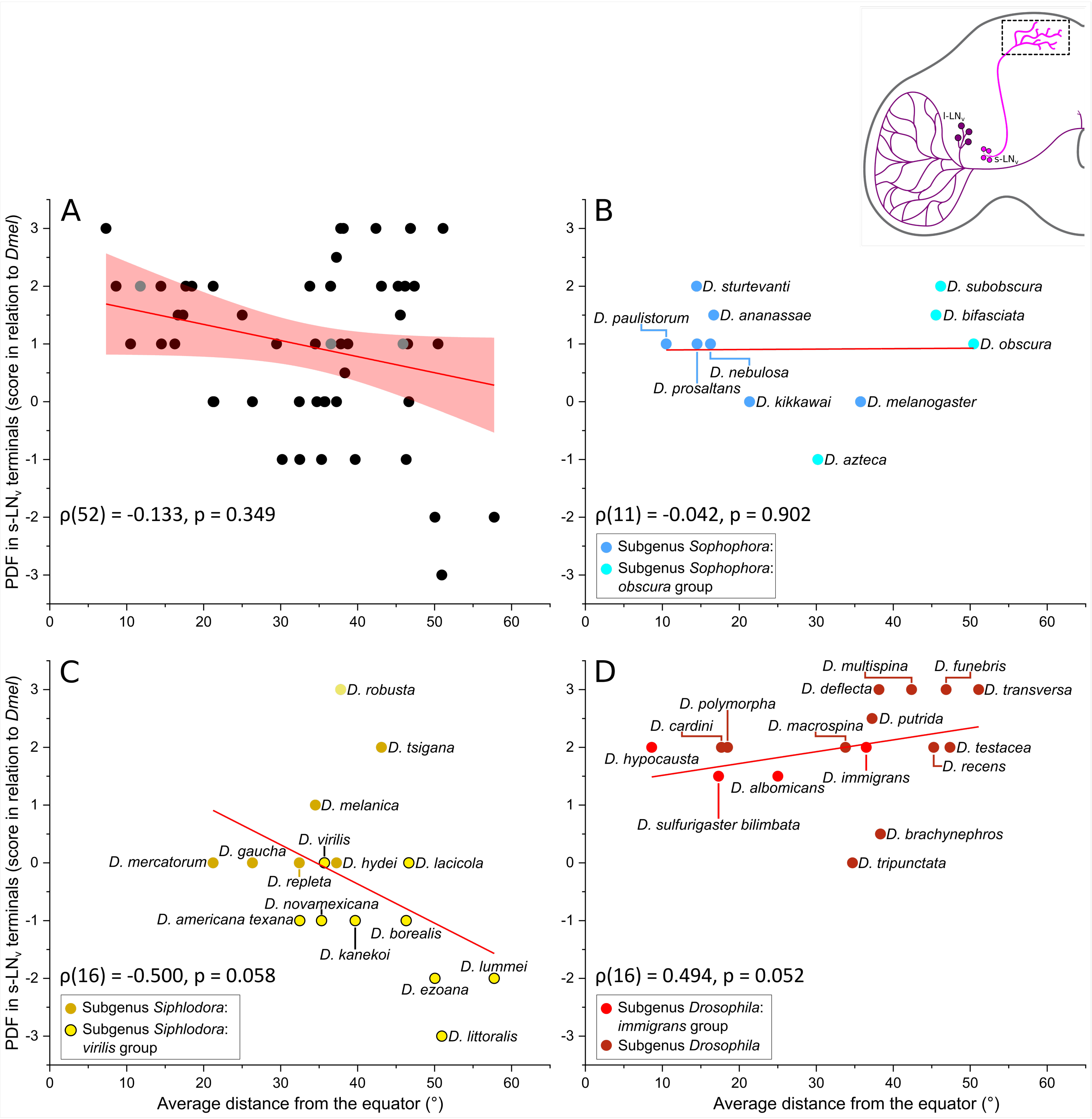

**Figure 11—figure supplement 1.** Correlation between PDF in the s-LNv terminals and latitude (average distance from the equator, trimmed by 3 % to exclude potential outlier reports, i.e., false extreme latitudinal values), shown across all *Drosophila* species **(A)** and within the three main subgenera **(B–D)**. The inset displayed in the upper right indicates the quantified brain region.

**Figure 13—figure supplements**

➔ see next pages

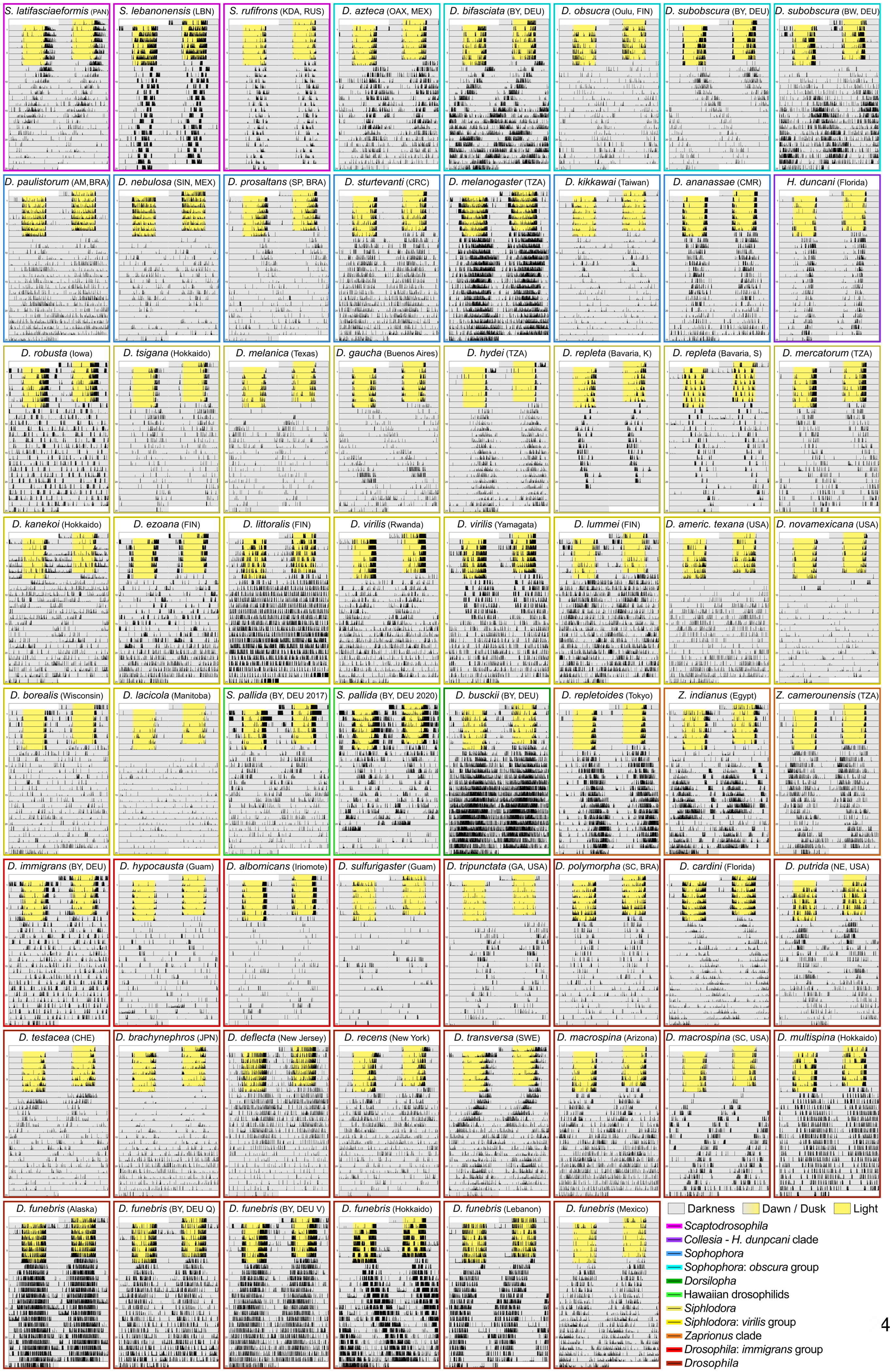
S. latifasciaeformis (PAN)
S. lebanonensis (LBN)
S. rufifrons (KDA, RUS)
D. azteca (OAX, MEX)
D. bifasciata (BY, DEU)
D. obsucra (Oulu, FIN)
D. subobscura (BY, DEU)
D. subobscura (BW, DEU)
D. paulistorum (AM,BRA)
D. nebulosa (SIN, MEX)
D. prosaltans (SP, BRA)
D. sturtevanti (CRC)
D. melanogaster (TZA)
D. kikkawai (Taiwan)
D. ananassae (CMR)
H. duncani (Florida)
D. robusta (Iowa)
D. tsigana (Hokkaido)
D. melanica (Texas)
D. gaucha (Buenos Aires)
D. hydei (TZA)
D. repleta (Bavaria, K)
D. repleta (Bavaria, S)
D. mercatorum (TZA)
D. kanekoi (Hokkaido)
D. ezoana (FIN)
D. littoralis (FIN)
D. virilis (Rwanda)
D. virilis (Yamagata)
D. lummei (FIN)
D. americ. texana (USA)
D. novamexicana (USA)
D. borealis (Wisconsin)
D. lacicola (Manitoba)
S. pallida (BY, DEU 2017)
S. pallida (BY, DEU 2020)
D. busckii (BY, DEU)
D. repletoides (Tokyo)
Z. indianus (Egypt)
Z. camerounensis (TZA)
D. immigrans (BY, DEU)
D. hypocausta (Guam)
D. albomicans (Iriomote)
D. sulfurigaster (Guam)
D. tripunctata (GA, USA)
D. polymorpha (SC, BRA)
D. cardini (Florida)
D. putrida (NE, USA)
D. testacea (CHE)
D. brachynephros (JPN)
D. deflecta (New Jersey)
D. recens (New York)
D. transversa (SWE)
D. macrospina (Arizona)
D. macrospina (SC, USA)
D. multispina (Hokkaido)
D. funebris (Alaska)
D. funebris (BY, DEU Q)
D. funebris (BY, DEU V)
D. funebris (Hokkaido)
D. funebris (Lebanon)
D. funebris (Mexico)
Darkness
Dawn / Dusk
Light
Scaptodrosophila
Collesia - H. dunpcani clade
Sophophora
Sophophora: obscura group
Dorsilopha
Hawaiian drosophilids
Siphlodora
Siphlodora: virilis group
Zaprionus clade
Drosophila: immigrans group
Drosophila

4

**Figure 13—figure supplement 1**. Representative actograms of behavior in constant darkness (DD) for each species and strain. After seven days of entrainment in LD 12:12, flies were recorded for 18 days in DD. Shown are double-plotted actograms. Colored frames indicate the phylogenetic position of each species; geographic origin of the strains is given in parentheses, using ISO 3166-1 and ISO 3166-2 codes for country and region/state abbreviations, respectively. Actograms were exported from ActogramJ with an upper threshold limit of 1.0 and a zoom value of 6.0 (each single bar in the depiction represents the average of 6-minute bins).

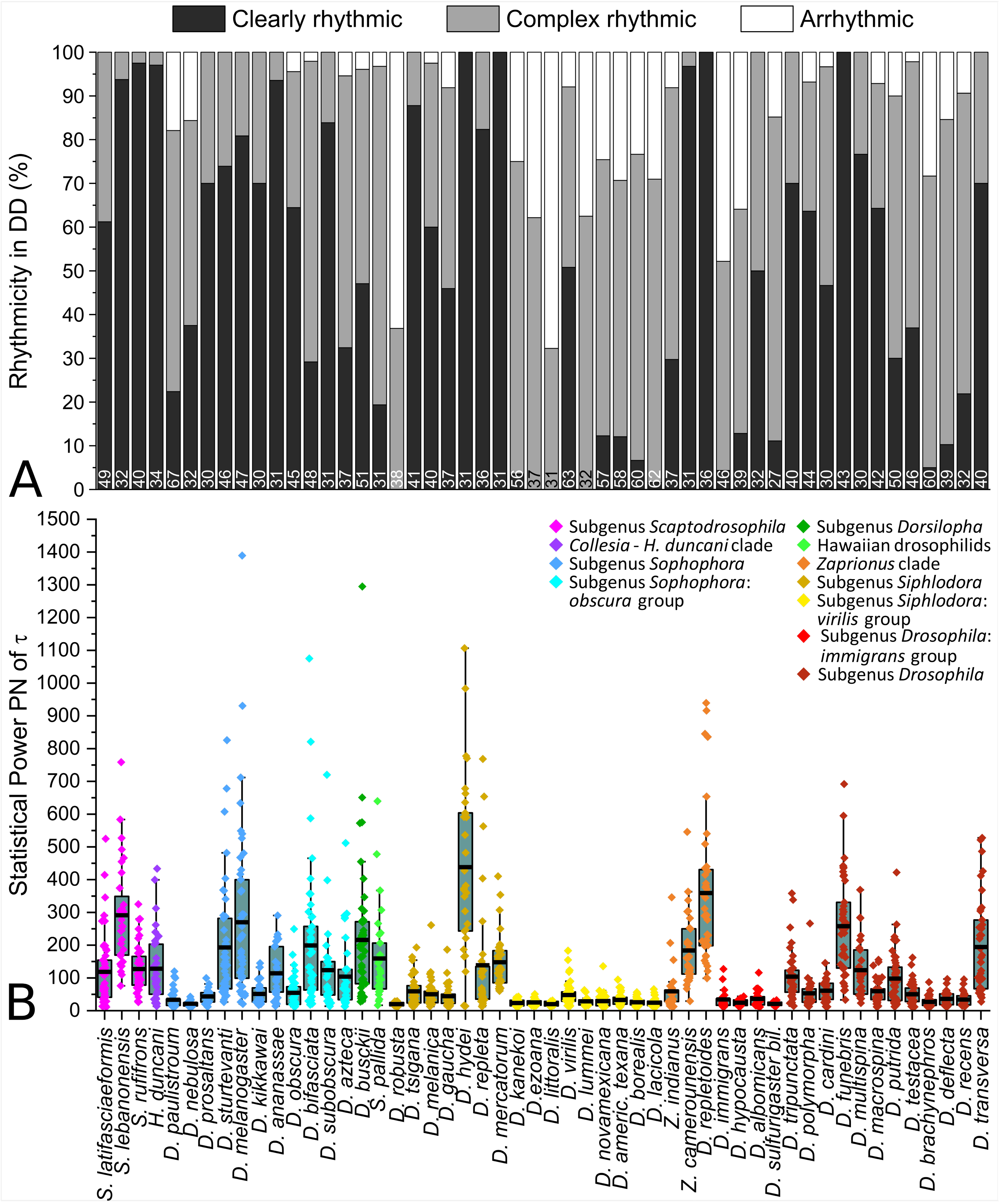

**Figure 13—figure supplement 2**. Statistics of DD behavior across species. The same strains used for the correlation analyses are displayed here. **(A)** Rhythmicity scores for each strain. Every single fly was analyzed by Lomb-Scargle (LS) periodogram analysis, cross-checked against its actogram, and eventually categorized as clearly rhythmic, complex rhythmic, or arrhythmic. The total number of flies analyzed is given at the bottom of each column. **(B)** Distribution of periodogram power values at the free-running period τ. For strains with two free-running components, only the major component (i.e., the one with the stronger power) was considered. Black horizontal lines indicate the arithmetic mean; boxes span the 25th to 75th percentiles. For species with multiple strains, only the strain also used in the correlation analysis is displayed here.

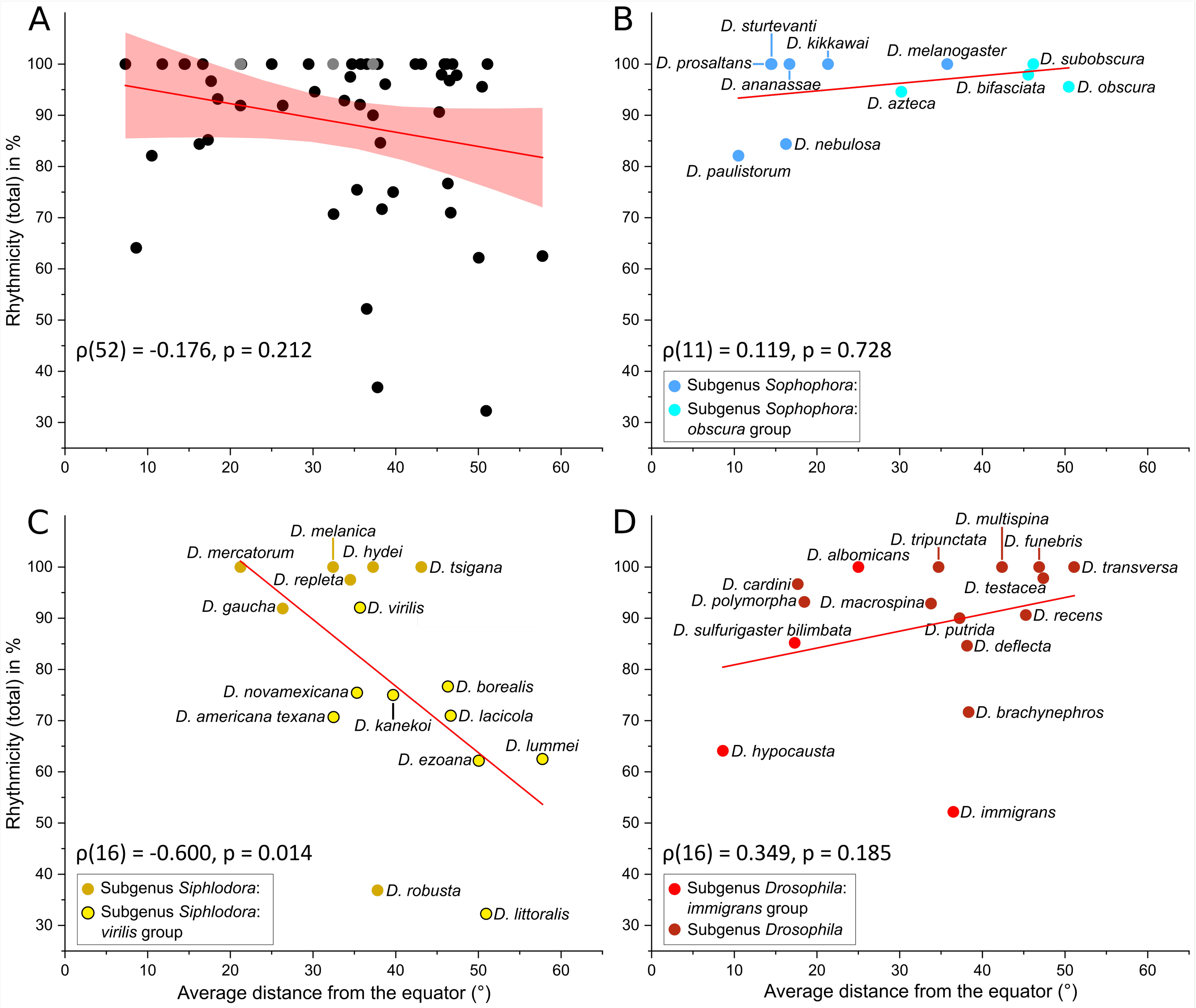

**Figure 13—figure supplement 3.** Percentage of flies with robust rhythms as a function of latitudinal distribution (mean trimmed by 3 % to exclude potential outlier reports, i.e., false extreme latitudinal values). Spearman's rank correlation coefficient (ρ) is given in each panels. **(A)** Geographic range does not influence rhythm power across Drosophilidae as a whole. **(B)** and **(D)** No significant correlations between species range and rhythm strength was found in the subgenera *Sophophora* and *Drosophila*. **(C)** Within the *Siphlodora* subgenus, there was a significant negative relationship between latitudinal distribution and robust rhythmicity.

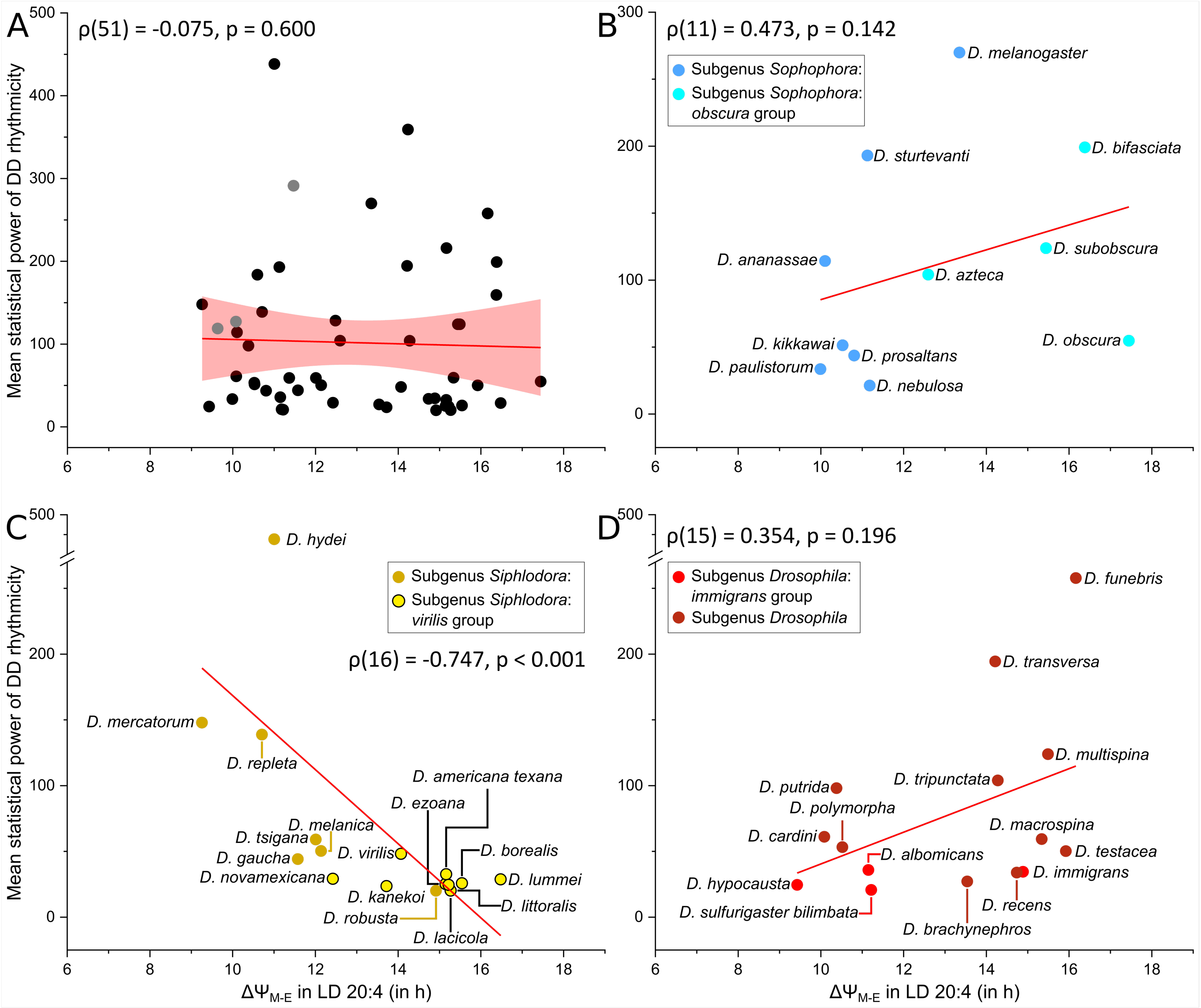

**Figure 13—figure supplement 4.** Correlation between the power of rhythmicity and $\Delta\Psi_{M\text{-}E}$ of the different species. Spearman's rank correlation coefficient (ρ) is given in each panels. Only in the *Siphlodora* subgenus, a lower power of rhythmicity correlates with a larger $\Delta\Psi_{M\text{-}E}$.

Figure 14—figure supplement

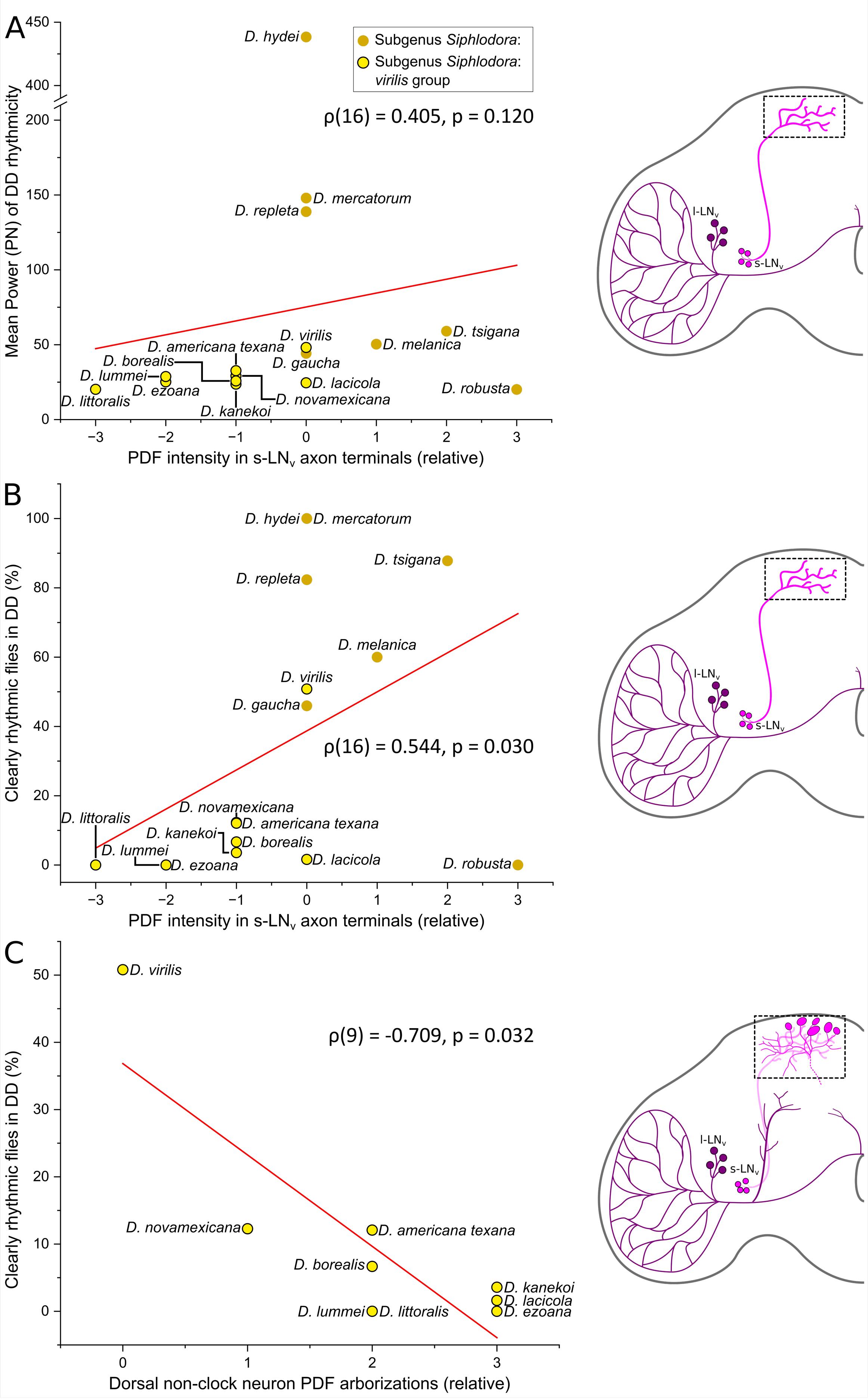

**Figure 14—figure supplement 1.** PDF staining in the s-LN$_v$ terminals in relation to rhythm power or clearly rhythmic flies in the S*iphlodora* subgenus and to PDF in the CRZ neurons in the *virlilis* group of *Siphlodora* flies.
